# Population-scale skeletal muscle single-nucleus multi-omic profiling reveals extensive context specific genetic regulation

**DOI:** 10.1101/2023.12.15.571696

**Authors:** Arushi Varshney, Nandini Manickam, Peter Orchard, Adelaide Tovar, Christa Ventresca, Zhenhao Zhang, Fan Feng, Joseph Mears, Michael R Erdos, Narisu Narisu, Kirsten Nishino, Vivek Rai, Heather M Stringham, Anne U Jackson, Tricia Tamsen, Chao Gao, Mao Yang, Olivia I Koues, Joshua D Welch, Charles F Burant, L Keoki Williams, Chris Jenkinson, Ralph A DeFronzo, Luke Norton, Jouko Saramies, Timo A Lakka, Markku Laakso, Jaakko Tuomilehto, Karen L Mohlke, Jacob O Kitzman, Heikki A Koistinen, Jie Liu, Michael Boehnke, Francis S Collins, Laura J Scott, Stephen C J Parker

## Abstract

Skeletal muscle, the largest human organ by weight, is relevant in several polygenic metabolic traits and diseases including type 2 diabetes (T2D). Identifying genetic mechanisms underlying these traits requires pinpointing cell types, regulatory elements, target genes, and causal variants. Here, we use genetic multiplexing to generate population-scale single nucleus (sn) chromatin accessibility (snATAC-seq) and transcriptome (snRNA-seq) maps across 287 frozen human skeletal muscle biopsies representing nearly half a million nuclei. We identify 13 cell types and integrate genetic variation to discover *>*7,000 expression quantitative trait loci (eQTL) and *>*100,000 chromatin accessibility QTLs (caQTL) across cell types. Learning patterns of e/caQTL sharing across cell types increased precision of effect estimates. We identify high-resolution cell-states and context-specific e/caQTL with significant genotype by context interaction. We identify nearly 2,000 eGenes colocalized with caQTL and construct causal directional maps for chromatin accessibility and gene expression. Almost 3,500 genome-wide association study (GWAS) signals across 38 relevant traits colocalize with sn-e/caQTL, most in a cell-specific manner. These signals typically colocalize with caQTL and not eQTL, highlighting the importance of population-scale chromatin profiling for GWAS functional studies. Finally, our GWAS-caQTL colocalization data reveal distinct cell-specific regulatory paradigms. Our results illuminate the genetic regulatory architecture of human skeletal muscle at high resolution epigenomic, transcriptomic, and cell-state scales and serve as a template for population-scale multi-omic mapping in complex tissues and traits.

## 1 Introduction

Skeletal muscle, the largest organ in the adult human body by mass (*>*40%)^1^, facilitates mobility, sustaining life functions, and influences quality of life. Beyond its mechanical functions, skeletal muscle plays a central role in metabolic processes, particularly in glucose uptake and insulin resistance^1–5^. Metabolic diseases and traits, such as type 2 diabetes (T2D), fasting insulin, waist-to-hip ratio (WHR), and others are complex and polygenic, involving a multitude of genetic factors. Genome-wide association studies (GWAS) have identified thousands of genetic signals associated with these diseases and traits^6–11^. However, ∼90% of these variants lie within non-coding regions^12^, are enriched to overlap tissue-specific enhancers, and are therefore expected to regulate gene expression^8,13–15^. Additionally, GWAS loci are often tagged by numerous variants in high linkage disequilibrium (LD), and can harbor multiple causal variants^16^. For these reasons, identifying the biological mechanisms and pinpointing causal variants in GWAS loci remains challenging.

Information encoded in DNA, which is largely invariant across cells in the body, likely percolates through several molecular layers to influence disease. The mostly non-coding genetic variation identified through GWAS likely has the most proximal effect on the molecules bound to DNA (epigenome), which in turn can influence the expression of target genes (transcriptome), and then levels of proteins, all of which can vary by the cell type^17^. This molecular cascade is not completely unidirectional and it is dynamic in nature. For example, changes in expression of a transcription factor (TF) can feed back to changes in the epigenome. The epigenome and the transcriptome layers are therefore valuable to gain insights about gene regulation. One approach to link these layers with GWAS is through identification of quantitative trait loci (QTL) for epigenomic modalities such as chromatin accessibility QTL (caQTL) and gene expression quantitative trait loci (eQTL) followed by testing whether common causal variants underlie the molecular QTL and GWAS signals (i.e. if the signals are formally colocalized)^16,18–28^.

Previous studies profiling the epigenome and transcriptome in bulk skeletal muscle across hundreds of samples identified expression and DNA methylation QTLs and provided valuable insights^29–31^. However, bulk skeletal muscle profiles are dominated by the most prominent muscle fiber types, and other less abundant but relevant cell types are largely missed. Several resident cell types are essential for muscle function^3^. For example, muscle fibro-adipogenic progenitors (FAPs) are resident interstitial stem cells involved in muscle homeostasis and along with muscle satellite cells, regulate muscle regeneration^32–35^. Diabetes and obesity not only lead to structural and metabolic changes of the muscle fibers but also exert detrimental effects on these progenitor cells^36–38^. Endothelial cells and smooth muscle cells comprise the muscle vasculature which is another important component in diabetes-associated complications, involving insulin uptake^39^. Immune cells are also critical, especially following injury^40^. Recent studies have generated reference epigenome and transcriptome maps in human skeletal muscle at a single-nucleus/single-cell resolution^41–44^. However, population-scale studies are imperative to identify e/caQTL within each cell type to enable exhaustive interrogation of mechanistic signatures underlying GWAS signals. To date, there is no single-nucleus/cell resolution population-scale study that maps e/caQTL in hundreds of samples.

We hypothesize that single-nucleus epigenome (snATAC-seq) and transcriptome (snRNA-seq) profiling across hundreds of genotyped samples will help identify the appropriate cell type, regulatory elements, target genes, and causal variants(s) in elucidating context-specific regulatory mechanisms within skeletal muscle. In this work, we perform snRNA-seq and snATAC-seq across skeletal muscle samples from 287 Finnish individuals^29^. We integrate these molecular profiles with genetic variation to identify cell-specific eQTL and caQTL. We further integrate the e/caQTL signals with GWAS by testing for colocalization and infer the chain of causality between these modalities using mediation analyses, and highlight our findings with orthogonal methods at multiple example loci.

## 2 Results

### 2.1 snRNA and snATAC profiling and integration identifies 13 distinct cell type clusters

We generated a rich dataset of snRNA and snATAC across 287 frozen human skeletal muscle (*vastus lateralis*) biopsies from the FUSION study^29^ (**Figure 1A**), as part of a larger study with 408 total samples including three separate smaller cohorts. We processed the samples in ten batches of 40 or 41 samples multiplexed together using a randomized block study design to balance across experimental contrasts of interest (cohort, age, sex, BMI, oral glucose tolerance test (OGTT), **Figures S1A**–**S1E**). We also included multiome data (snRNA and snATAC on the same nucleus) for one muscle sample to help assess our cross-modality clustering. We performed rigorous quality control (QC) of all nuclei and only included those deemed as high-quality (Methods). This led to a total of 188,337 pass-QC RNA nuclei and 268,543 pass-QC ATAC nuclei (**Figures S1F**–**S1J**, **Figures S2A**–**S2D**, **Figures S3A**– **S3E**). As expected, there is a strong correlation across samples for the number of pass-QC RNA and ATAC nuclei (**Figure S3F**), and nuclei counts correlate with the initial weights of the tissue samples (**Figure S3G**), indicating that our genetic demultiplexing and QC recovered high-quality nuclei in expected proportions. Collectively, we generated total N = 625,722 high-quality RNA or ATAC nuclei from all 408 samples, and in this work we analyze N = 456,880 nuclei from the 287 FUSION and one multiome sample.

**Figure 1:**
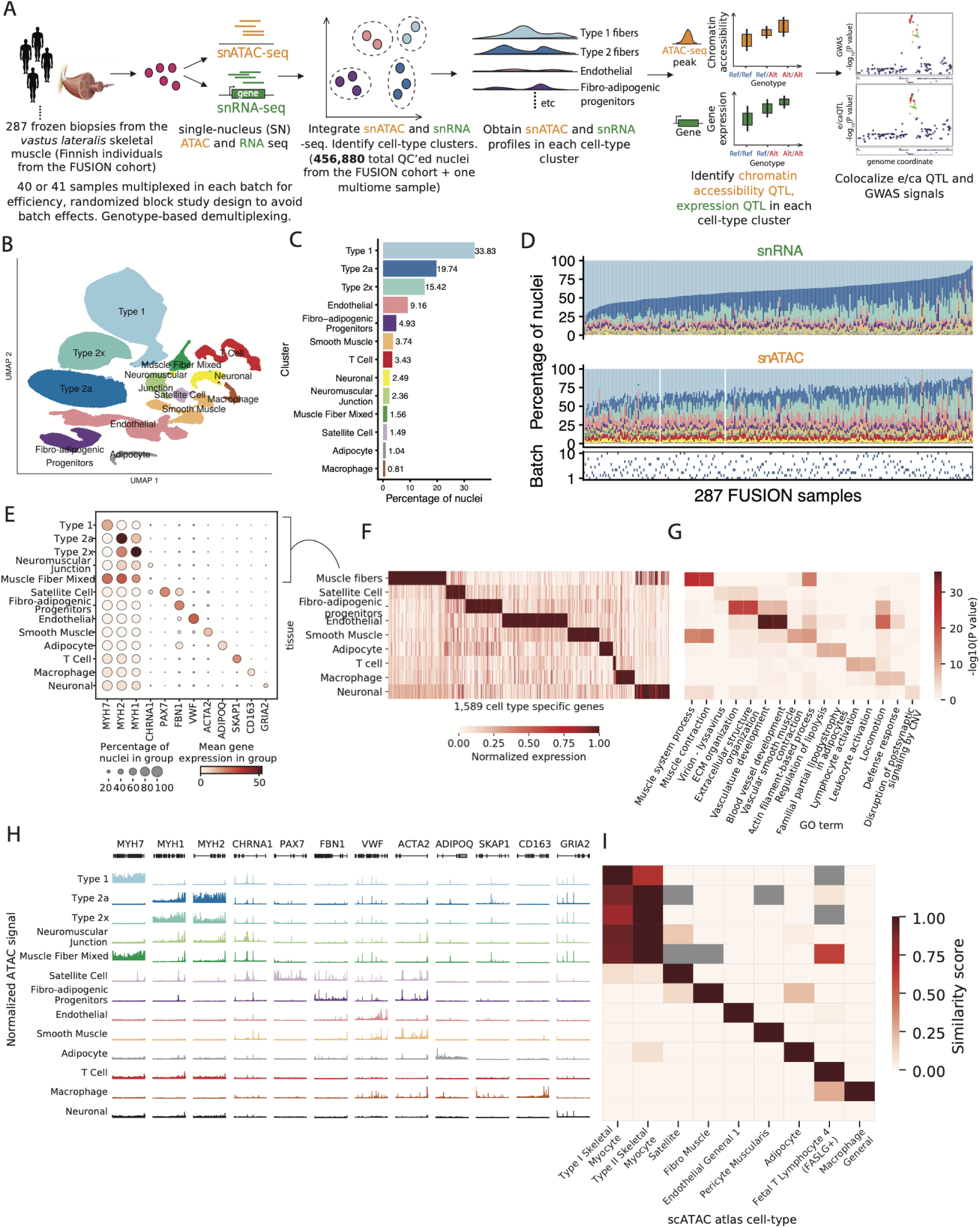
snRNA and snATAC -seq data generation and integration identifies 13 high quality cell-type clusters. (A) Study design including sample processing, snRNA and snATAC -seq profiling, and analyses. (B) UMAP plot showing the 13 identified clusters after jointly clustering the snRNA and snATAC modalities. (C) Cluster abundance shown as percentage of total nuclei. (D) Cluster proportions across samples and modalities. Bottom row denotes the processing batch number (1-10) for samples, indicating that the proportions are not driven by batch effects. (E) Gene expression (post ambient-RNA adjustment) in clusters for known marker genes for various cell-types. (F) Identification of cell-type-specific genes across clusters. Five related muscle fiber clusters (type 1, 2a, 2x, neuromuscular junction and muscle fiber mixed were taken together as a“muscle fiber” cell type). (G) GO term enrichment for cell-type-specific genes identified in (F), showing two GO terms for each cluster. (H) snATAC-seq profiles over known marker genes in clusters. (I) Comparison of snATAC-seq peaks identified for clusters in this study with reference data across various cell-types from the Zhang *et al.* [42] scATAC-seq atlas. Gray cells denote no overlaps between cell-type specific peaks in our dataset and those in the Zhang et al dataset.

We jointly clustered the snRNA and snATAC data, while avoiding batch and modality-specific effects using Liger^45,46^ (**Figure S4A**). We identified 13 distinct clusters representing diverse cell types (**Figure 1B**) that ranged in abundance (**Figure 1C**) from 34% (type 1 fiber) to *<*1% (macrophages). The aggregate cell-specific profiles provide clear evidence of muscle tissue heterogeneity (**Figure 1D**). When treating the multiome RNA and ATAC modalities separate and integrating across them, we found that 82.8% of the non-muscle fiber multiome nuclei had the same RNA and ATAC cluster assignments (**Figure S4B**). This is consistent with previous multiome studies^47,48^ (Supplementary note); for example, integrating 92 brain snATAC+snRNA samples (19 of which were multiome) obtained 79.5%-85% concordant cluster assignments depending on the clustering approach^48^.

The annotated clusters showed expected patterns of expression for known marker genes (**Figure 1E**, **Figure S4C**). We merged the five closely-related muscle fiber types 1, 2a, 2x, mixed and neuromuscular junction (NMJ) together and annotated them as “muscle fiber” and identified 1,569 cellspecific genes using pair-wise differential gene expression analyses (**Figure 1F**). Relevant gene ontology (GO) terms were enriched in these cell-specific genes (**Figure 1G**), for example, muscle system process and muscle contraction terms for muscle fiber and regulation of lipolysis in adipocytes and familial partial lipodystrophy terms for the adipocyte cluster.

The ATAC modality also showed clear patterns of chromatin accessibility over known marker genes for various cell types (**Figure 1H**). We optimized ATAC peak calls to be of similar statistical power, reproducible, and non-redundant across clusters to create a harmonized list of 983,155 consensus peak summits across the 13 cell types (Methods, **Figures S5A**–**S5D**). We compared our snATAC profiles with reference snATAC data from 222 cell types from a previous study^42^. Our snATAC peaks were enriched to overlap peaks identified in related cell types (**Figure 1I**), which reinforces the quality of our cluster labels using the independent ATAC modality. We identified 95,442 snATAC peaks that were specific for a cell type cluster (**Figure S5E**). We computed chromatin co-accessibility between all peak pairs within 1Mb in each cluster using Cicero^49^, which enabled peak to gene TSS links.

DNA-binding motifs for cell type-relevant TFs were enriched in these cluster-specific peaks (**Figure S5F**). For instance, motifs for the myocyte enhancer factor 2 (MEF2) family of TFs that are known regulators of skeletal muscle development and function^50,51^ were enriched for muscle fiber peaks; motifs for the SRY (Sex Determining Region Y)-related HMG box of DNA binding (SOX) TFs, implicated in endothelial differentiation and endothelial-mesenchymal cell transitions^52–54^ were enriched in endothelial-specific peaks. Specifically expressed TF genes appeared to drive corresponding TF motif enrichment in cluster-specific peaks (**Figure S6**). For example, *PAX7* gene, critical for satellite cell function^55^ is expressed with high specificity in muscle satellite cells and PAX7 TF motifs are enriched in satellite cell specific peaks. Other examples included known TF regulators such as SPI1 in macrophages^56^, EB1 in adipocytes^57^, and GATA2 for endothelial^58^ cells. This analysis revealed LHX6 - known for its role in cortical interneuron development^59,60^ - as another key endothelial cell regulator. Collectively, these data demonstrate the high-quality of our snRNA and snATAC profiles and data integration.

### 2.2 Integrating genetic variation with snRNA and snATAC profiles identifies thousands of e/caQTL

We next identified genetic associations with gene expression and chromatin accessibility QTL (e/ca QTL) in clusters. Optimizing QTL discovery (**Figures S7A**–**S7B**, **Figures S8A**–**S8B**), we identified 7,062 eQTL and 106,059 caQTL across clusters (**Figures 2A**–**2B**, **Figure S7C**, **Figure S8C**). 2,452 eQTL (34.7%) and 37,095 caQTL (34.5%) were only detected in one cluster (**Figure S7C**, **Figure S8C**), which is attributable to cell-type specific effects but also differences in power to detect QTL in clusters. Despite differences in power, the e/caQTL effect sizes were highly concordant across clusters (**Figure S7D**,**Figure S8D**). Out of 4,206 unique eGenes identified in our sn-eQTL, 1,014 (24%) were not identified in bulk skeletal muscle eQTL^29^. Notably, out of 2,452 cell-type specific eGenes, 720 (29.4%) were not identified in bulk skeletal muscle eQTL, highlighting the novel findings in our sn-eQTL scans. Down-sampling analyses in type 1 fibers showed an almost linear increase in detectable QTL with the number of samples and number of nuclei, which could be a useful benchmark while designing future studies **Figures S9A**–**S9E**.

**Figure 2:**
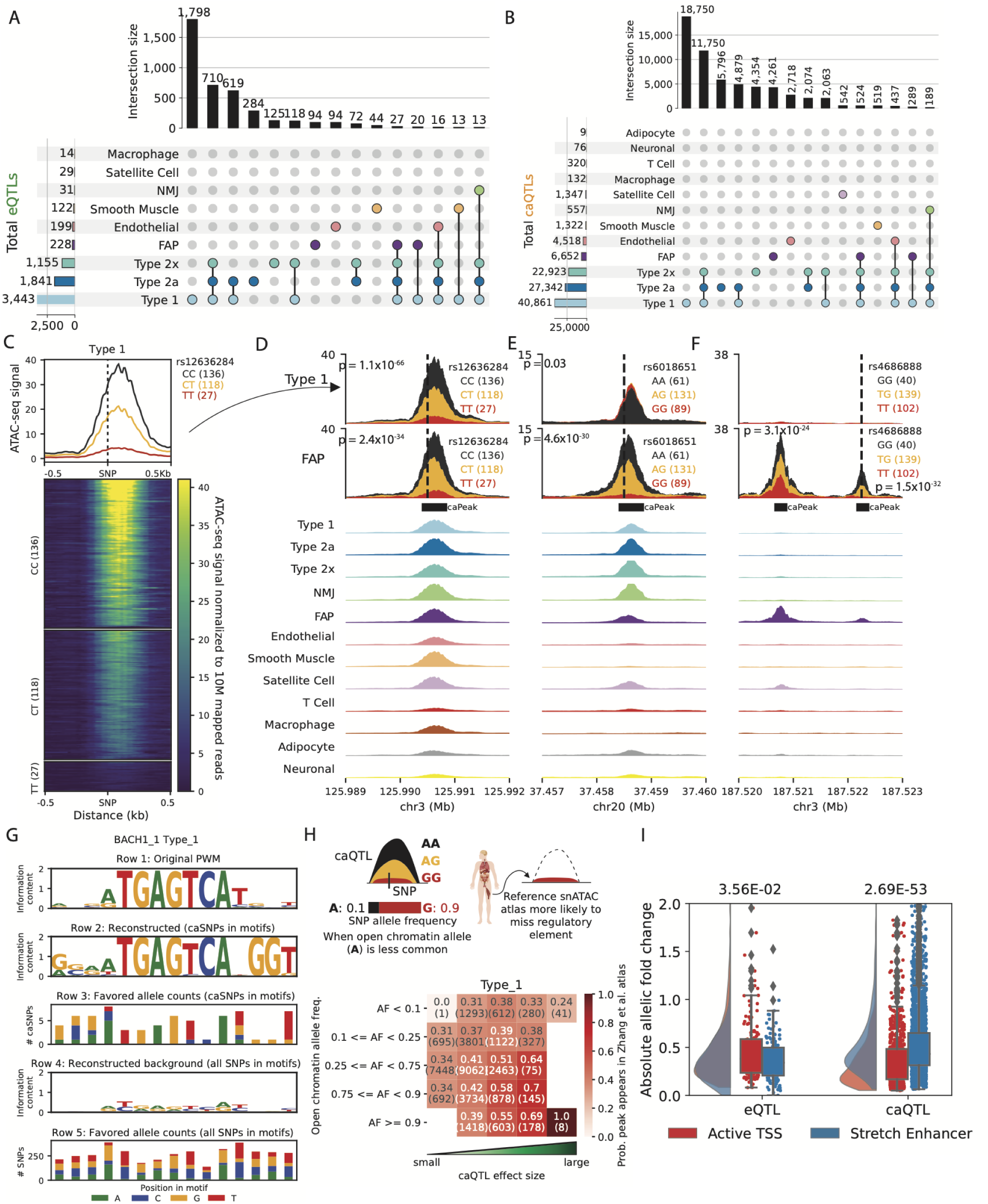
Thousands of e/caQTLs identified in clusters. (A) UpSet plot showing eGenes, and (B) caPeaks in clusters (FDR*<*5%) (C) An example caQTL. Heatmap shows normalized snATAC-seq reads across samples in the type 1 cluster, separated by caSNP rs12336284 genotype classes. Aggregate profiles by genotype are shown on top. Examples of shared and cluster-specific caQTL are shown in (D), (E), and (F). Top two rows show snATAC-seq profiles by the caSNP genotype in type 1 and FAP cell types, followed by aggregate snATAC profiles across clusters. (G) Reconstruction of the BACH 1 TF motif using caQTL data. From top, row 1: original motif PWM. Row 2: genetically reconstructed motif PWM. For all BACH 1 motifs occurring in type 1 snATAC-seq peaks (peak-motifs) that also overlapped type 1 caSNPs, alleles associated with higher chromatin accessibility (“favored alleles”) were quantified using the caQTL aFC, followed by PWM generation. Row 3: favored allele counts for caSNPs in BACH 1 peak-motifs. Row 4: PWM reconstructed using the nucleotide counts for all heterozygous SNPs overlapping the BACH 1 peak-motifs. Row 5: nucleotide counts for all heterozygous SNPs in the BACH 1 peak-motifs. (H) Comparison of caSNP effect size and MAF with the replication of snATAC-seq peaks in a reference scATAC dataset^42^. (I) Allelic fold change for type 1 e/caSNPs that overlap skeletal muscle active TSS or stretch enhancer chromatin states. P values from a two-sided Wilcoxon rank sum test.

**Figure 2C** shows an example type 1 caQTL signal (P = 1.1×10^−66^) where the caQTL SNP (caSNP) rs12636284 lies within the caQTL peak (caPeak), and the C allele is associated with higher chromatin accessibility. This caQTL is also identified in FAPs (P = 2.4×10^−34^), and the peak is shared across multiple clusters (**Figure 2D**). We identified cluster-specific caQTL even for peaks shared across cell types, indicating context-specific genetic effects on chromatin accessibility. For example, **Figure 2E** shows a caQTL identified in FAPs (∼5% ATAC nuclei) and not type 1 fibers (∼30% ATAC nuclei), even when the overall peak was comparable in size between the two clusters (**Figure 2E**, aggregate cluster snATAC tracks). Additionally, we identified cluster-specific peaks as caQTL (**Figure 2F**). caPeaks in clusters were enriched to overlap TF motifs relevant to the corresponding cell type (**Figure S8E**).

We next asked if the genetic regulatory signatures from our caQTL scans recapitulate patterns of TF binding. Most TFs bind accessible chromatin regions by recognizing specific DNA motifs. For genetic variants within bound activator motifs, the allele preferred by the TF should be preferentially associated with higher chromatin accessibility^24^. In **Figure 2G**, we show the known position weight matrix (PWM) for the TF motif BACH 1 (row 1). We considered all BACH 1 motif occurrences across snATAC peaks in type 1 fibers that also overlapped caSNPs, and used the caQTL allelic fold change (aFC) to quantify alleles associated with higher chromatin accessibility (“favored alleles”). We then used these favored alleles to genetically reconstruct the PWM (**Figure 2G**, row 2) (**Figure 2G**, row 3) and found it closely matches the canonical motif PWM (**Figure 2G**, row 1), providing a caQTL-informed *in vivo* verification of the cognate PWM. To further verify that the caQTL-based genetically reconstructed PWM does not simply reflect the allelic composition of SNPs in motifs, we constructed the PWM using the allele count for all heterozygous SNPs observed in the BACH 1 motif occurrences in snATAC peaks (**Figure 2G**, row 4,5). The resulting PWM had low information content and little similarity to the cognate motif (**Figure 2G**, row 4,1). Several other examples of caQTL-informed reconstructions, including for motifs relevant for muscle (MYF6, MYOD1), chromatin architecture (CTCF), and other motifs enriched to occur in type 1 caPeaks (**Figure S8E**) are shown in **Figure S10A**. PWM motifs were highly concordant with caQTL allele preferences. Motifs enriched in caPeaks across cell types had a higher fraction of caQTL alleles consistent with PWM base preferences than the non-enriched motifs (**Figure S10B**). Overall, these results demonstrate how high-quality snATAC and caQTL information can provide base-resolution insights into TF binding and regulation.

Given our deep caQTL results, we next compared caPeaks to snATAC peaks in the same cell types from reference atlas datasets. We reasoned that for caPeaks where the more commonly occurring caSNP allele is associated with lower chromatin accessibility, the caPeak is more likely to be missed in reference datasets that usually only include one or a few representative tissue samples and therefore do not capture population-scale genetic effects. We additionally reasoned that caPeak reproducibility in reference atlases will be lower for large effect-size caSNPs when the allele associated with high chromatin-accessibility occurs rarely in the population. **Figure 2H** delineates this observation comparing type 1 fiber caPeaks with the Zhang *et al.* [42] snATAC atlas type 1 fiber peaks. Even with moderate effect sizes and allele frequencies, the snATAC caPeak was missed in the snATAC atlas about equally as often as it was observed (**Figure 2H**). Overall, this observation underscores the importance of population-scale snATAC studies to exhaustively identify regulatory elements in the human population.

To examine the local chromatin context, we compared chromatin state patterns at e/caQTL in muscle fibers. Type 1 caPeaks were enriched to overlap TSS and enhancer chromHMM states in skeletal muscle (**Figure S8F**). We contrasted two classes of functional regulatory elements, the active TSS chromHMM state that constitutes shared and cell type-specific promoter elements and stretch enhancers that constitute cell identity enhancer elements^13,61,62^. Type 1 fiber eSNPs occurring in the skeletal muscle active TSS chromHMM state had higher eQTL absolute aFC than eSNPs occurring in stretch enhancers (**Figure 2I**, P = 3.56×10^-2^), whereas, type 1 fiber caSNPs occurring in stretch enhancers had higher caQTL absolute aFC than caSNPs in active TSS states (**Figure 2I**, P = 2.69×10^−53^). These results suggest that eQTL scans identify signals largely in proximal gene promoter regions, whereas caQTL scans are able to identify signals in distal and cell-specific regulatory elements, elucidating an important distinction in the two modalities. Collectively, these results reinforce the importance of joint snRNA and snATAC profiling along with e/caQTL analyses to gain mechanistic insights into the genetic regulation of gene expression and distal regulatory element accessibility.

### 2.3 Identifying patterns of shared and cell-type specific e/caQTL signals across clusters

Following our e/caQTL discovery within each cell-type cluster, we sough to learn patterns of shared QTL signals across clusters to increase power and obtain more precise QTL effect estimates. We used multivariate adaptive shrinkage (mash,^63^), an empirical Bayes hierarchical modeling approach that learns correlations among (usually sparse) QTL effects across cell-types. Mash provides posterior effect estimates and the local false sign rate (lfsr) as a condition-specific measure of significance which is a more stringent analog of FDR since it requires effects to be both non-zero and correctly signed^63^. This multivariate approach identified more e/caQTL (lfsr*<*5%, **Figures 3A**–**3B**) than the initial univariate approach (**Figures 2A**–**2B**). NMJ cluster - which represents a small but distinct subset of muscle fiber nuclei at the synaptic junction with motor nerve ends saw the most increase in the significant e/caQTL, since most signals would be shared with the larger type 1, 2a and 2x muscle fiber clusters. NMJ e/caQTL also showed high pairwise QTL sign sharing with other muscle fibers (**Figures 3C**–**3D**). **Figures 3E**–**3F** show example eQTL and caQTL where the mash approach identifies significant effects (orange, confidence intervals don’t overlap 0) in the NMJ and other lower-abundance cell-types, learning shared patterns, while also identifying truly cluster-specific e/caQTL. These results show that learning from data across clusters can increase power for e/caQTL discovery.

**Figure 3:**
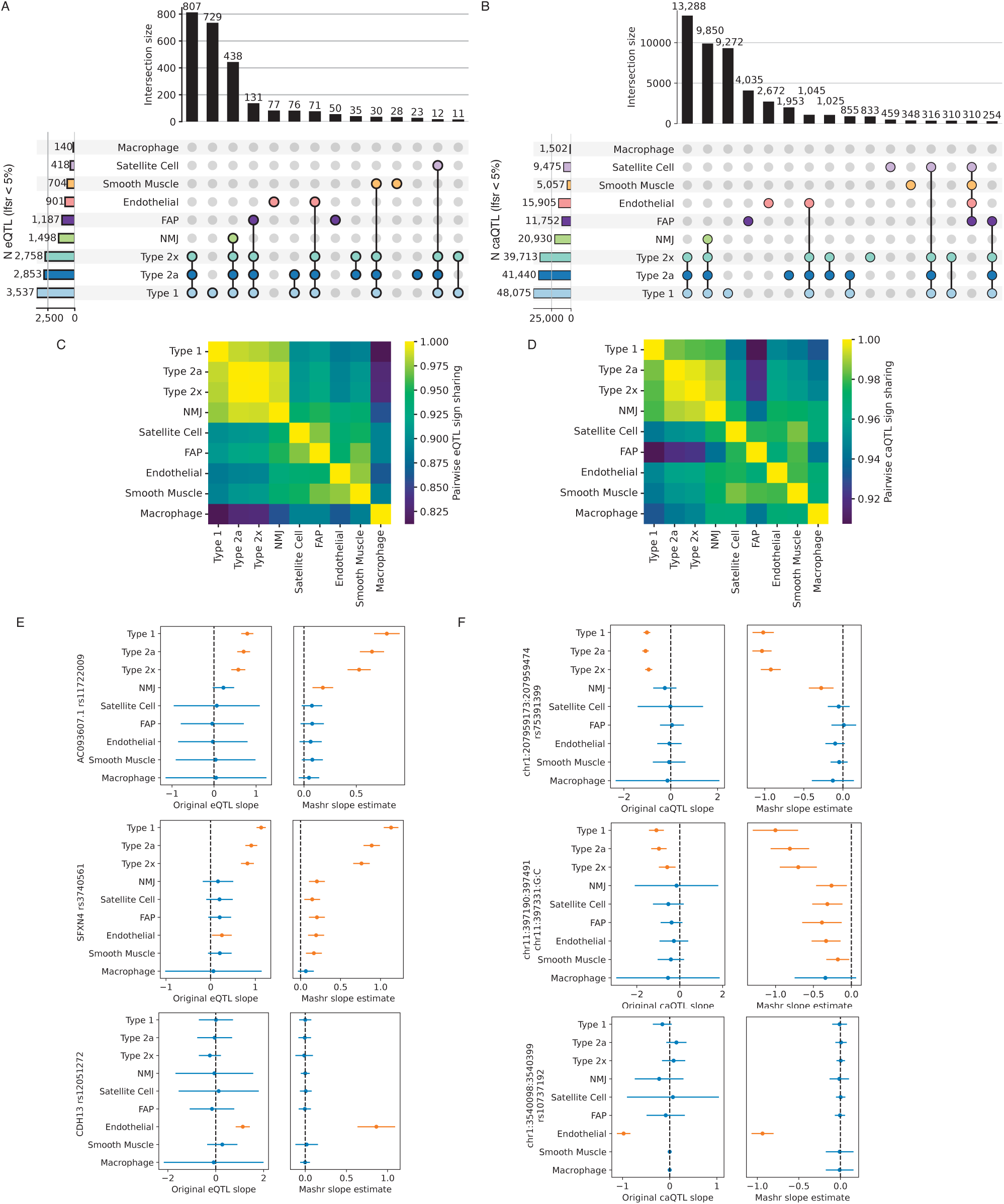
Learning patterns of e/caQTLs signal sharing across clusters inform effect estimates. (A) Fitting a mash model and estimating effects across clusters, UpSet plots show the number of shared and specific eGenes, and (B) caPeaks at a local false sign rate (lfsr)*<* 5%. (C) Fraction of eQTL or (D) caQTL effect estimates with the same sign for each pair of clusters. (E) Example eQTL and (F) caQTL showing original effects (slope) from the QTL scan and the effects estimated from mash. Bars show 95% confidence intervals. For the original eQTL results, standard errors are calculated from qvalues correcting for the total numbers of features tested after a Benjamini-Hochberg correction (hence equivalent of Mashr lfsr). For the Mashr results, estimate is the posterior mean, and error bars depict ± 1.96 * posterior standard deviations. Orange color highlights estimates where CIs don’t overlap zero.

### 2.4 Identifying context-specific e/caQTL

We next sought to identify context-specific e/caQTL effects while considering individual nucleus profiles. We sub-clustered the endothelial ATAC and RNA nuclei while defining five latent factors using liger, and identified four distinct endothelial cell contexts: capillary, arterial, venous and lymphatic (**Figure S11A**, **Figure 4A**). We then utilized the endothelial subclusters as discrete context and the latent factors as a continuous context for nuclei to test for genotype by context (GxC) interactions in a linear mixed model using CellRegMap^64^. All 198 eQTLs identified previously in the endothelial cell-type pseudobulk analyses (**Figure 2B**) showed significant (P*<*0.05) and highly correlated additive genetic (G) effect in the nucleus-level scan (P) (**Figure S11B**). Notably, using the five factors as continuous context provided higher resolution and identified more GxC interactions (92 eGenes) than discrete subcluster contexts (87 eGenes) (**Figure S11C**, **Figure 4B**). Nucleus-level caQTL modeling was impractical due to the high sparsity of the snATAC data. Therefore, we computed pseudobulk sample peak counts in each endothelial snATAC subcluster, and tested for a GxC interaction with subclusters as context for the 4,518 caPeaks identified in the initial pseudobulk scan (**Figure 4C**). These analyses identified 94% (n=4,279) of the caPeaks with significant and correlated additive G effects with the pseudobulk endothelial caQTL scan (**Figure S11D**). 43% (n=1,960) caPeaks showed significant GxC interaction effects (**Figure 4D**). These analyses demonstrate the exciting potential of snRNA/snATAC data in identifying high-resolution context-specific e/caQTL effects.

**Figure 4:**
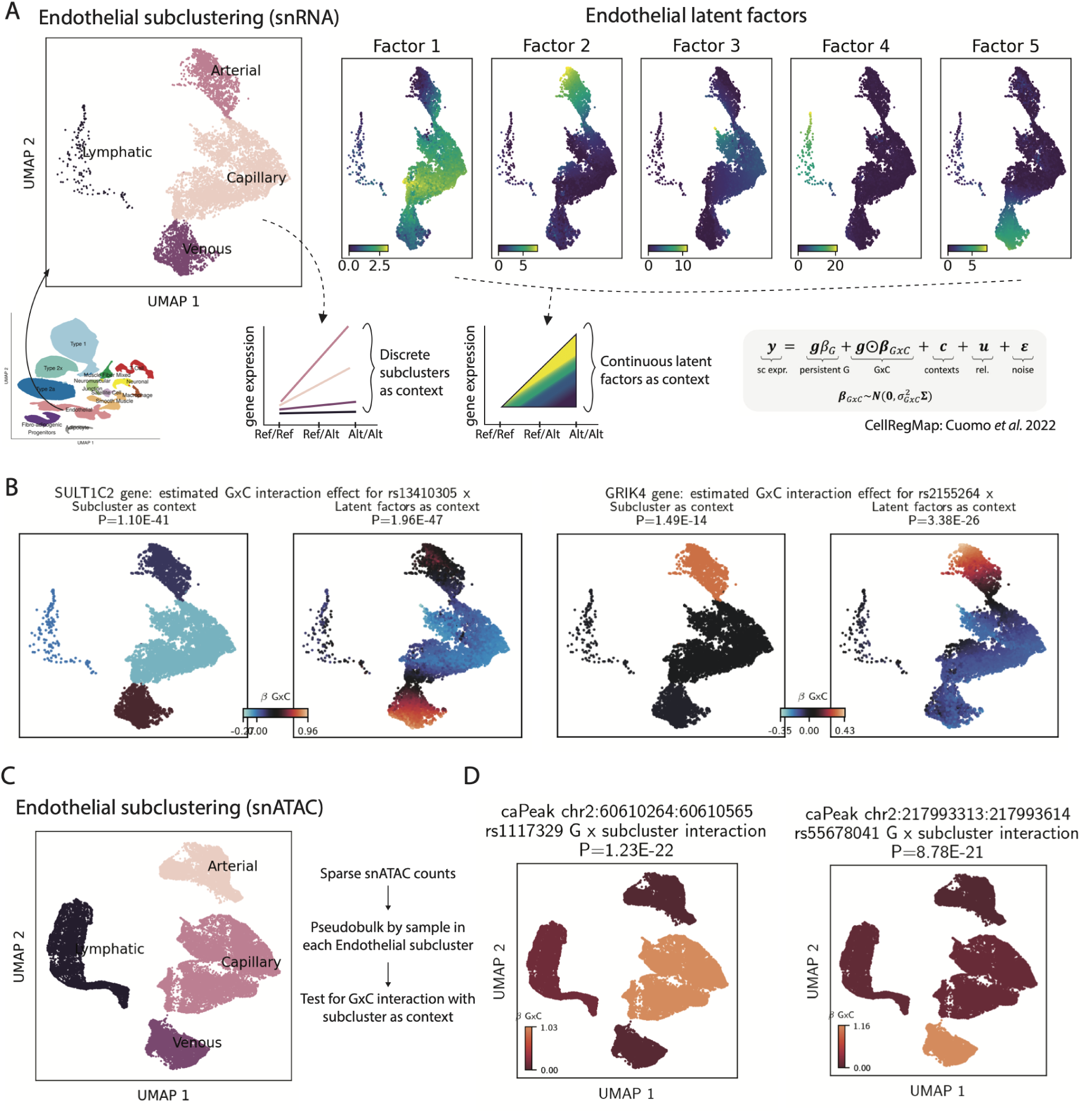
Identifying state-specific e/caQTL in endothelial cluster by testing genotype by context interaction. (A) Subclustering of the endothelial nuclei. Left: snRNA UMAP plot showing discrete subcluster contexts; right: snRNA UMAP plots show five latent factors as continuous contexts. (B) eGene examples with significant GxC interaction with subclusters (left) or factors (right) as context. (C) snATAC UMAP plot showing endothelial subclusters. Due to sparsity of snATAC data, counts were pseudobulked by sample within each subcluster prior to testing for GxC interaction. (D) caPeak examples with significant G x subcluster interaction.

### 2.5 e/caQTL finemapping, colocalization and causal inference informs cell-specific multi-omic genetic regulation

We performed genetic finemapping to identify independent e/caQTL signals and generate 95% credible sets using the sum of single effects (SuSiE) approach^65^. 284 out of 7,062 eQTL and 4,671 out of 106,059 caQTL signals could be finemapped to a single variant in the 95% credible set (**Figures 5A**–**5B**). eSNPs occurring in snATAC peaks and caSNPs occurring in the corresponding caPeaks have higher finemapping posterior inclusion probability (PIP) in the e/caQTL signal credible sets, which reinforces the quality of our e/caQTL scans and the utility of finemapping to nominate causal e/caSNPs (**Figures 5C**–**5D**). We next tested if the eQTL and caQTL signals shared causal variant(s), i.e. if the e/caQTL signals were colocalized using coloc v5^19^ (**Figure 5E**). We identified colocalized caQTL signals (coloc posterior probability for shared variant(s) (PPH4) *>* 0.5) across clusters for 1,990 eGenes; the majority (60%) of these e-caQTL colocalizations were cluster-specific (**Figure 5E**). Notably, while we detected fewer e/caQTLs in lower abundance cell-types like endothelial cells and FAP relative to muscle fibers, a larger percentage of these e/caQTLs colocalize with eGenes in only one cell-type (**Figure S12**), suggesting that QTL colocalization identifies cell-specific regulatory signals. Several relevant TF motifs were enriched in caPeaks that colocalized with an eQTL relative to caPeaks that did not colocalize (**Figure 5F**); for example, the motif for NKX2-5, a regulator of skeletal muscle differentiation^66^ is enriched in colocalized caPeaks in muscle fibers. These results suggest that e-caQTL colocalizations nominate biologically relevant gene regulatory mechanisms and emphasizes the value of our sn-e/caQTL catalog.

**Figure 5:**
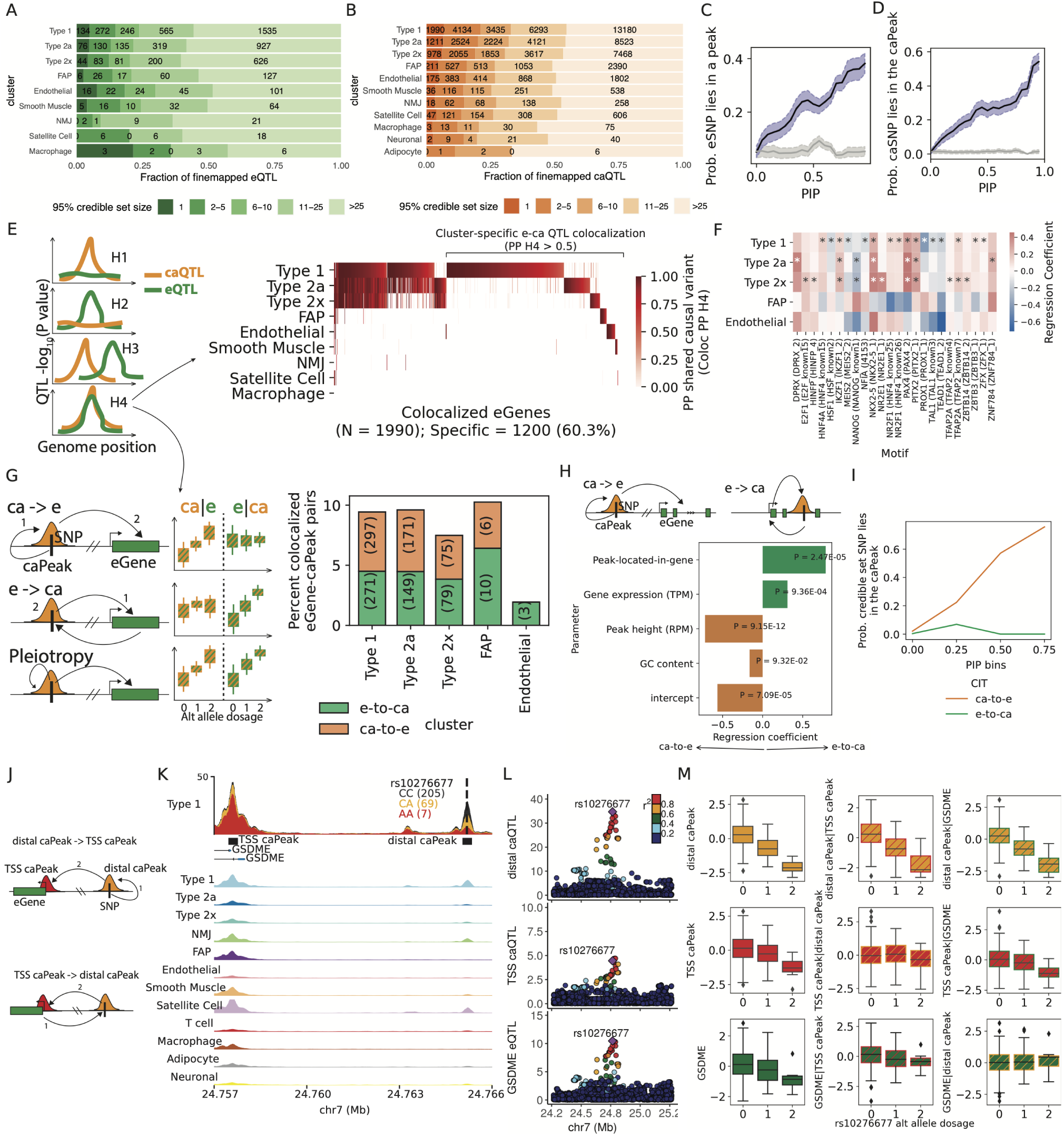
e/caQTL finemapping, colocalization and causal inference informs regulatory grammar in clusters. (A) Fraction of finemapped eQTL and (B) caQTL signals by the 95% credible set size. Probability of (C) eSNPs overlapping snATAC peaks relative to the eSNP PIPs; and (D) caSNPs overlapping the caPeak relative to the caSNP PIPs. Gray lines and confidence intervals are obtained after from shuffling e/caSNP PIPs. (E) eQTL-caQTL pairs with lead SNPs within 100kb in each cluster were tested for colocalization. Heatmap shows the posterior probability of shared causal variant (PP H4) from coloc v5. (F) TF motif enrichment in caPeaks that colocalize with eGenes relative to all caPeaks in a cluster. Clusters with at least 100 colocalized caPeaks are shown. * denotes significant logistic regression coefficient (5% FDR). (G) For each colocalized eGene-caPeak pair, causal inference tests (CIT) can inform the causal direction - Chromatin accessibility over gene expression (ca-to-e) or vice versa (e-to-ca) using e/ca SNPs as instrument variables. Barplot shows the percentage of colocalized eGene-caPeak pairs where the putative causal direction could be determined consistently from CIT and MR Steiger directionality test (5% FDR). (H) (I) (J) (K) (L) (M) continued on the next page. (H) Logistic regression modeling the causal direction between caPeak-eGene pairs with whether the caPeak lies within the eGene body, along with eGene expression (TPM,) caPeak height (RPM), and GC content. (I) Probability that a caSNP lies in the caPeak relative to caSNP PIP bins. Colors depict if the caPeak was inferred as ca-to-e or e-to-ca from CIT. (J) Where multiple caPeaks colocalize with an eGene, CIT can help delineate causal direction. (K) At the *GSDME* locus, caQTLs for a distal-peak and a TSS-peak both colocalized with the eQTL. Type 1 snATAC-seq signal track by rs10276677 genotype at this locus shows the distal-caPeak, TSS-caPeak and the *GDSME* gene TSS. Aggregate snATAC-seq in clusters are shown below. (L) Locus-zoom plots show the distal-caQTL, TSS-caQTL and the *GDSME* eQTL. (M) Causal inference between the distal-caPeak, TSS-caPeak and the *GDSME* gene using rs10276677 as the instrument variable. Boxplots show inverse normalized chromatin accessibility or gene expression relative to the alternate allele dosages at rs10276677 before and after regressing out the corresponding modality.

For colocalized e/caQTL signals, we inferred the causal relationship between chromatin accessibility and gene expression using causal inference tests (CIT) and Mendelian randomization (MR) approaches^67–69^ (**Figure 5G**). We tested if chromatin accessibility mediates the effect of genetic variation on gene expression (**Figure 5G**, row 1, “ca-to-e”), or if gene expression mediates the effect of genetic variation on chromatin accessibility (row 2, “e-to-ca”), compared to a model consistent with pleiotropic effects (row 3). In these analyses, “causal” implies that variance in the mediator determines some proportion of the variance in the outcome^67^. Since measurement errors in the molecular phenotypes can affect causal inference, we conservatively required consistent causal direction reported by both the CIT and the MR Steiger directionality test, and also performed sensitivity analyses that measured how consistent the inferred direction was over the estimated bounds of measurement error^69^ (**Figure S13A**). We discovered 1,061 colocalized e/caQTL signal pairs as ca-to-e or e-to-ca (consistent CIT and MR Steiger directionality test, 5% FDR **Figure 5G**). The e-to-ca model may represent gene expression effects on chromatin accessibility for caPeaks within the body of the transcribed gene. To test this hypothesis, we modeled the inferred causal direction in a logistic regression coding e-to-ca as 1 and ca-to-e as 0, adjusting for caPeak height (reads per million mapped reads, RPM), eGene expression level (transcripts per million mapped reads, TPM), caPeak GC content and a binary variable specifying if the caPeak was located within the eGene body. This model fit was better than a model without the caPeak-within-eGene body term (likelihood ratio test P = 1.5e-4). We found that e-to-ca caPeaks occurred within the eGene body significantly more than ca-to-e caPeaks (regression coefficient = 0.79, P = 2.47×10^-5^; **Figure 5H**), indicating that colocalized e/caQTL caPeaks in the gene body are more likely to be influenced by the act of transcription across the underlying DNA region. ca-to-e caPeaks were higher (CPM) than e-to-ca caPeaks (coefficient = −0.72, P = 9.15×10^−12^), whereas e-to-ca eGenes were more highly expressed than ca-to-e eGenes (coefficient = 0.31, P = 9.36×10^-4^).

High PIP caSNPs were more likely to occur within ca-to-e caPeaks than e-to-ca caPeaks (**Figure 5I**), consistent with expectation for caPeaks that are causal on eGenes. For TSS-distal ca-to-e caPeaks where additional caPeaks were identified in TSS+1kb upstream region of the eGene (**Figure 5J**), the distal caPeak was often causal on the TSS-caPeak as well (**Figure S13B**), Fisher’s exact test P = 4.0×10^−17^). For example, a distal caPeak ∼7.6 kb from the *GSDME* gene TSS is causal on both *GSDME* gene expression (CIT P = 5.4×10^-5^) and a TSS-caPeak accessibility (CIT P = 4.2×10^-5^) (**Figures 5K**–**5M**). These analyses support an enhancer model for the ca-to-e caPeaks where the caSNP affects chromatin accessibility at the TSS-distal caPeak that then regulates gene expression.

We highlight a locus on chromosome 8 where two independent caQTL signals for a caPeak tagged by caSNPs rs700037 and rs1400506 (**Figure S13C**), both of which lie within the caPeak (**Figure S13D**) are colocalized with two independent eQTL signals for the lincRNA gene *AC023095.1* (PPH4 0.99 and 0.76). This caPeak is specific for the type 1 fiber cluster (**Figure S13D**). Considering the independent signals as instruments, we identified the caPeak to be causal on the *AC023095.1* gene expression (CIT P value 2.11×10^−07^) (**Figure S13E**). Collectively, these results demonstrate how signal identification, finemapping, colocalization and causal inference analyses illuminate cell-specific causal event chains for the regulatory element, target gene and causal variant(s).

### 2.6 Cell-specific e/caQTL and GWAS signal integration to inform disease/trait regulatory mechanisms

To identify mechanisms underlying disease/trait associations, we integrated our e/caQTL signals with GWAS signals. We considered 302 publicly available disease/trait GWAS datasets from the UK Biobank (UKBB), along with 17 other GWAS datasets that included other skeletal muscle-relevant diseases/traits such as T2D, fasting insulin, WHR, body mass index (BMI), creatinine, and others. To further assess the relevance of skeletal muscle regulatory elements in T2D and related metabolic trait heritability, we profiled the histone marks H3K27ac (associated with enhancer and promoter activity) and H3K27me3 (associated with repressed chromatin) using CUT&Tag in skeletal muscle tissue. Enrichment of H3K27ac signal at TSSs of highly expressed genes confirmed the high-quality of this dataset (**Figures S14A**–**S14H**). We used stratified-LD score regression (S-LDSC) to compute GWAS enrichment in muscle snATAC cluster and bulk chromatin peaks^70–72^ (**Figure 6A**). Muscle fiber snATAC peaks were enriched for atrial fibrillation, creatinine, height, and pulse rate (consistent with the previous Zhang *et al.* [42] study). Notably, muscle fibers were enriched for T2D, along with fasting insulin and modified Stumvoll insulin sensitivity index (ISI) - two key measures of insulin resistance (**Figure 6A**). FAPs were enriched for various traits such as waist-to-hip ratio, bone mineral density, height, and ocular trait signals among others. Skeletal muscle H3K27ac peaks were enriched for ISI, although to a lesser extent than the muscle fiber snATAC peaks, confirming the importance of skeletal muscle in the insulin resistance phenotype and the added value in snATAC data over bulk chromatin profiles. Type 1 fiber peaks containing caSNPs were enriched to overlap T2D signals whereas peaks containing eSNPs or peaks without e/caSNPs were not enriched (after subsampling all three peak sets to the same number of peaks) (**Figure 6B**). These results indicate that trait-associated genetic variants are especially enriched in open chromatin peaks that are sensitive to genetic variation, and further highlight the importance of sn-caQTL data in identifying key disease associated regulatory elements.

**Figure 6:**
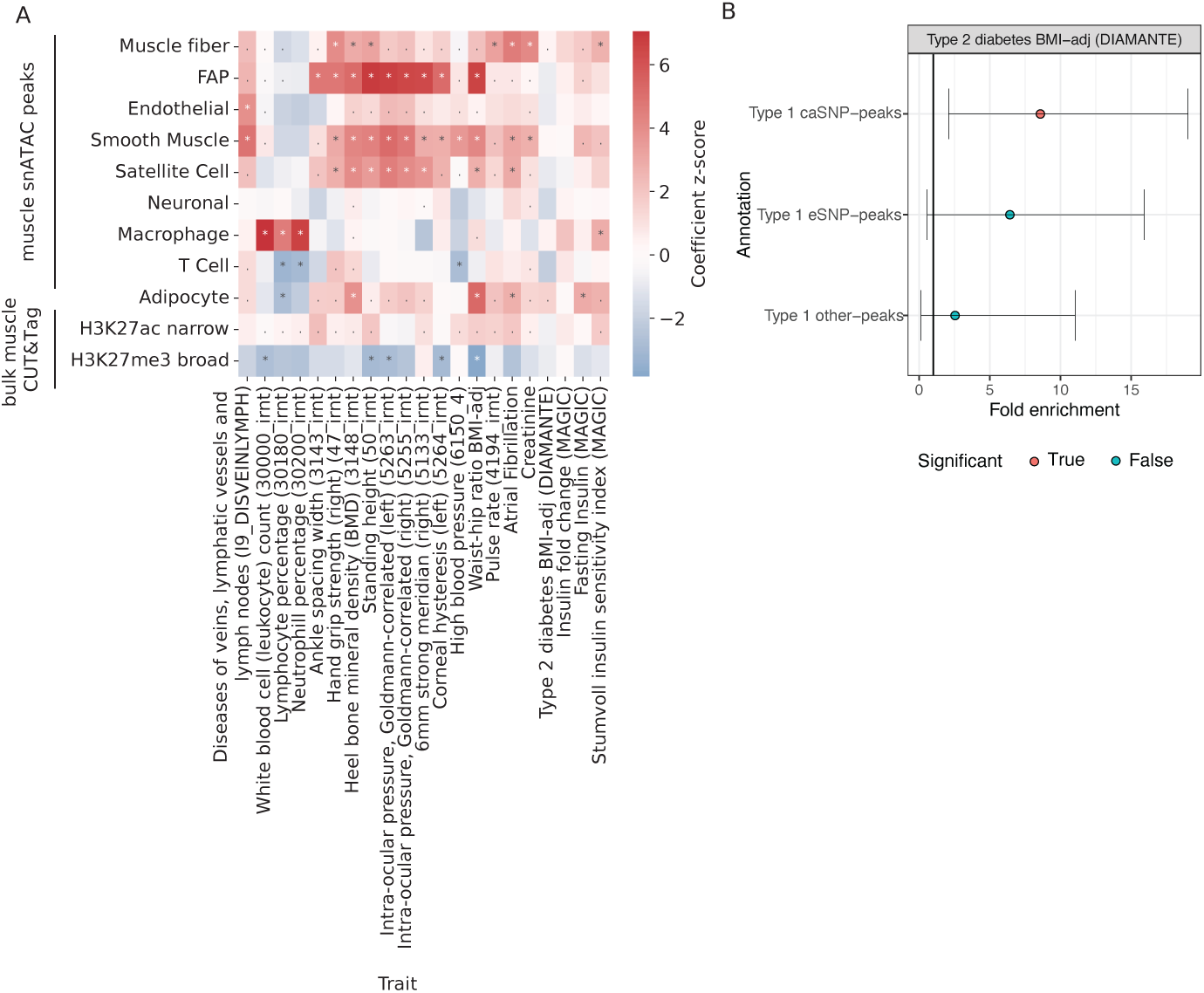
Enrichment of GWAS traits in cluster snATAC peaks. (A) GWAS enrichment in cluster snATAC peak features. Heatmap shows the LDSC regression coefficient Z scores. (B) T2D GWAS Enrichment fin type 1 fiber snATAC peaks that contain a caSNP or eSNP or peaks that do not overlap e/caSNPs. Error bars represent the 95% confidence intervals. * = FDR *<* 5% on the regression coefficient, and. = FDR *<* 5% on the heritability enrichment.

Focusing on a shortlist of 38 relevant diseases/traits, we identified 3,487 GWAS signals colocalized with e/caQTL from our study (**Figures 7A**–**7B**, **Figure S15A**), the vast majority (2,791 signals, 80%) of which were GWAS-caQTL (not GWAS-eQTL) colocalizations (**Figure 7C**). Since coloc results can be sensitive to the prior probability for the SNP being associated with both traits (p12), we performed sensitivity analyses relative to the p12 prior (**Figures S15B**–**S15D**) and include the minimum p12 prior for PPH4*>*0.5 as a potential QC metric for colocalization analyses. We highlight GWAS signals for T2D, BMI, and fasting insulin that colocalize with e/caQTL across the tested clusters, both in a shared and cell-specific manner (**Figure 7D**, **Figures S15E**–**S15F**). We also identified caQTL specific to individual muscle fiber types colocalized with several GWAS trait signals (select examples shown in **Figure S16**). In addition to eQTL, we systematically integrated snATAC co-accessibility data from Cicero^49^ as an orthogonal approach to nominate target genes. For each colocalized T2D GWAS signal, we considered if the caPeak was in the TSS region or was co-accessible with a TSS-peak of a gene; and further if the caQTL colocalized GWAS signal had a nominal eQTL association with the nominated target gene in that cluster (**Figure 7D**, bottom heatmap).

**Figure 7:**
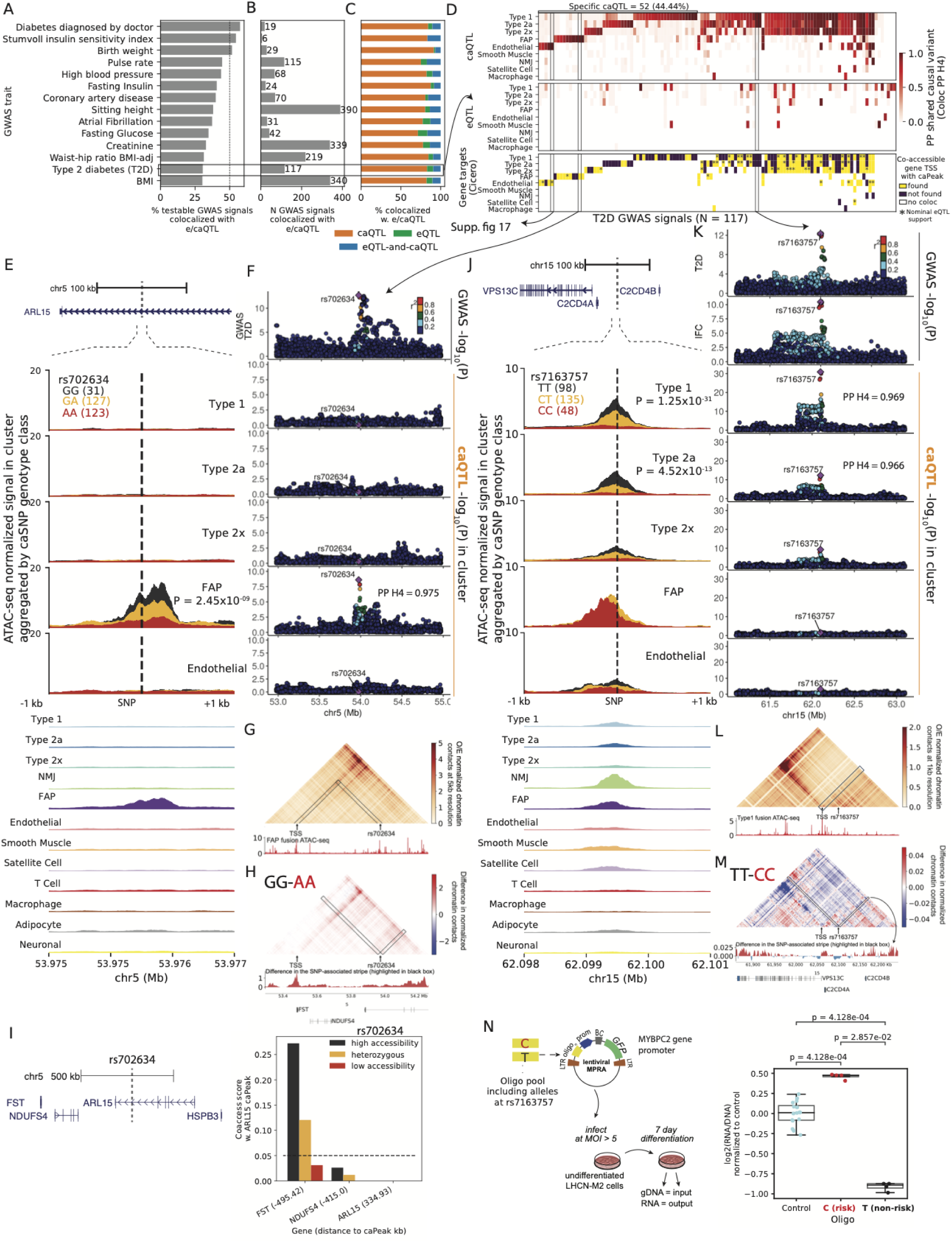
Integrating e/caQTL signals with GWAS informs disease/trait relevant regulatory mechanisms. (A) Percentage and (B) Number of GWAS signals across traits that colocalize with e/caQTL signals across the five clusters. (C) Proportion of colocalized GWAS signals (from B) that colocalize with only caQTL or only eQTL or both e-and-caQTL. (D) (E) (F) (G) (H) (I) (J) (K) (L) (M) (N) continued on the next page. (D) Heatmaps showing T2D GWAS signal colocalization with caQTL (top) and eQTL (middle). Target gene predictions using snATAC co-accessibility (Cicero) between colocalized caPeak and gene TSS peak are shown in the bottom heatmap. * indicates that the GWAS hit also had a nominally significant eQTL P value for the Cicero-nominated gene in that cluster. (E) T2D GWAS signal at the *ARL15* locus is colocalized with an FAP caQTL. The genomic locus is shown at the top, followed by zooming into a *±*1kb neighborhood of the caSNP rs702634. snATAC-seq profiles in five clusters by the caSNP genotype are shown, followed by aggregate profiles across clusters. (F) Locuszoom plots showing the *ARL15* GWAS signal (top) followed by the caQTL signal in five clusters. (G) Hi-C chromatin contacts at 5kb resolution imputed by EPCOT using the FAP snATAC-seq data (shown below the heatmap) in a 1Mb region over rs702634. (H) Difference in the predicted normalized chromatin contacts using FAP ATAC-seq from samples with the high accessibility genotype (GG) and low accessibility genotype (AA) at rs702634. Interactions with rs702634 highlighted in black are shown as a signal track below the heatmap. (I) Genes in the 1Mb neighborhood of the *ARL15* gene. Chromatin co-accessibility scores between the caPeak and TSS peaks for the neighboring genes, classified by genotype classes at rs702634. Distance of the TSS peak to the caPeak in kb is shown in parentheses. (J) GWAS signals for T2D and insulin fold change (IFC) at the *C2CD4A/B* colocalize with a caQTL in type 1 and type 2a fibers. The genomic locus, snATAC-seq profiles by the caSNP genotype and aggregated profiles are shown. (K) Locuszoom plots showing the *C2CD4A/B* GWAS and caQTL signals. (L) Micro-C chromatin contacts imputed at 1kb resolution by EPCOT using the type 1 snATAC-seq showing rs7163757 and the neighboring 500kb region. (M) Difference in the predicted normalized chromatin contacts by rs7163757 genotype. Interactions with rs7163757 highlighted in black are shown as a signal track below. (N) A massively parallel reporter assay in the muscle cell line LHCN-M2 tested a 198bp element centered on the caSNP rs7163757. Enhancer activity is measured as log2(RNA/DNA) normalized to controls.

The *GLI2* locus T2D GWAS signal (P = 4.2×10^-9^) is colocalized (PPH4 = 1.0) with a caQTL identified specifically in the endothelial cells (P = 1.37×10^−11^, **Figures S17A**–**S17B**), and the caSNP rs11688682 (PIP=1.0) occurs within the caPeak. While we didn’t identify any colocalized eQTL with this GWAS signal, alternative approaches helped nominate a target gene. We employed a deep learning framework capable of predicting the epigenome, chromatin organization and transcription (EPCOT)^73^ to impute high-resolution 3D chromatin contacts (Micro-C) using the endothelial ATAC profile. This approach predicted high contacts of the caSNP-caPeak region with the *INHBB* gene TSS, nominating the gene as a target (**Figure S17C**). Notably, we detected allelic differences in the predicted contacts, where the homozygous high accessibility genotype (GG) showed higher contacts with the *INHBB* gene than the homozygous low accessibility genotype (CC) (**Figure S17D**). The caPeak was co-accessible with the TSS peaks of genes *RALB* and *INHBB* in a genotype specific manner (**Figure S17E**); and the caSNP was nominally associated with *INHBB* expression (P=0.02).

The *ARL15* locus T2D GWAS signal (P = 7.7×10^−14^) is colocalized (PPH4 = 0.975) with an FAP-specific caQTL (P = 2.5×10^-9^) (**Figures 7E**–**7F**). EPCOT predicted high chromatin contact frequency of the caSNP rs702634 region with the *FST* gene TSS (**Figure 7G**), and the predicted contacts were higher with the homozygous high accessibility genotype (GG) compared to the homozygous low accessibility genotype (AA) at the caSNP (**Figure 7H**). This FAP-specific caPeak is present in the analogous cell type at the orthologous region in the rat genome, and its allelic enhancer activity was validated in a luciferase assay in human mesenchymal stem cells^41^. The caPeak was highly co-accessible with the *FST* gene TSS peak in a genotype-specific manner (**Figure 7I**). The nominated target gene for this GWAS signal, *FST*, encodes follistatin, which is involved in increasing muscle growth and reducing fat mass and insulin resistance^74–77^.

The *C2CD4A/B* locus T2D GWAS signal (P = 2.6×10^−13^) colocalizes (PPH4 = 0.969, 0.966) with caQTL signals in the type 1 and type 2a fibers (P = 1.25×10^−31^, 4.52×10^−13^) (**Figures 7J**–**7K**). This GWAS signal is also identified for fasting glucose and insulin fold change (IFC) post 2 hour oral glucose tolerance test (OGTT) - a measure of insulin sensitivity^78^. The caSNP rs7163757 lies within the caPeak; the T (T2D non-risk) allele is associated with higher chromatin accessibility (**Figure 7J**). Notably, this caPeak was not found as a type I skeletal myocyte cis regulatory element in the Zhang *et al.* [42] snATAC atlas. EPCOT predicted high chromatin contacts with the *VPS13C* gene TSS (**Figure 7L**), higher for the high accessibility genotype (TT) compared to the low accessibility genotype (CC) (**Figure 7M**). We didn’t detect an eQTL for *VPS13C* in muscle fibers, however, the caSNP is associated with *VPS13C* expression in whole blood (GTEx) P=2.8×10^-7^). While this caQTL is observed in muscle fibers, the snATAC peak is strongest in the lower-abundance NMJ cluster, where co-accessibility analyses also predict the *VPS13C* as the target gene (**Figure 7J**, **Figures S17F**–**S17G**). An siRNA-mediated knock-down of *VPS13C* in an adipocyte cell line affected the cell-surface-abundance of the glucose transporter GLUT4 upon insulin stimulation^78^, implicating the nominated target gene, *VPS13C*, in insulin resistance mechanisms^79^. We validated the enhancer activity of the caPeak 198 bp distal regulatory element centered on caSNP rs7163757 in a massively parallel reporter assay (MPRA) framework in the LHCN-M2 human skeletal myoblast cell line. The T2D risk allele C showed significantly higher activity relative to the empty vector control (P = 4.1×10^-4^) which was significantly higher than the activity of the non-risk T allele (P value = 2.9×10^-2^, **Figure 7N**). Previously, Kycia *et al.* [80] reported that rs7163757 occurred in accessible chromatin in pancreatic islets, the risk allele C showed higher enhancer activity in rodent islet model systems, and this allele was also associated with higher C2CD4A/B gene expression, thereby implicating this T2D GWAS signal in islet dysfunction, which was supported by an independent publication^81^. Our results highlight skeletal muscle fibers as another key cell type where this signal could modulate the genetic risk for T2D and insulin resistance through the *VPS13C* gene.

Collectively, these results demonstrate the importance of the snATAC modality and caQTL information in nominating mechanisms underlying GWAS associations and identifying causal variants in disease-relevant cell types.

## 3 Discussion

In this study, we present population-scale single-nucleus profiling of chromatin accessibility and gene expression on 287 frozen human skeletal muscle biopsies. We multiplexed 40 or 41 samples in each batch using a randomized block design to control for sample variables. Demultiplexing the data downstream using known genetic variation enabled reduced costs, helped protect against batch effects, allowed genetic detection of doublets, and overall increased rigor of the work. The integration and joint-clustering of multi-omic modalities provided a comprehensive view of the cell-specific molecular landscape within human skeletal muscle.

We identified 7,062 eQTL and 106,059 caQTL across the clusters. Concordant e/caQTL effects across clusters supported the high-quality of our e/caQTL scans. Chromatin accessibility directional allelic effects discovered from the caQTL scans mirrored the DNA-binding preferences of TF motifs which is a powerful demonstration of the depth of information snATAC and caQTL data capture. Notably, we identified 14-fold more caQTL compared to eQTL, which can be attributed to two factors: first, more peaks were tested for caQTL than genes for eQTL, and second, chromatin accessibility modality is likely an overall more proximal molecular trait to genetic variation than gene expression in the sequence of causal events, which likely contributes to the larger enhancer effects we observed and therefore results in higher power to detect caQTL with the same sample size.

The majority (80%) of GWAS signals colocalized with only caQTL rather than eQTL, in part because we detected many more caQTL than eQTL. As a corollary, we identified fewer triple GWAS-caQTL-eQTL colocalizations, which limited our efforts in using eQTL to identify target genes inferring the causal direction between omic modalities. It is becoming evident that eQTL alone fall short in fully elucidating the regulatory architecture of GWAS loci^82,83^. Our analyses revealed an intrinsic distinction between e- and caQTLs that may help reconcile these observations. Active TSS regions contained higher effect eSNPs compared to caSNPs whereas stretch enhancer regions, which are enriched for cell-type-relevant GWAS signals^8,13,84^, contained higher effect caSNPs compared to eSNPs. Therefore, eQTL scans identify signals largely in gene TSS regions, whereas caQTL scans are able to identify strong effects in cell-specific distal enhancer elements enriched for GWAS signals.

Because complex traits are influenced by both genetic and environmental effects, examining gene expression in the conditions most relevant for disease could be more informative. The larger genetic effects on stretch enhancer chromatin accessibility could propagate to gene expression effects under specific environmental conditions. Alasoo *et al.* [85] provided support for this hypothesis using bulk RNA and ATAC data in a macrophage model system where ∼60% of eQTL identified only under stimulatory conditions (response eQTL) were caQTL in the basal state. Aracena *et al.* [86] also showed that basal epigenomic profiles are strongly predictive of the transcriptional response to an antigen in immune cells. Another study reported that response-eQTL overlapped basal-caQTL in a human neural progenitor system^87^. These studies, along with our data, suggest that chromatin in cell-identity stretch enhancers is primed to potentiate changes in gene expression under relevant conditions. Future larger studies may indeed identify more eQTLs. However, if the relevant gene is not expressed at the basal state, an eQTL won’t be identified for caQTL variants even with increased sample size unless the appropriate stimulatory condition is available. Notably, recent sn-multiome studies observing lower cell-state resolution from chromatin accessibility compared to transcription also posited that cells could retain a primed or permissive chromatin landscape that can allow dynamic state transitions in response to relevant conditions^48,88^.

About half of GWAS-caQTL colocalizations were cluster-specific across traits, with many specific for the lower powered (due to nuclei abundance) Endothelial and FAP clusters, which adds to the importance of single nucleus chromatin accessibility profiling in identifying cell-specific genetic regulatory elements. Our snATAC caQTL data help delineate heterogeneity in the mechanistic pathways shaping T2D pathophysiology. We show the *GLI2* signal is most relevant for endothelial cells and the *ARL15* signal targets the *FST* gene in FAPs, implicating an interplay of fat and muscle mass regulation by these progenitor cells in T2D. We find evidence for the *C2CD4A/B* T2D GWAS signal, previously implicated in islet dysfunction through inflammatory cytokine-responsive *C2CD4A/B* genes, to also be involved in glucose uptake mechanisms in muscle fibers through the *VPS13C* gene. Cell types such as FAPs and endothelial occur in other T2D-relevant tissues such as adipose; comparing the snRNA/snATAC and e/caQTL profiles for these cell types from a wider array of tissues will help glean the similarities and differences in disease mechanisms in related cell type populations. Layering sn-e/caQTL colocalization information over GWAS signals across multiple relevant tissues will help generate a conceptual “signal scoreboard” that can help prioritize cell types, regulatory elements, target genes and causal variants(s) for each GWAS signal towards experimental validation.

To date, there have been some single cell/nucleus eQTL studies^89–94^, few sn-caQTL studies^28,95^; however, these all had modest sample sizes, and were mainly in blood cell types or cell lines. There are no population-scale single cell/nucleus studies in skeletal muscle and none with both RNA and ATAC modality for hundreds of samples in any tissue. Our work bridges a large gap in knowledge in that it is the first study identifying both sn-eQTL and sn-caQTL across hundreds of samples in any tissue. Our findings emphasize the need to consider chromatin accessibility in addition to gene expression when investigating the functional mechanisms underlying complex traits, and serves as a template for multi-omics maps in other tissue and disease contexts.

### 3.1 Limitations of the study

In our single-nucleus study, most nuclei were identified as muscle fibers; this distribution of cell type proportions was especially skewed since muscle fibers are multi-nucleated. Lower abundance clusters had relatively less power to identify e/caQTL. Generating single-nucleus data involves several tissue-dependent considerations and challenges. Other examples include diseased liver that can have fibrosis and brain that has high lipid content, both of which can make processing of frozen tissue, like in this study, challenging. Pancreas has high levels of RNase activity which degrades the snRNA modality quality. Comparing e/caQTL effect sizes across clusters enabled more precise effect estimates and identified more significant associations across clusters, especially for the NMJ cluster. Instead of QTL scans within discrete clusters, identifying contiguous cell states through latent embedding and related approaches^64,96^ helps mitigate power issues and can identify state-specific QTLs. Approaches such as deeper sequencing, pre-selecting relevant cell types via fluorescence activated cell sorting (FACS) could further enrich for targeted rare cell types and allow for greater power to identify QTLs^97–99^. Cleaner nuclei preps with low ambient transcripts and better approaches to adjust for these would enable retrieving more quality nuclei from rare cell types. The feasibility of these approaches again heavily depends on the tissue. Using our down-sampling results, for 200 samples, we find that ∼75 nuclei per sample yields ∼1,000 eQTL and *>*10,000 caQTL. The number of nuclei to target in future experiments can thus be calculated based on the expected proportion of rare cells of interest in a given tissue. Signal upscaling via deep learning methods such as AtacWorks and PillowNet^100,101^ is another possible avenue to enhance caQTL scans in lower abundance cell types. The multiome protocol for profiling RNA and ATAC on the same nucleus was not available when our FUSION study samples were processed. However, it has several advantages including 1) ease in genetic demultiplexing, sample assignment, and clustering as these analyses can be done on one modality (eg snRNA) and can then be mapped easily to the other modality or by weighting both modalities; 2) established cross-modality approaches to link regulatory elements to genes. We recommend all future studies to perform multiome profiling.

We recognize that while our findings offer cell-specific mechanistic insights at hundreds of loci, comprehensive orthogonal testing of the identified e/caQTL associations and e/caQTL-GWAS colocalizations to confirm their impact on disease remains a critical step for future studies. Several studies have demonstrated large-scale validation of existing genome-wide associations using functional allelic MPRA assays, CRISPRi screens among others^102–104^. We demonstrate successful MPRA in the LHCN-M2 skeletal muscle cell line, for the first time, thus providing feasibility for these future studies.

In further work, co-activity QTLs (e.g. QTLs on co-expression, co-accessibility) could provide additional resolution to regulatory mechanisms. Cell-specific caQTL and eQTL maps could be used for biobank-scale polygenic scoring of individuals. Collapsing caQTL peaks and eQTL genes into pathways and aggregating pathway-level effects based on individual genotype dosages would allow for cell- and pathway-specific polygenic scores, paving the way for partitioning tissue-agnostic polygenic risk scores into cell-specific personalized pathophysiological risk profiles.

## 4 Methods

### 4.1 Sample collection

#### 4.1.1 FUSION cohort

The Finland-United States Investigation of NIDDM Genetics (FUSION) study is a long-term project aimed at identifying genetic variants that contribute to the development of type 2 diabetes (T2D) or affect the variability of T2D-related quantitative traits. To conduct the FUSION Tissue Biopsy Study, we obtained *vastus lateralis* muscle biopsy samples from 331 individuals across the glucose tolerance spectrum, including 124 with normal glucose tolerance (NGT), 77 with impaired glucose tolerance (IGT), 44 with impaired fasting glucose (IFG), and 86 with newly-diagnosed T2D^29^.

To ensure the validity of the study results, certain individuals were excluded from the study, including those receiving drug treatment for diabetes, those with conditions that could interfere with the analysis (such as cancer, inflammatory diseases, or skeletal muscle diseases), those with conditions that increase hemorrhage risk during biopsy (such as hemophilia, von Willebrand’s disease, or severe liver disease), those taking medications that increase hemorrhage risk during the biopsy (such as warfarin), those taking medications that could confound the analysis (for example oral corticosteroids, or other anti-inflimmatory drugs such as infliximab or methotrexate), and those under 18 years of age.

Clinical and muscle biopsy visits were conducted at three different study sites (Helsinki, Savitaipale, and Kuopio). The clinical visit included a 2-hour four-point oral glucose tolerance test (OGTT), BMI, waist-to-hip ratio (WHR), lipids, blood pressure, and other phenotypes measured after a 12-hour overnight fast, as well as health history, medication, and lifestyle questionnaires. The clinical visit was conducted an average of 14 days before the biopsy visit.

The muscle biopsies were performed using a standardized protocol. Participants were instructed to avoid strenuous exercise for at least 24 hours prior to the biopsy. After an overnight fast, approximately 250 mg of skeletal muscle from the vastus lateralis was obtained using a conchotome, under local anesthesia with 20 mg/mL lidocaine hydrochloride without epinephrine. A total of 331 muscle biopsies were collected by nine experienced and well-trained physicians at the three different study sites between 2009 and 2013, with three physicians performing the majority of the biopsies. All physicians were trained to perform the biopsy in an identical manner. The muscle samples were cleaned of blood, fat, and other non-muscle tissue by scalpel and forceps, rinsed with NaCl 0.9% solution, and frozen in liquid nitrogen within 30 seconds after sampling. Muscle samples were then stored at −80 degrees Celsius.

### 4.2 Sample preparation, snRNA-seq and ATAC profiling

The frozen tissue biopsy samples were processed in ten batches, each consisting of 40-41 samples. These batches were organized using a randomized block design to protect against experimental contrasts of interest including cohort, age, sex, BMI and stimulatory condition (relevant for a smaller cohort not focused on in this study) (**Figures S1A**–**S1E**). Samples in each batch were pulverized in four groups of 10 or 11 samples (each sample weighing between 6-9 mg) using a CP02 cryoPREP automated dry pulverizer (Covaris 500001) and resuspended in 1 mL of ice-cold PBS. Following, the material from all 40/41 samples was pooled together and nuclei were isolated. We developed a customized protocol (protocol S1, supplementary text) derived from the previously published ENCODE protocol https://www.encodeproject.org/experiments/ENCSR515CDW/ and used it to isolate nuclei, which is compatible with both snATAC-seq and snRNA-seq. The desired concentration of nuclei was achieved by re-suspending the appropriate number of nuclei in 1X diluted nuclei buffer (supplied by 10X genomics for snATAC, and RNA nuclei buffer (1% BSA in PBS containing 0.2U/uL of RNAse inhibitor) for snRNA). The nuclei at appropriate concentration for snATAC-seq and snRNA-seq were submitted to the University of Michigan Advanced Genomics core for all the snATAC-seq and snRNA-seq processing on the 10X Genomics Chromium platform (v. 3.1 chemistry for snRNA-seq). Nuclei to profile each modality from each batch were loaded onto 8 channels/wells of a 10X chip at 50k nuclei/channel concentration. For snRNA-seq, the libraries were single-ended, 50 bp, stranded. For snATAC-seq, the libraries were paired-ended, 50 bp. The sequencing for each modality and batch was performed on one NovaSeq S4 flowcell.

### 4.3 Muscle multiome sample

We obtained “multiome” data, i.e. snATAC-seq and snRNA-seq performed on the same nucleus for one muscle sample as part of newer ongoing projects in the lab. We used 70mg of pulverized human skeletal muscle tissue sample. The sample was pulverized using an automated dry cryo pulverizer (Covaris 500001). We developed a customized protocol (hybrid protocol with sucrose) from the previously published ENCODE protocol, and used it to isolate nuclei for single nuclei multiome ATAC and 3’GEX assay. The desired concentration of nuclei was achieved by re-suspending the appropriate number of nuclei in 1X diluted nuclei buffer (supplied by 10X genomics). The nuclei at the appropriate concentration for single nuclei multiome ATAC and 3’GEX assay was processed on the 10X genomics chromium platform. 20K nuclei were loaded on one well of the 8 well strip.

### 4.4 Genotyping and imputation

The FUSION cohort samples were genotyped using DNA extracted from blood on the HumanOmni2.5 4v1 H BeadChip array (Illumina, San Diego, CA, USA) during a previous study^30^. The Texas and Sapphire cohort samples were genotyped using DNA extracted from blood on the Infinium Multi-Ethnic Global-8 v1.0 kit. Probes were mapped to Build 37. We removed variants with multi mapping probes and updated the variant rsIDs using Illumina support files Multi-EthnicGlobal D1 Mapping-Comment.txt and annotated.txt downloaded from https://support.illumina.com/downloads/infinium-multi-ethnic-global-8-v1-support-files.html. We performed pre-imputation QC using the HRC-1000G-check-bim.pl script (v. 4.2.9) obtained from the Marc McCarthy lab website https://www.well.ox.ac.uk/∼wrayner/tools/ to check for strand, alleles, position, Ref/Alt assignments and update the same based on the 1000G reference (https://www.well.ox.ac.uk/∼wrayner/tools/1000GP_Phase3_combined.legend.gz). We did not conduct allele frequency checks at this step (i.e. used the –noexclude flag) since we had samples from mixed ancestries.

For all samples, we performed pre-phasing and imputation using the Michigan Imputation Server^105^. The standard pipeline (https://imputationserver.readthedocs.io/en/latest/pipeline/) included pre-phasing using Eagle2^106^ and genotype dosage imputation using Minimac4 (https://github.com/statgen/Minimac4) and the 1000g phase 3 v5 (build GRCh37/hg19) reference panel (The 1000 Genomes Project Consortium 2015). Post-imputation, we selected biallelic variants with estimated imputation accuracy (r2) *>* 0.3, variants not significantly deviating from Hardy Weinberg Equilibrium P*>*1e-6, MAF in 1000G European individuals *>* 0.05.

### 4.5 snRNA-seq data processing and quality control

snRNA: We mapped the reads to the human genome (hg38) using STARsolo https://github.com/alexdobin/STAR/blob/master/docs/STARsolo.md (v. 2.7.3a). We performed rigorous quality control (QC) to identify high-quality droplets containing single nuclei (**Figures S1F**–**S1G**). We required the following criteria: 1) nUMI *>* 1000; 2) fraction of mitochondrial reads *<* 0.01; 3) identified as a “singlet” and assigned to a sample using Demuxlet^107^ 4) identified as “non-empty”, i.e. where the RNA profile was statistically different from the background ambient RNA signal, using the testEmtpyDrops function from the Dropletutils package^108^; and 5) passing the cluster-specific thresholds for the estimated ambient contamination from the DecontX package^109^. This led to a total of 255,930 pass-QC RNA nuclei, 180,583 from the FUSION cohort. These individual qc steps are further described below.

### 4.6 snATAC-seq data processing and quality control

We made barcode corrections using the 10X Genomics whitelist using an approach implemented by the 10X Genomics Cell Ranger ATAC v. 1.0 software via a custom python script and counted the number of read pairs from each droplet barcode. We trimmed the adapter sequences using cta https://github.com/ParkerLab/cta and generated updated fastqs by replacing the cellular barcodes with the corrected cellular barcodes, while selecting reads corresponding to cellular barcodes that had at least 1000 pairs. Droplets with less than 1000 read pairs would not contain useful/high quality data from single nuclei and so were removed from processing. We mapped the reads to the human genome (hg38) using bwa mem (v. 0.7.15-r1140)^110^ with flags “-I 200,200,5000 -M”. We performed rigorous quality control (QC) and retained high-quality droplets based on the following definitions (**Figures S1H**–**S1I**): 1) 4,000 *<* high quality autosomal alignments (HQAA) *<* 300,000, 2) transcription start site (TSS) enrichment ≥ 2, 3) mitochondrial fraction *<* 0.2. For each snATAC-seq library bam file, we used the subset-bam tool (v. 1.0.0) https://github.com/10XGenomics/subset-bam to subset for the selected cellular barcodes, and used SAMtools to filter for high-quality, properly-paired autosomal read pairs (-f 3 -F 4 -F 8 -F 256 -F 1024 -F 2048 -q 30). To identify droplets containing a single nucleus “singlet” and determine the sample identity, we used the Demuxlet^107^ tool. For each library (8 10X channels/wells in each of the 10 batches, N=80), we ran Demuxlet using default parameters providing the snATAC-seq library bam files the genotype vcf files containing all samples included in that batch and selected all the droplets assigned as singlets. This led to a total of 3,69,792 pass-QC ATAC nuclei, 2,68,543 from the FUSION cohort.

#### 4.6.1 Two-stage Demuxlet pipeline

Multiplexing 40/41 samples in each batch in a randomized block study design helped protect against batch effects and it was cost-effective approach. To identify droplets containing a single nucleus “singlet” and determine the sample identity, we used the Demuxlet^107^ tool. For each library (8 10X channels/wells in each of the 10 batches, N=80), we ran Demuxlet using default parameters providing the library bam files the genotype vcf files containing all samples included in that batch and selected all the droplets assigned as singlets. Background/ambient RNA contamination can influence singlet assignments, so we accounted for that next. We performed clustering of these pass-qc RNA droplets and annotated clusters using known marker genes. A large proportion of our data was muscle fiber nuclei, this is expected since muscle fibers are multi-nucleated. Therefore, a large proportion of ambient RNA would come from muscle fiber cells. Observing the barcode-nUMI rank plots (**Figure S1F**), we considered droplets with less than 100 reads as unlikely to contain an intact nucleus and therefore representative of the ambient RNA profile. Top 100 genes contained top ∼30% of ambient RNA reads (**Figure S2A**). Most abundant genes in the ambient RNA were expectantly mitochondrial and muscle fiber genes such as MYH1, MYH7 etc (**Figure S2B**). We reasoned that “masking” top n% of these top genes should reduce ambiguity arising due the ambient RNA, enabling more droplets to be assigned as a singlet. We tested masking to n% of genes from Demuxlet and observed that masking the top 30% of genes in the ambient RNA maximized singlet assignment (**Figure S2C**). We therefore completed a second Demuxlet run masking top 30% genes, and any new droplets that were identified as singlets to the set of selected droplets. The singlet nuclei recovered from the masked stage 2 came mostly from lower abundance non-fiber clusters (**Figure S2D**) (using cluster labels identified downstream).

#### 4.6.2 Adjusting RNA counts for overlapping gene annotations

RNA mapping and gene quantification using STARsolo outputs a “GeneFull” matrix that quantifies intronic+exonic reads and a “Gene” matrix that quantifies only exonic reads. For our nuclear RNA expriment, we used the GeneFull matrices for all downstream applications. As of the STAR version 2.7.3a which was used in our analysis, in case of overlapping gene annotations, the program renders some read assignments ambiguous and therefore some genes receive less counts in the GeneFull matrix compared to the Gene matrix. We observed the distribution of count differences between the exon+intron (Gene-Full) and exon (Gene) matrices for each gene across all 80 libraries and created a list of genes where this difference was consistently negative in at least 10 libraries. We then created custom counts matrices keeping the “Gene” counts for these 6,888 selected genes and kept the “GeneFull” counts for all other genes.

#### 4.6.3 Ambient RNA adjustment

We used DecontX (celda v. 1.8.1, in R v. 4.1.1)^109^ to adjust the nucleus x gene expression count matrices for ambient RNA. Taking all the qc’ed RNA nuclei up to this stage (N = 260,806), we identified cell type clusters using Liger (rliger R package v. 1.0.0)^45^. Liger employs integrative non-negative matrix factorization (iNMF) to learn a low-dimensional space in which each nucleus is defined by both dataset-specific and shared factors called as metagenes. It then builds a graph in the resulting factor space, based on comparing neighborhoods of maximum factor loadings. We selected the top 2000 variable genes using the selectGene function and clustered with number of factors k=20 and regularization parameter lambda=5 along with other default parameters as it identified expected clusters (**Figure S3A**). We then ran DecontX on a per-library basis using the decontX() function, passing our custom created RNA raw matrices (adjusted for overlapping gene annotations) for the QC’ed nuclei, barcodes with total UMIs *<* 100 for the background argument, cluster labels from liger, and set the delta parameter (prior for ambient RNA counts) as 30. This prior value was more stringent than the DecontX default of 10 and it was selected after exploring the parameter space and observing that delta=30 better reduced fiber type marker gene such as *MYH7*, *MYH2* counts in rarer clusters such as Endothelial, Satellite Cell, while retaining respective marker gene *VWF* and *PAX7* counts (**Figure S3B**). Since the decontamination is sensitive to the provided cluster labels, we performed a second clustering using adjusted counts from the first DecontX run to obtain better optimized cluster labels. We also included the snATAC modality for this clustering. Liger’s online integrative non-negative matrix factorization (iNMF) algorithm was used at this step^45,46^ which enabled efficient processing of this large snATAC+snRNA dataset by iteratively sampling a subset of nuclei at a time. We selected the clustering with liger k=19, lambda=5, epoch=5, batchsize=10,000 along with other default parameters (**Figure S3C**). We then performed a second DecontX run using raw snRNA matrices (adjusted for overlapping gene annotations), droplets with UMIs *<* 100 as background, delta set to 30 while including the updated snRNA cluster labels.

DecontX also estimates fraction of ambient RNA per nucleus. We used this metric to further filter out RNA nuclei. We observed that this metric varied across clusters, and the immune cell, muscle fiber mixed and the smooth muscle clusters has a visible population of nuclei with high estimated ambient RNA fraction (**Figure S3D**). We therefore fitted two Gaussians for these three clusters per batch and removed nuclei that obtained the probability of being from the high contamination population *>* probability of being from the low contamination population (**Figure S3E**). For the rest of the clusters, we removed nuclei with estimated ambient RNA *>* 0.8. We retained all pass QC nuclei and used rounded decontaminated counts for the final joint clustering and all downstream analyses.

### 4.7 Joint clustering and cell type annotation

We jointly clustered snRNA and snATAC from the FUSION cohort and the one multiome muscle sample using Liger’s online iterative non-negative matrix factorization (iNMF) algorithm version (https://github.com/MacoskoLab/liger/tree/online)^45,46^. Liger’s online iNMF was capable of processing our large dataset because it factorizes the data using mini-batches read on demand (we used a mini-batch size = 10,000 nuclei). We factorized the RNA nuclei first using adjusted gene by nucleus count matrices for autosomal protein-coding genes as input. We used the following parameters: top 2000 variable genes, k=21, lambda=5, epoch=5, max iterations=4, batchsize=10,000, along with other default parameters. We then performed quantile normalization to align across batches. Next, we projected the snATAC datasets using gene (gene body + 3kb promoter region) by nucleus fragment counts as input to the existing RNA factorization. This process uses the existing gene loading in the factors for computing the factor loading in ATAC nuclei. We then quantile normalized the snATAC data and finally used the Louvain graph based community detection algorithm with resolution 0.04 to identify clusters. This process resulted in a joint clustering without batch or modality specific effects (**Figure S4A**). We annotated the clusters using known marker gene expression patterns (**Figure S4B**).

### 4.8 ATAC-seq peak calling and consensus peak feature definition

We created per-cluster snATAC-seq bam files by merging reads from all pass-QC ATAC nuclei for each cluster. We randomly subsampled bam files to 1 Billion reads and called narrow peaks using MACS2 (v. 2.1.1.20160309)^111^. We used BEDTools bamToBed^112^ to convert the bam files to the BED format, and then used that file as input to MACS2 callpeak (command “macs2 callpeak -t atac-$cluster.bed –outdir $cluster -f BED -n $cluster -g hs –nomodel –shift −100 –seed 762873 –extsize 200 -B –keep-dup all”) to call narrow peaks. We removed peaks overlapping the ENCODE blacklisted regions^113^, and selected peaks passing 0.1% FDR from macs2. We then defined a set of consensus snATAC-seq peak summits across all 13 clusters. We considered the filtered narrow peak summits across all clusters and sorted by MACS2 q value. We sequentially collapsed summits across clusters within 150bp and retained the most significant one, identifying N=983,155 consensus summits (**Figures S5A**–**S5C**). Aggregating ATAC-seq signal over broad peaks in a cluster while centering on the left-most summit showed the second summit usually occurred ∼300bp away (**Figure S5D**), in line with the nucleosome length being ∼147 bp^114^. We therefore considered each consensus summit extended by 150 bp on each side as the consensus peak-feature for all downstream analyses. To visualize the signal, we converted the bedGraph files output by MACS2 to bigWig files using bedGraphToBigWig^115^.

### 4.9 Identification of cell type-specific genes and GO enrichments

Differential gene expression analyses between all pairs of cell types were performed to identify cell type-specific genes. Muscle fiber nuclei clusters (Type 1, Type 2a, Type 2x, Neuromuscular junction, Muscle Fiber Mixed) were merged for this analysis due to their expected similarity. For each pair of cell types we used DESeq2^116^ to call differential genes between the cell types. Samples with less than 3,000 genes detected in either of the cell types were dropped, as were genes with less than 3 counts across all of the samples (when combining the cell types). The DESeq2 analysis was done in a paired sample fashion. A gene was considered to be a cell type-specific gene for cell type X if that gene was more highly expressed in cell type X than in all other cell types (5% FDR).

### 4.10 Comparison to snATAC atlas

Per-cell type comparisons to the snATAC atlas from^42^ were performed using a modified version of the logistic regression-based technique described previously^41^. First, narrowPeaks from each cell type cluster were merged to produce a set of master peaks. Next, master peaks within 5kb upstream of a GENCODE TSS (GENCODE v40;^117^) were dropped. Master peaks were annotated to muscle cell types according to whether or not they overlapped a narrowPeak in that cell type, and master peaks annotated to more than one cell type were dropped, resulting in a set of cell type-specific peaks. Next, for each of our cell types and each of the 222 cell types from^42^, we ran the logistic regression model: (master peak is specific to muscle cell type ∼ *β*_0_ + *β*_1_ *master peak overlaps peak from snATAC atlas cell type), where *β*_0_ represents a model intercept. Within each of our cell types, we then produced a matching score for each of the snATAC atlas cell types by re-normalizing the resulting model coefficient *β*_1_ to range between 0 and 1 (by dividing the coefficients by the maximum coefficient, first setting coefficients to 0 if the model p-value was not significant after Bonferroni correction or the coefficient was negative). The snATAC atlas cell type with score = 1 was determined to be the best match.

GO enrichments were performed using g:Profiler (python API, v. 1.0.0;^118^), using all genes with at least one count in one cell type as the background set.

### 4.11 Identification of cell type-specific open chromatin summits and motif enrichments

Using the per-cluster peak summit counts, we identified cell type-specific summits using the *τ* metric from^119^. As muscle fiber types show high gene expression similarity, we merged any nuclei assigned to muscle fibers (Type 1, Type 2a, Type 2x, NMJ, and Muscle fiber mixed clusters). Summits with *τ >* 0.8 were considered to be cell type-specific, and were assigned to the cell type showing greatest accessibility of that summit.

Motif enrichments were performed using the 540 non-redundant motifs from a previous study^120^, with the logistic regression model (one model per motif per cell type): summit is specific to cell type ∼ intersect + summit is TSS proximal + summit GC content + number of motif hits in summit where TSS proximal was defined as within 2kb upstream of a TSS, and the number of motif hits was determined using FIMO (v. 5.0.4, with default parameters and a 0-order Markov background model generated using fasta-get-markov^121^). We excluded two cell types (Neuronal and T cell) with less than 500 cell type specific summits and excluded cases where the model didn’t converge. A motif was considered significantly enriched if the coefficient for the “number of motif hits in summit” term was significantly positive after Bonferroni correction within each cell type. The corresponding heatmap figure displays motifs that were amongst the top 5 significantly enriched motifs by either p-value or coefficient in at least one cell type.

### 4.12 snATAC-seq coaccessiblity

We ran CICERO^49^ (v. 1.4.0; R v. 4.0.1) on the narrow peak fragment counts in each cluster to score peak-peak co-accessibility. We used UMAP dimensions 1 and 2 (**Figure 1B**) as the reduced coordinates and set window size to 500 kb. A peak was considered to be a TSS peak for a gene if it overlapped the 1kb window upstream of that gene’s TSS. If multiple TSS peaks were present for a gene, the maximum co-accessibility score was considered.

### 4.13 QTL scan in clusters

We performed expression and chromatin accessibility QTL analysis in clusters using QTLtools (v. 1.3.1-25-g6e49f85f20)^122^. The mixed muscle fiber cluster showed higher fraction of reads mapping to exon relative to the full gene body in certain batches (indicating lower quality, Supplementary note), therefore, this cluster was not considered for QTL scans and downstream analyses. We removed three samples from out QTL analyses: one because it appeared to be of non-Finnish ancestry from PCA analysis, and two others which were found to be first degree related to other samples. We created a vcf file with imputed genotypes of all the selected FUSION samples, and filtered for autosomal, bi-allelic variants with MAF ≥ 5%, non-significant deviation from Hardy-Weinberg equilibrium P*>*1×10^-6^. We performed PCA using QTLtools pca with options –scale, –center and –distance 50,000.

### 4.14 eQTL scan

We selected the following gene biotypes (Gencode V30): protein coding, lincRNA, 3prime overlapping ncRNA, antisense, bidirectional promoter lncRNA, macro lncRNA, non coding, sense intronic, and sense overlapping. For each cluster, we considered samples with at least 10 nuclei for the eQTL analysis. We generated RNA count matrices by summing up gene counts (post-ambient RNA decontamination) from nuclei for each sample in each cluster. We converted the gene counts into transcript per million (TPMs) and inverse-normalized across samples. TPM = RPK/factor, where RPK = counts/(length in kb) and factor = sum(RPK)/1M for each cluster. We used the top 10,000 genes based on median TPM to perform PCA using QTLtools. eQTL scans were performed considering variants within 250kb of gene TSSs. For each cluster, we ran test eQTL scans while considering the top 3 genotype PCs and a varying number of phenotype PCs to account for unknown biological and technical factors. We selected the number of phenotype PCs that maximized eQTL discovery as covariates **Figure S7A**. We optimized within-cluster thresholds for minimum gene counts across at least 10 samples that defined our final set of testable genes that minimized the multiple testing burden **Figure S7B**. We performed the cis eQTL scans with 1,000 permutations, then applied an across-feature multiple testing correction using the qvalue Storey function on the beta distribution adjusted P values and reported eGenes at FDR ≤ 5%.

### 4.15 caQTL scan

For each cluster, we considered samples with at least 10 nuclei for the caQTL analysis. We didn’t restrict our caQTL scans to only peaks identified in a cluster, instead considered all testable consensus peaks to allow for comparisons across clusters. We quantified each consensus feature and obtained the sum of fragment counts across all nuclei from each samples in each cluster. For an initial lenient caQTL scan, we selected all consensus features in a cluster that had at least 2 counts in at least 10 samples to test for caQTL in each cluster. We used inverse-normalized counts per million (CPMs) as quantification for caQTL. CPM = RPK/factor, where RPK = counts/(feature length in kb) and factor = sum(RPK)/1M for each cluster. We performed PCA on the inverse-normalized CPMs and included the top n phenotype PCs that maximized QTL discovery in each cluster, along with the top 3 genotype PCs as covariates. We optimized within-cluster thresholds for minimum peak counts across 10 samples that defined our final set of testable peak that minimized the multiple testing burden (**Figure S8A**). We then calculated PCs for these selected features and again optimized the number of PCs within each cluster that maximized caQTL discovery (**Figure S8B**). caQTL scans were performed using the selected samples, optimized features, 3 genotyped PCs and final set of optimized phenotype PCs considering variants within 10kb of the feature midpoint (peak summit). We performed the cis caQTL scans with 1,000 permutations, then applied an across-feature multiple testing correction using the qvalue Storey function on the beta distribution adjusted P values and reported caPeaks at FDR ≤ 5%.

### 4.16 Motif reconstruction using caQTL results

We used a library of 540 non-redundant PWMs for the motif reconstruction analyses(D’Oliveira Albanus et al. 2021). Motif hits were determined by scanning the genomic sequence in a variant-aware manner using FIMO (v. 5.0.4, with default parameters and a 0-order Markov background model generated using fasta-get-markov^121^), i.e. scanning the genomic sequence containing the reference and the alternative allele. For a given cell type and motif, we identified all lead caQTL variants or their LD r2*>*0.8 proxies that sat within the corresponding caPeak and that overlapped a motif hit (n=31 - 10,646 (27 - 42%) depending on the cell type). For each such overlapping caQTL, we calculated the caQTL allelic fold change^123^ using tensorQTL^124^. To reconstruct the motif, for each of the four nucleotides and each position in the motif, we summed the absolute value of the allelic fold change for all caQTLs overlapping that position in the motif hit and having that nucleotide as the favored (open chromatin) allele. This was converted to a probability matrix (such that the four values at each motif position summed to one) for the final reconstructed motif. To demonstrate that the observed similarity between the original and reconstructed motif was not simply a result of the fact that a motif hit was called by FIMO, we additionally reconstructed motifs based on all variants that met filtering requirements for the caQTL scan, overlapped motif hits, and were in peaks tested in the caQTL scan. To do this, for each of the four nucleotides and each position in the motif, we counted the number of variants overlapping that position in the motif hit and having that nucleotide as either the ref or the alt allele, and then converted this to a probability matrix as before.

### 4.17 mash analyses

We utilized mash^63^ to learn correlation patterns of QTL effect sizes across clusters to in turn obtain more precise effect size estimates. We considered the top 9 clusters in which both eQTL and caQTL were identified from our original e/caQTL scans (FDR*<*5%) for setting up the mash model. For both e and caQTL, we created the Bhat (effect size) and Shat (standard error) matrices for sets of “strong” and “random” tests as per the recommendations of the original authors https://stephenslab.github.io/mashr/articles/eQTL_outline.html. For eQTL, we first compiled a set of all genes that were testable across the 9 clusters (n=12,891). The “strong” tests included the top SNPs for these genes, top SNP being the one with the minimum nominal p value across the nine clusters. The “random” tests included n=50,000 randomly selected snp-gene pairs for the gene set from the original eQTL scan. For caQTL, there were 62,187 caPeaks total identified across 9 clusters (FDR*<*5%), whereas, only 20,000 peaks were testable in all 9 clusters. Therefore, for an appropriate representation of the “strong” signals, we included the union of both these sets of peaks (total n=87,003) to set up the mash model. When a peak was not testable in a cluster, we set the effect to 0 and standard error to infinity. The “strong” tests included the top SNPs for these mash peaks. The “random” tests included n=100,000 randomly selected SNP-peak pairs for the mash peak set from the original caQTL scan.

We learned the correlation structure among random tests (Vhat, function estimate null correlation simple) followed by setting up the strong and random mash data sets (function mash set data). We learned data-driven covariance matrices using strong tests, first computing PCA (function cov pca), then running the extreme deconvolution algorithm (function cov ed). We computed the canonical covariance matrices using the function cov canonical on the random set. We fit the mash model using both these covariance matrices. Lastly, we computed posterior summaries on the strong tests using the mash model fit - lfsr, posterior mean and posterior standard deviation, which are equivalent of the FDR, effect size and standard error of a QTL scan respectively. We utilized the function get pairwise sharing to plot the pairwise sign sharing between each pair of clusters (**Figures 3C**–**3D**). While plotting the original eQTL effects (**Figures 3E**–**3F**), we obtained qvalues using Benjamini-Hochberg on the strong tests nominal p values to compute the standard errors, so as to make the results comparable to mash posterior summaries.

### 4.18 context-specific QTL

We used CellRegMap^64^ to identify context-specific e/caQTL. We first separated the RNA and ATAC nuclei identified as endothelial cell-type and jointly clustered using the liger online iNMF approach as described previously for the main clustering. We computed five latent factors for the RNA nuclei first using the following parameters: top 2000 variable genes, k=5, lambda=5, epoch=5, max iterations=4, batchsize=5,000, along with other default parameters. We performed quantile normalization to align across batches followed by projecting the snATAC datasets. Louvain clustering at a resolution of 0.025 identified four endothelial subclusters, which we annotated using known marker genes.

The CellRegMap linear mixed model is of the form: *y* = *gβ* + *g* ∗ *β*_GxC_ + *c* + *u* + *ɛ*, where single-cell gene expression values of a given gene (*y*) are modeled as a function of a persistent genetic effect (*g*), GxC interactions (*g*∗), additive effects of cellular context (*c*), relatedness (*u*) and residual noise (*ɛ*). For snRNA, we tested the top SNP-eGene pairs for the 198 eQTLs identified for the endothelial cluster from our initial pseudobulk eQTL scan. We set up the CellRegMap model using either the subcluster labels as discrete context or the five latent factors as continuous context. We computed the kinship matrix to represent the relatedness within the data including the fact each sample contributes multiple nuclei. We considered genotyped variants, pruned these to LD r^2^*<*0.2 using the plink flag –indep-pairwise 250 50 0.2, followed by using flag –make-king square. We transformed this matrix to a positive semi-definite matrix by adding the minimum eigenvalue to the diagonal elements. We normalized the endothelial nuclei by gene expression matrix to log2(counts per million (CPM) + 1) using scanpy preprocessing functions pp.normalize total(adata, target sum=1e6, exclude highly expressed=True), followed by pp.log1p(adata, base=2). We included age, sex, batch, BMI and the fraction of mitochondrial reads in nuclei as additional covariates in the model. We first tested linear association with genotype using the function run association, then tested interaction using the function run interaction, followed by estimating betas using the function estimate betas.

For snATAC, we tested the top SNP-caPeaks pairs for the 4,518 caPeaks for the endothelial cluster from our initial pseudobulk eQTL scan. Since snATAC data is much more sparse than snRNA, nucleus-level linear mixed models were impractical. We instead computed pseudobulk sample counts in each subcluster, and included subcluster as the discrete context. The count normalization, covariates and kinship matrix were performed as described for snRNA.

### 4.19 QTL finemapping

We used the sum of single effects (SuSiE)^125^ approach to identify independent e and caQTL signals and obtain 95% finemapped credible sets. We used QTLtools to adjust for the covariates optimized for e or caQTL scans and inverse-normalized the residuals. We used these adjusted phenotypes along with the sample genotype dosages for variants in a 250kb window in the susie function along with the following parameters: number of signals L=10, estimate residual variance=TRUE, estimate prior variance=TRUE, min abs cor=0.1.

### 4.20 Relationship between caQTL effect size, caSNP MAF, and caQTL peak presence in scATAC atlas

Type 1 muscle fiber caPeaks were grouped based on the open chromatin allele frequency (calculated using the FUSION samples) and the caQTL effect size (absolute value of the slope, binned by 0-0.4, 0.4-0.8, 0.8-1.2, 1.2-1.6, and 1.6-2.0). We then calculated the fraction of the caPeaks within that bin that overlapped with a Type I Skeletal Myocyte peak from^42^.

### 4.21 caPeak chromatin state enrichments

CaPeak enrichment in chromatin states was computed using the Skeletal Muscle Female (E108) chromatin states (15-state model) from Roadmap Epigenomics^126^. First, muscle ATAC peaks were lifted from hg38 to hg19 using liftOver (kentUtils v. 343^127^). For each of the Type 1, Type 2a, and Type 2x cell types, we then ran the logistic regression:

peak is caPeak ∼ intercept + peak size + overlaps state 1 + … + overlaps state 15

where peak size was set as the average peak reads per million across samples. Only peaks tested for caQTL were included in the model. caPeaks were enriched for a state if the coefficient for the corresponding state term in the model was significantly positive after Bonferroni correction (Bonferroni correction performed within each cell type, across the 16 non-intercept terms).

### 4.22 Motif enrichment in caPeaks

Motif enrichments were performed using the 540 non-redundant motifs from^120^, with the logistic regression model (one model per motif per cell type):

peak is caPeak ∼ intercept + peak is TSS proximal + peak GC content + peak size + number of motif hits in peak where TSS proximal was defined as within 2kb upstream of a TSS, peak size was set as the average peak reads per million across samples, and the number of motif hits was determined using FIMO (v. 5.0.4, with default parameters and a 0-order Markov background model generated using fasta-get-markov^121^). Only peaks tested the caQTL scans were included in each model. A motif was considered significantly enriched if the coefficient for the “number of motif hits in summit” term was significantly positive after Bonferroni correction within each cell type. The corresponding heatmap figure displays motifs that were amongst the top 3 significantly enriched motifs by either p-value or coefficient in at least one cell type.

### 4.23 eQTL and caQTL colocalization

We used coloc v5^19^ to test for colocalization between e and ca QTL. We used the e and ca QTL finemapping output from SuSiE over the 250kb window as inputs to coloc v5. We considered colocalization between two signals if the PP H4 *>* 0.5.

### 4.24 Causal inferrence between chromatin accessibility and gene expression

For all pairs of colocalized eGenes and caPeaks, we inferred the causal chain between chromatin accessibility and gene expression using two orthogonal approaches - a mediation-based approach causal inference test (CIT, v2.3.1)^67,68^ and a Mendelian randomization approach MR Steiger directionality test^69^. We required consistent direction from both CIT and MR Steiger at 5% FDR to consider an inferred causal direction between an eGene and caPeak pair.

#### 4.24.1 CIT

To test if an exposure mediates an effect on an outcome, CIT uses genetic instruments (eg SNPs) requiring a set of mathematical conditions to be met in a series of regressions under a formal hypothesis testing framework. If a SNP (L) is associated with an outcome (T) only through an exposure (G), outcome when conditioned on the exposure should be independent of the SNP. The conditions therefore are: (i) L is associated with T, (ii) L is associated with G conditional on T, (iii) T is associated with G conditional on L and (iv) T is independent of L conditional on G. For each pair of caPeak and eGene for which one or more independent caQTL and eQTL signal(s) colocalized, we ran four CIT models each returning an omnibus P value- a) eSNP(s) -*>* eGene -*>* caPeak (P e-to-ca-causal), b) eSNP(s) -*>* caPeak -*>* eGene (P e-to-ca-revCausal), c) caSNP(s) -*>* caPeak -*>* eGene (P ca-to-e-causal) and d) caSNP(s) -*>* eGene -*>* caPeak (P ca-to-e-revCausal). We included sample batch, age, sex, BMI and top 3 genotype PCs as covariates in the CIT model. For each model, we computed the omnibus FDR values using the fdr.cit function to account for multiple testing. To infer a caPeak causal on an eGene, we required q-ca-e-causal *<* 0.05, q-ca-e-revCausal *>* 0.05, q-e-ca-causal *>* 0.05 and q-e-ca-revCausal *<* 0.05, and vice versa to infer an eGene causal on a caPeak. We note that eGene-caPeak pairs without a putative causal CIT prediction could be truly independent or could have a causal relationship obscured by measurement error.

#### 4.24.2 MR Steiger directionality test

In an MR-based approach, the genetic instrument (SNP) is used as a surrogate for the exposure to estimate its causal effect on an outcome, by scaling the association of SNP and outcome by the association between SNP and exposure. This approach is considered less susceptible to bias from measurement errors or confounding^69^. For each pair of caPeak and eGene for which one or more independent caQTL and eQTL signal(s) colocalized, we used the mr steiger function (TwoSampleMR R package version 0.5.6) to test both caPeak and eGene as exposure over the other modality as outcome. To infer a caPeak causal on an eGene, we required ca-to-e “correct causal direction” as “True” at 5% FDR, and e-to-ca “correct causal direction” as “False” at 5% FDR, while estimating steiger test q values using the R qvalue package (http://github.com/jdstorey/qvalue). For each model, we provided the respective QTL scan sample sizes and set r xxo = 1, r yyo = 1, r exp = NA and r out = NA to estimate the sensitivity ratio - which computes over the bounds of measurement errors in the exposure and outcome, how much more often is one causal direction observed versus the other. The higher the sensitivity ratio, more robust is the inferred causal direction to measurement errors.

### 4.25 GWAS enrichment in ATAC-seq peak features

We computed enrichment of GWAS variants in ATAC-seq peak features using stratified-LD score regression (s-LDSC)^70,128^. We downloaded GWAS summary statistics for 17 traits relevant for skeletal muscle such as T2D, glycemic traits, aftrial fibrillation. Where required, we lifted over the summary stats onto hg38 using the UCSC liftOver tool. We formatted the summary stats according to LDSC requirements using the ldsc munge sumstats.py script, which included keeping only the HapMap3 SNPs with minimum MAF of 0.01 (as recommended by the LDSC authors). We also downloaded several LDSC-formatted UKBB GWAS summary statistics from the Benjamin Neale lab website^129^ https://nealelab.github.io/UKBB_ldsc/downloads.html. We selected primary GWASs on both sexes for high confidence traits with h2 significance *>* z7, following guidelines described on the Ben Neal lab blog https://nealelab.github.io/UKBB_ldsc/details.html. We created a baseline model with cell type agnostic annotations such as MAF, coding, conserved regions, along with other epigenomic annotations such as DNase hypersensitiviy sites (DHS), transcription factor binding sites (TFBS) that are obtained from across multiple cell types. These annotations are among the list of baseline annotations included in the original LDSC paper^128^. The various annotation files (regression weights, frequencies, etc.) required for running LDSC were downloaded from https://data.broadinstitute.org/alkesgroup/LDSCORE/GRCh38/. We set up LDSC to test snATAC-seq peak features (consensus peak summit features that overlapped a peak summit called in that cluster) and the bulk muscle CUT&Tagin peaks along with the baseline annotations. LD scores were calculated using the Phase 3 1000 Genomes data. LDSC reports two types of output: first, the total heritability explained by SNPs in the annotation, which includes heritability attributable to other overlapping annotations in the baseline; and second, joint-fit regression coefficient for each annotation, that quantifies the contribution of that annotation to per-SNP heritability. The former estimates if the annotation contributes to the overall heritability and the latter estimates if the annotation contributes to the heritability in addition to all the other baseline annotations in the model. We reported significance using both these metrics in **Figure 6A**. We calculated coefficient P-values from the coefficient z-scores using a one-sided test assuming a standard normal distribution. We calculated FDR separately for enrichment p-values and coefficient p-values using the BH procedure and report traits with FDR¡5% for either measure.

While comparing GWAS enrichment in type 1 peaks that overlapped caSNPs, eSNPs or not e/caSNPs, since there is a large difference in the number of eSNPs and caSNPs features, we subsampled each annotations to have the same number of features: n=6,880 peaks. LDSC authors suggest that S-LDSC only produces well-calibrated p-values when annotations span at least 1.7% of 0.01cM blocks of the genome (roughly 51Mb assuming 1cM ∼ 1Mb, ten-fold larger than our current eSNP-peaks annotation)^130^. Therefore, we used an alternative enrichment approach, fGWAS^131^ and tested enrichment for the downsampled annotations.

### 4.26 eQTL and caQTL co-localization with GWAS

We considered the lead GWAS signals that if the individual study reported so; otherwise, we identified genome-wide significant (P *<* 5e-8) signals in 1Mb windows. We finemapped each GWAS signal using the available GWAS summary statistics along with 40,000 unrelated British individuals from the UKBB as the reference panel, over a 250kb window centered on the signal lead variant. We obatined pairwise r between variants using the cor() function in R on the genotype dosages for variants in the SuSiE window. We ran SuSiE using the following parameters: max number of signals L = 10; coverage = 0.95; r2.prune = 0.8; minimum absolute correlation = 0.1; maximum iterations = 10,000. We considered e/ca QTL signals where the lead variant was within 250kb of the GWAS lead variant to test for GWAS-QTL colocalization using the function coloc.susie from the coloc v5 package. We used the coloc sensitivity() function to assess sensitivity of findings to coloc’s priors. We considered two signals to be colocalized if the PP H4 *>* 0.5.

### 4.27 Imputing high-resolution 3D chromatin contact maps

We used EPCOT^73^ to impute the high-resolution 3D chromatin contact maps. EPCOT is a computational framework that predicts multiple genomic modalities using chromatin accessibility profiles and the reference genome sequence as input. We predicted chromatin contacts in genomic neighborhoods of selected caPeaks of interest using snATAC-seq from the respective cluster - either Micro-C at 1kb resolution for a 500kb genomic region or Hi-C at 5kb resolution for a 1Mb genomic region. EPCOT was trained with existing Micro-C contact maps from H1 and HFF, or Hi-C contact maps from GM12878, H1, and HFF. Both the Micro-C and Hi-C contact maps are O/E normalized (i.e., the contact values present the ratio of the observed contact counts over the expected contact counts).

We then generated Micro-C maps by the genotype at the caSNP of interest. We created genotype-specific snATAC-seq profiles by aggregating samples with either homozygous reference or homozygous alternate genotypes at the caSNP of interest. We downsampled the data using Picard when required to make the two profiles have similar depth. We respectively incorporated the reference of alternate allele in the DNA sequence input to EPCOT. Subsequently, we subtracted the predicted contact values associated with the low chromatin accessibility genotype from the high accessibility genotype.

EPCOT’s input ATAC-seq (bigWig) processing:

bamCoverage –normalizeUsing RPGC –effectiveGenomeSize 2913022398 –Offset 1 –binSize 1 –blackListFileName ENCODE black list.bed

### 4.28 Massively parallel reporter assay for validation

#### 4.28.1 Cloning

We ordered oligos as 230 bp sequences where 197 bp comprise the variant of interest flanked on both by 98 bp of genomic context, and the additional 33 bp are cloning adapters. Within this panel, we included a set of ∼50 negative control sequences defined by a previous publication^132^ We added 20 bp barcodes via a 2-step PCR amplification process then incorporated the barcoded oligos into a modified pMPRA1 vector (a gift from Tarjei Mikkelsen^133^, Addgene #49349) upstream of the GFP reporter gene using Golden Gate assembly. After transforming and expanding in NEB 10-beta electrocompetent bacteria, we sequenced this version of the MPRA library to establish a barcode-oligo pairing dictionary. We performed a second Golden Gate assembly step to insert an ENCODE-annotated promoter for the human MYBPC2 gene in between the oligo and barcode. Finally, we used restriction cloning to port the assembled MPRA block (oligo, barcode, promoter, GFP) to a lentiviral transfer vector, which was used by the University of Michigan viral vector core to produce infectious lentiviral particles. Primer sequences used for cloning and sequencing library preparation along with the MYBPC2 promoter sequence are included in a separate table.

#### 4.28.2 MPRA Experiment

For each replicate, we infected 4×10^6^ LHCN-M2 human skeletal myoblasts with our MPRA library at an MOI of ∼10. After infection, we passaged the cells for one week to remove any unincorporated virus or contaminating transfer plasmid, then differentiated the cells for one week. We isolated RNA and gDNA from each replicate using the Qiagen AllPrep DNA/RNA mini kit. We reverse transcribed RNA into cDNA with a GFP-specific primer, then constructed indexed sequencing libraries for both the cDNA and gDNA libraries using Illumina-compatible primers.

#### 4.28.3 Data Analysis

After quality checks and filtering, we calculated the sum of barcode counts for each oligo within a replicate. We used DESeq2 v1.34.0^116^ to perform normalization and differential expression analysis. We used a nested model to identify oligos with significant activity (relative to plasmid input) and significant allelic bias (between reference and alternate alleles). All results were subject to a Benjamini-Hochberg FDR of 5%.

### 4.29 Code availability

The code used to run analyses in this work are available on [GitHub](https://github.com/ParkerLab/sn_muscle_2023)

## 4.30 Acknowledgements

We thank Sarah Graham, Cristen Willer, Sarah Hanks, Marcel den Hoed, and Parker lab members for their help and feedback for this work. This work was supported by grants R01DK072193, R01DK117960, R01DK062370, UM1DK126185, and P30AR069620.

## 4.31 Author Contributions

AV performed batch design, data processing and computational analyses, interpreted and visualized the results, prepared figures and wrote the manuscript. NM processed samples, isolated nuclei and performed the snATAC-seq and snRNA-seq experiments. PO, ZZ, FF and JM performed computational analyses and generated figures. AT and KN performed MPRA experiments and AT analyzed the data. VR contributed to visualization. ME processed FUSION biopsy samples and HS, AUJ organized sample information and metadata. NN processed FUSION genotyping data. CV performed experiments. TT and OIK performed sequencing. CG, JDW assisted in joint clustering. MY, LKW, CJ, RAD, LN, JS, TAL, ML, JT, HK contributed biopsy samples. JDW, CB, KM, JOK, JL, MB, FSC, LJS and SCJP provided supervision and mentoring. LJS and SCJP designed the study and obtained funding. SCJP organized the project and supervised all aspects of data generation, analyses and interpretation, and edited the manuscript.

## 4.32 Competing interests

SCJP has a research grant from Pfizer.

## Supplemental Information

### Supplemental Figures

**Figure S1:**
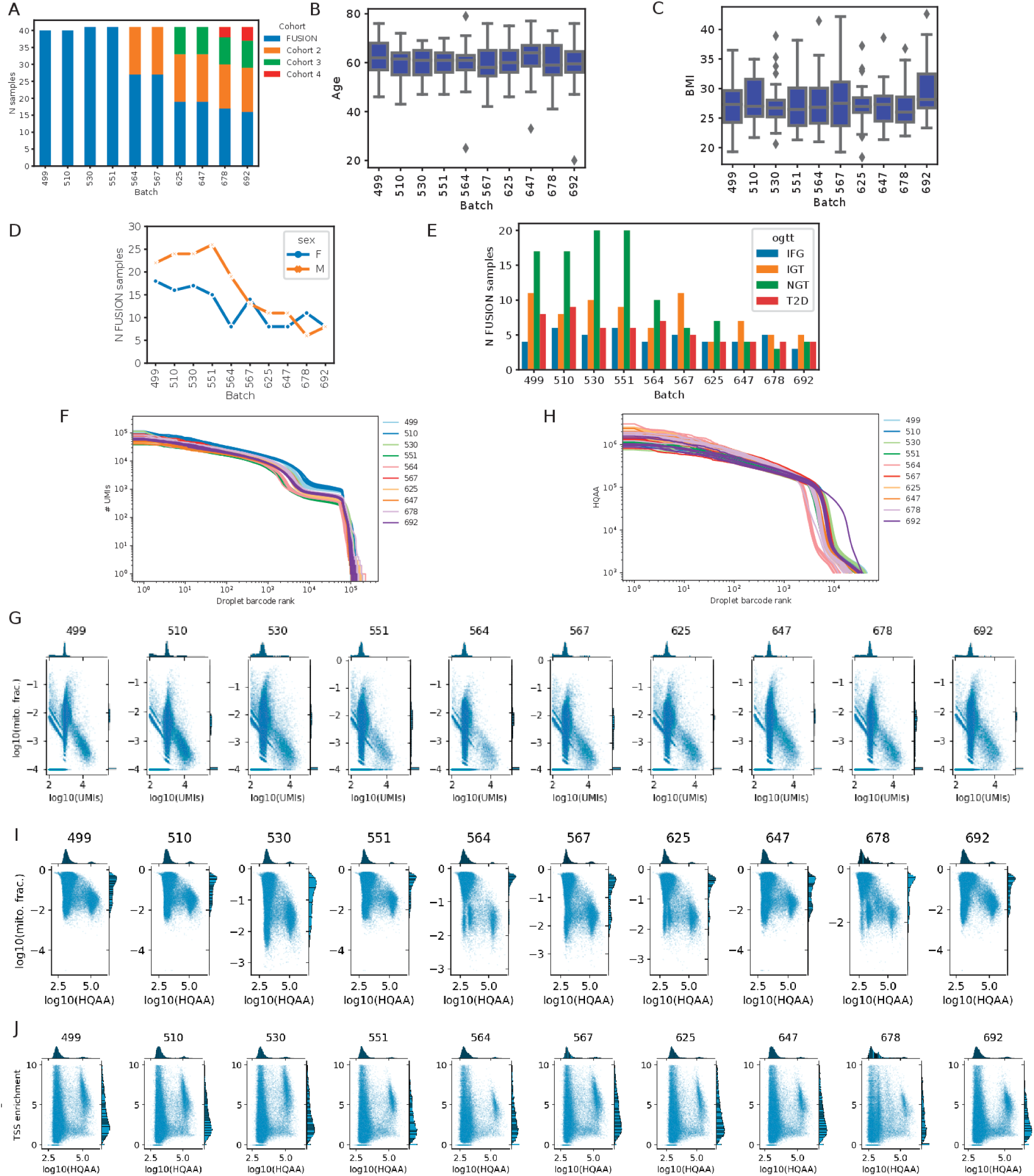
Batch design and quality control. (A) Cohort representation across batches (B) Age (C) BMI (D) Sex (E) OGTT status for individuals with samples across batches (F) snRNA-seq barcode rank plot showing number of UMIs (G) snRNA UMIs vs mitochondrial read fraction across batches (columns). (H) snATAC-seq barcode rank plot showing high quality autosomal alignments (HQAA) (I) snATAC HQAA vs mitochondrial read fraction across batches (J) snATAC HQAA vs TSS enrichment across batches. Panels G, I and J show nuclei from one 10X channel per batch.

**Figure S2:**
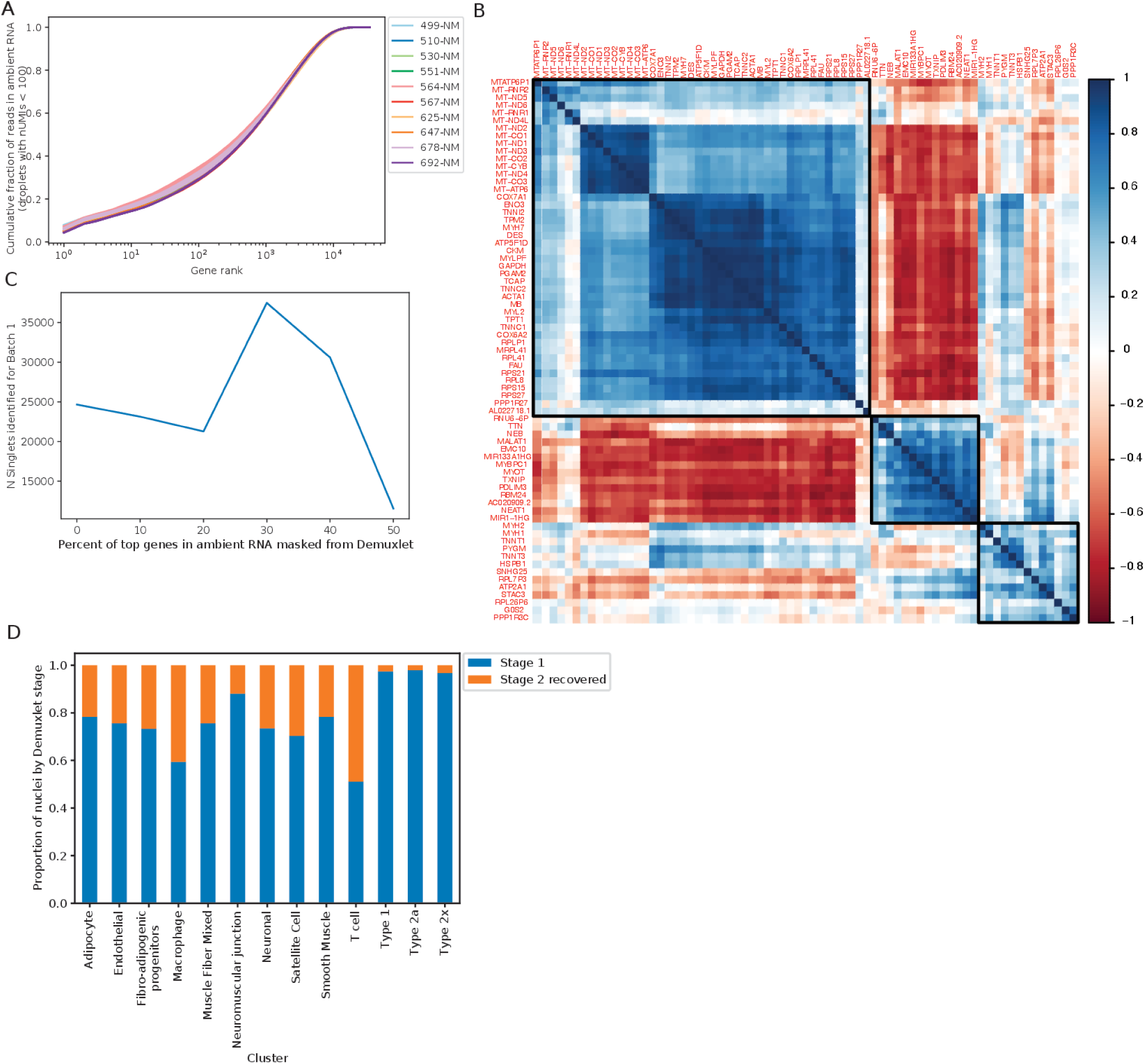
Identifying singlets and sample assignment. (A) Gene rank plot showing the number of reads per gene in the ambient RNA profile(droplets with *<* 100 UMIs) (B) Heatmap showing the pairwise Pearson correlation of the top expressed genes in the ambient RNA. (C) Titration to optimize masking genes to maximize singlet identification. Shown are the total number of singlets identified by Demuxlet for batch 1 after masking the top n% of genes expressed in the ambient RNA profile. (D) Proportion of singlet nuclei recovered at demuxlet stage 1 (default) and stage 2 (’masked’) for each cluster.

**Figure S3:**
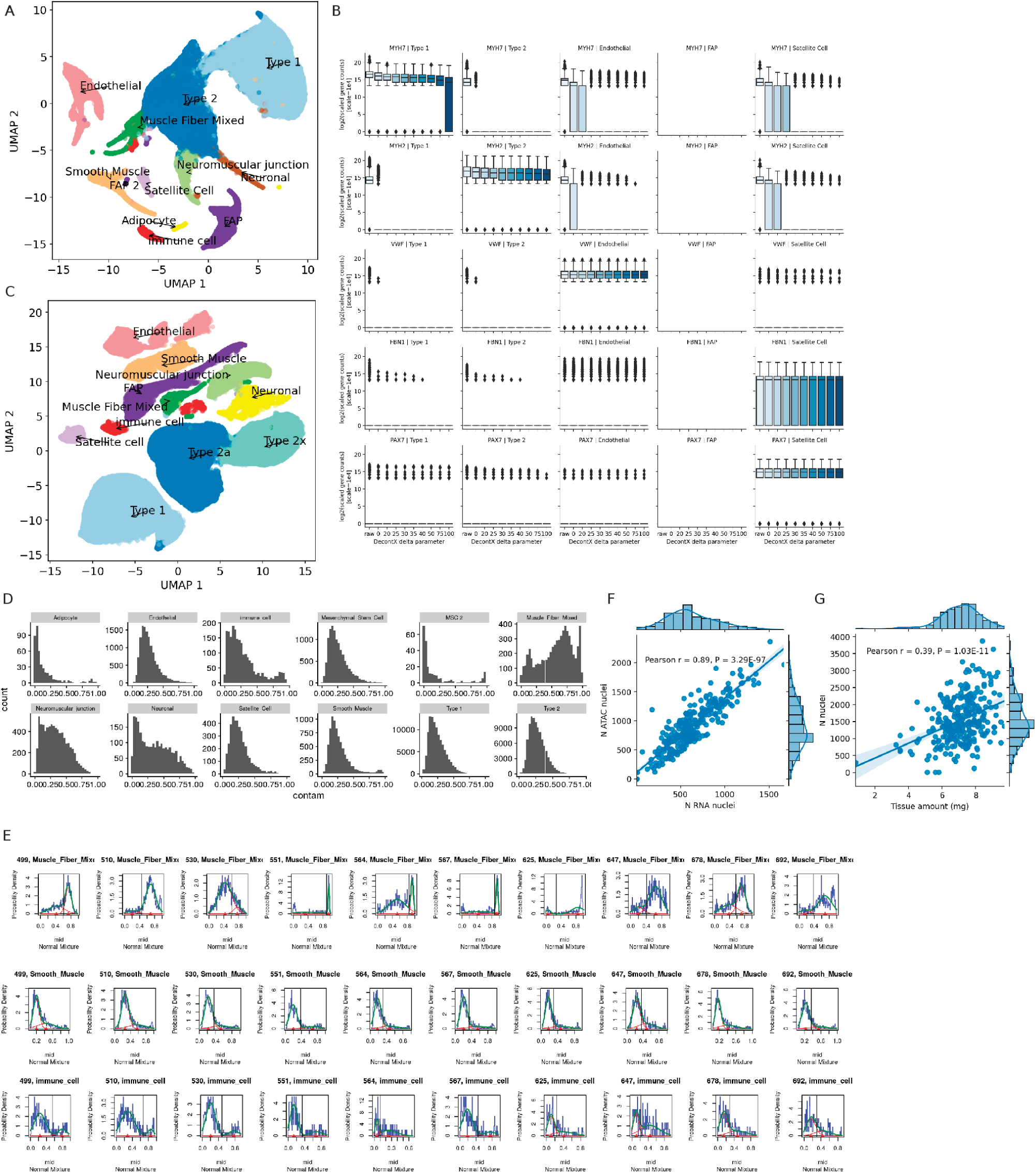
Adjusting for ambient RNA. (A) Initial RNA clustering using RNA counts unadjusted for ambient RNA (B) Marker gene counts across clusters without ambient RNA adjustment (raw) and after adjustment using DecontX run with a various delta parameter (prior for the ambient RNA counts) values) (C) Post-decontX snRNA-seq modality clustered jointly with snATAC-seq to obtain better optimized cluster labels. (D) Estimated ambient RNA fraction from DecontX across cluster labels (E) Batch and cluster-specific threshold for immune cell, muscle fiber mixed and smooth muscle clusters to further QC out nuclei due to high estimated ambient RNA fraction. For all other clusters, this threshold was set to 0.8 (F) After all stages of QC, the number of pass-QC RNA nuclei are correlated with the number of pass-QC ATAC nuclei per sample. (G) After all stages of QC, total pass-QC nuclei are correlated with the sample weights during nuclei isolation.

**Figure S4:**
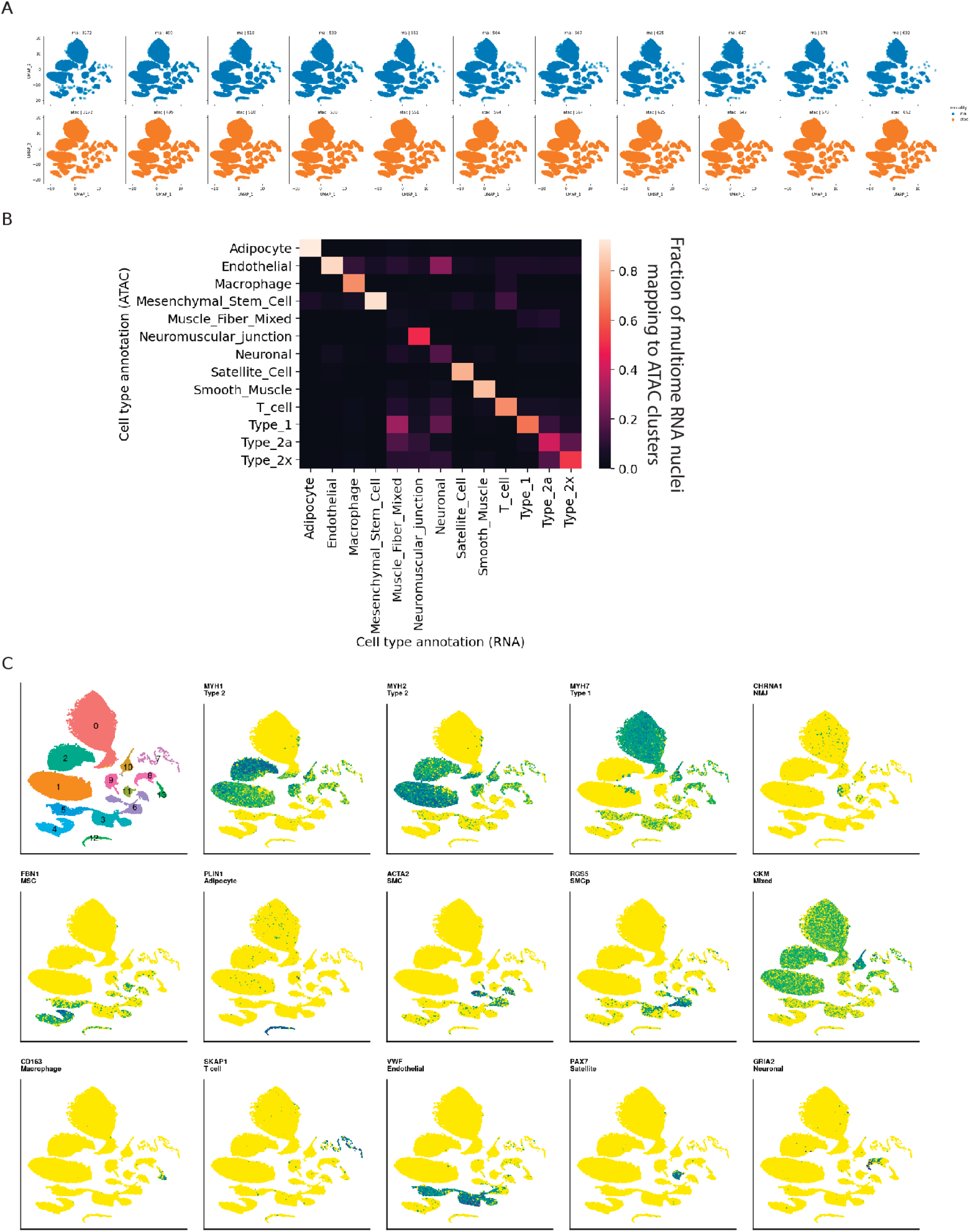
Joint clustering of the snRNA-seq and snATAC-seq modalities identified 13 cell-type clusters. (A) UMAP plots by batch and modality (B) Concordance between cluster annotations for the RNA and ATAC modalities of the multiome nuclei. Plotted are the fraction of nuclei in the RNA cluster that are assigned each annotation in the ATAC cluster. (C) UMAP plots showing cluster assignments and snRNA-seq expression of known marker genes

**Figure S5:**
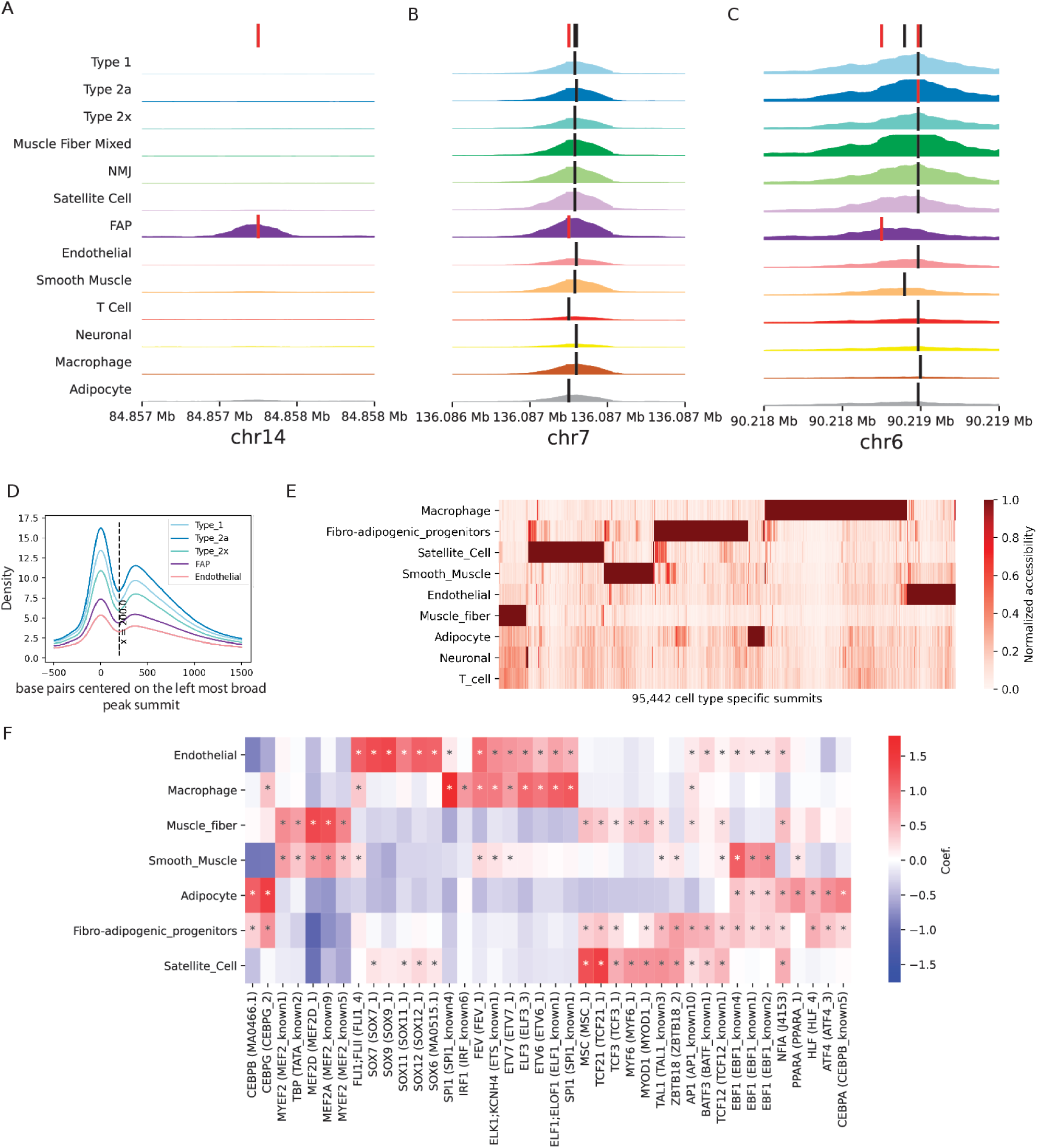
Identifying snATAC-seq peak features in clusters. (A) (B) (C) show example genomic locations showing identified consensus summit(s) (red) among all nearby summit calls (black), shown together in the top track and on thre respective cluster ATAC-seq signal tracks. (D) Aggregated ATAC-seq signal across all broad peaks in a cluster while centering on the left-most summit in the peak. (E) Heatmap showing peak summits identified as cluster-specific (F) Motif enrichment in cluster-specific peak summits calculated using a logistic regression approach. ‘*’ indicates significant enrichment (5% FDR).

**Figure S6:**
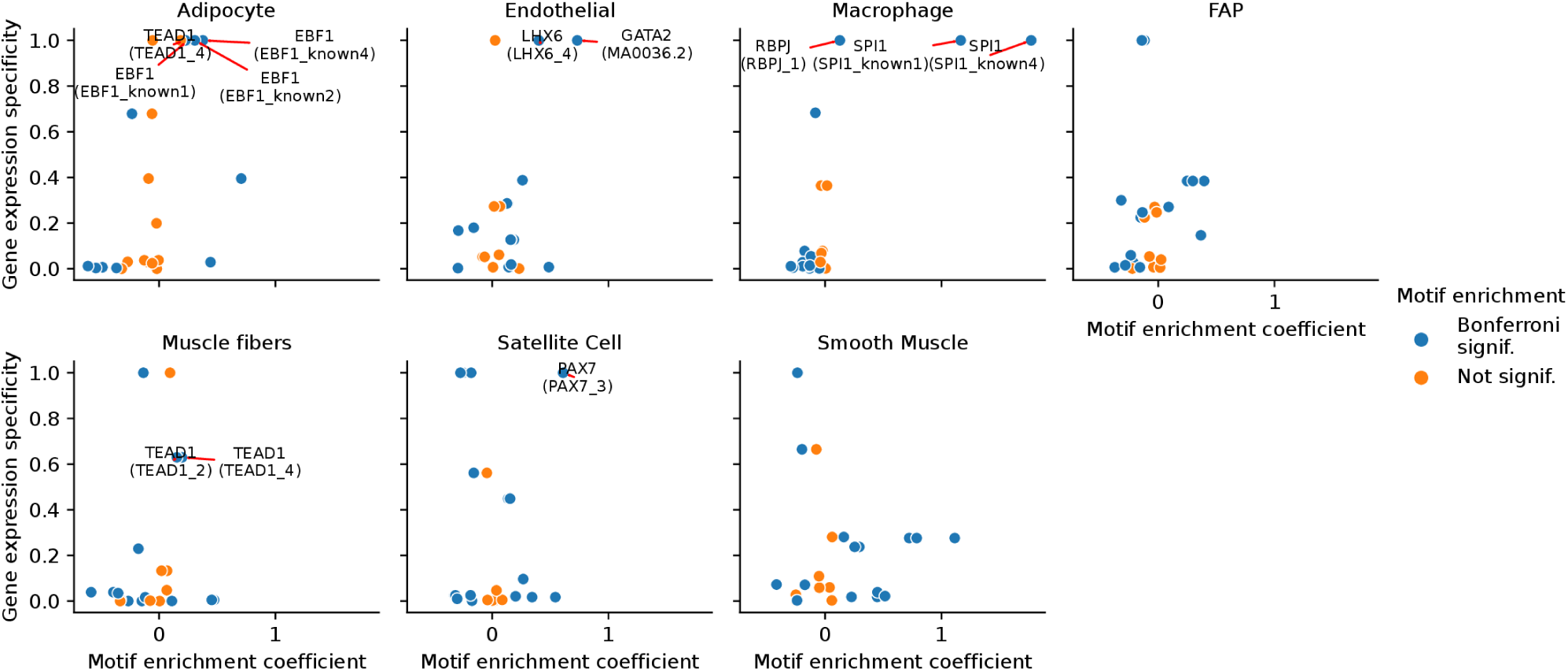
Cluster-specific expression of TF-genes drives cluster-specific motif enrichment. TF motif enrichment in cluster-specific peaks (regression coefficients) against the expression specificity scores in the for the corresponding TF gene in cell-types. Blue color indicates that the regression coefficient obtained a P-value lower than the Bonferroni correction threshold.

**Figure S7:**
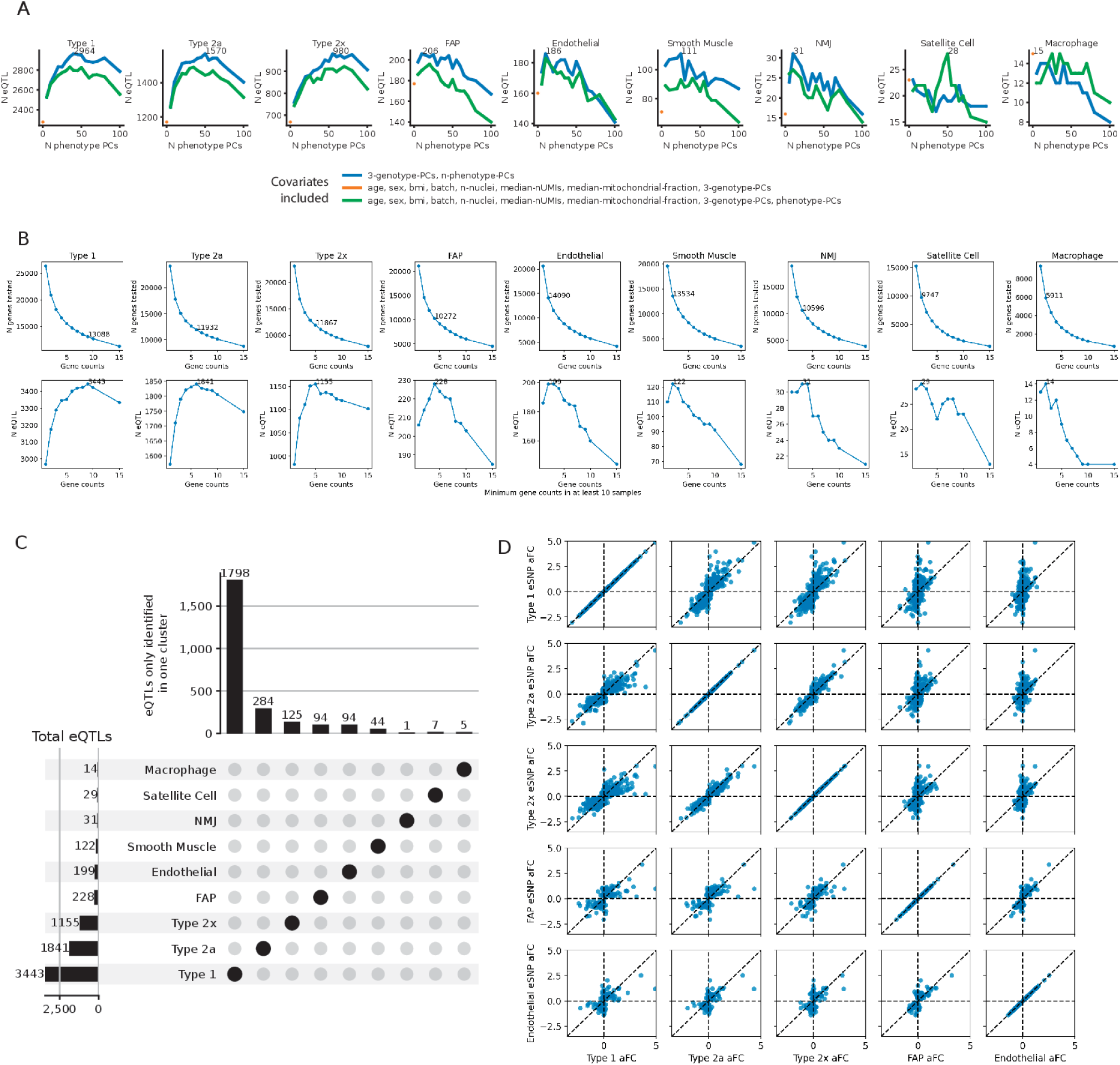
Identifying eQTL in clusters. (A) PC scan to maximize eQTL discovery. (B) Identifying testable genes with minimum n counts across at least 10 samples that maximize eQTL discovery. Number of testable genes and the number of eGenes (FDR 5%) at the selected minimum count threshold are labeled. (C) UpSet plot showing the total number of eGenes in each cluster and the number of genes identified in only one cluster. (D) eSNP allelic fold change (aFC) in clusters to compare eQTL effect sizes between pairs of clusters. Each facet shows aFC in both clusters for eSNPs identified in the cluster labeled on the y axis (5% FDR).

**Figure S8:**
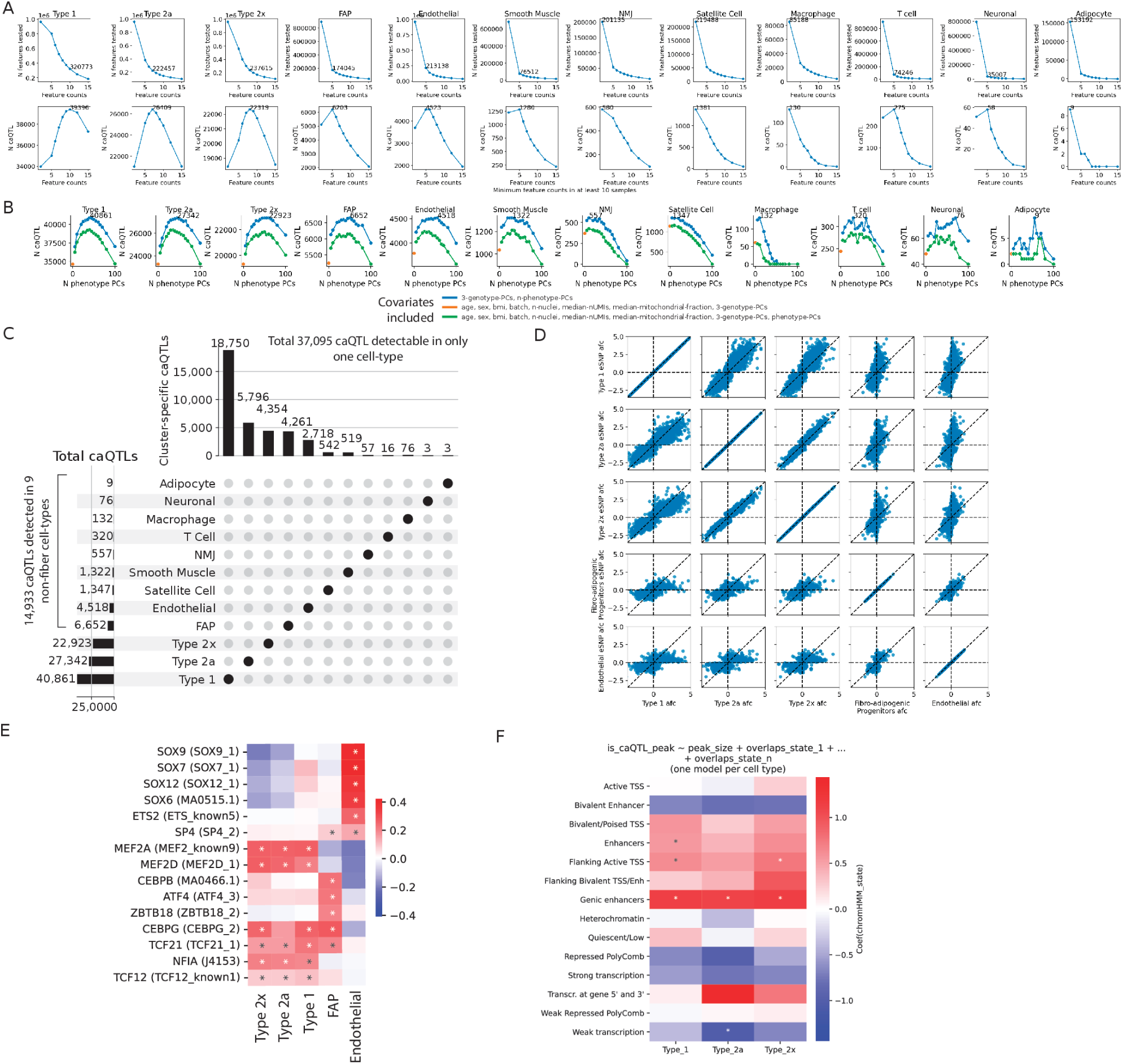
Identifying caQTL in clusters. (A) Identifying testable peak features with minimum n counts across at least 10 samples that maximize caQTL discovery. Number of testable peaks and the number of caPeaks (FDR 5%) at the selected minimum count threshold are labeled. (B) PC scan to maximize caQTL discovery. (C) UpSet plot showing the total number of caPeaks in each cluster and the number of caPeaks identified in only one cluster. (D) caSNP allelic fold change (aFC) in clusters to compare caQTL effect sizes between pairs of clusters. Each facet shows aFC in both clusters for caSNPs identified in the cluster labeled on the y axis (5% FDR). (E) Motif enrichment in caPeaks in five clusters. (F) Enrichment of ChromHMM states identified in bulk skeletal muscle to overlap with caPeaks in three muscle fiber clusters.

**Figure S9:**
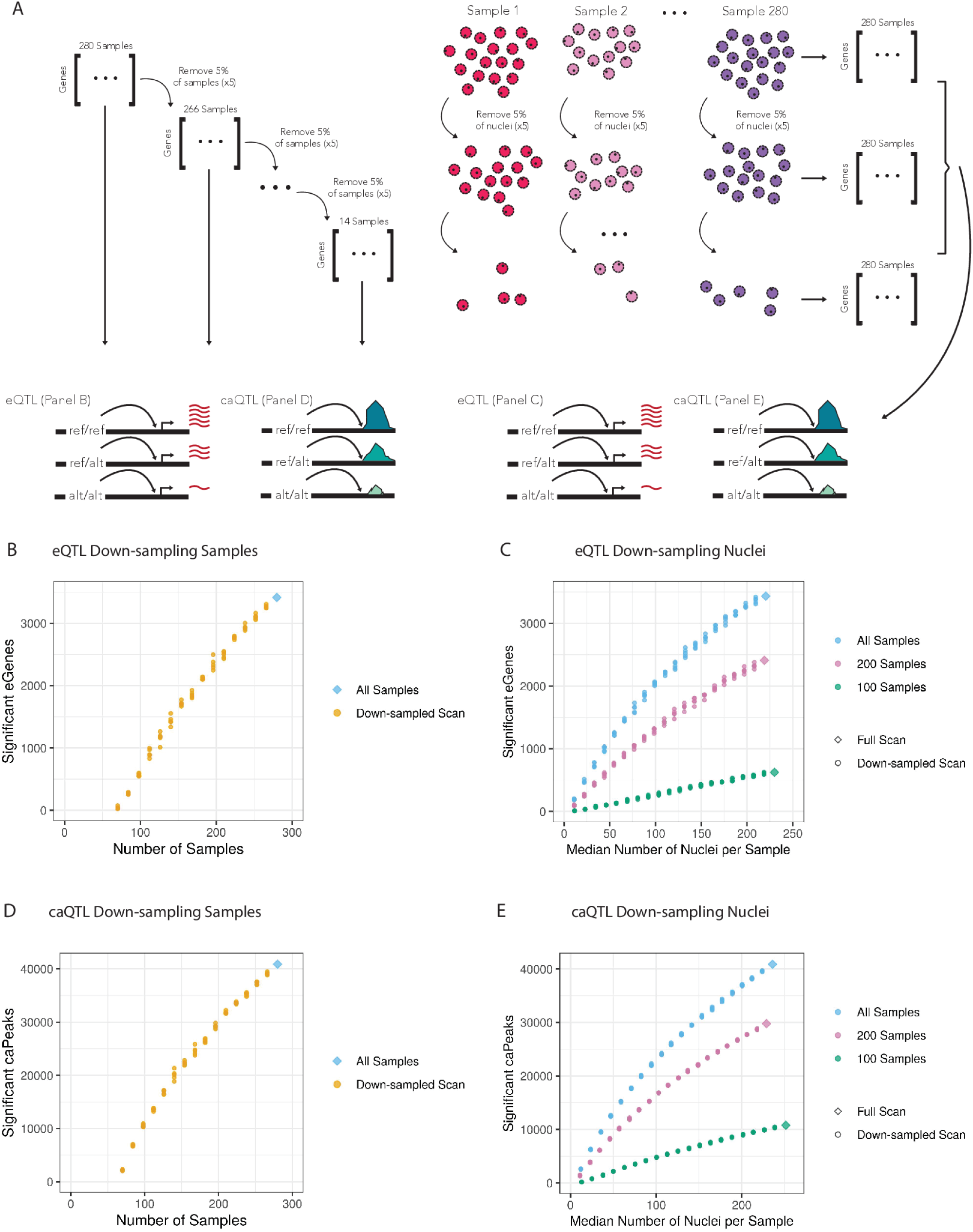
Down-sampling samples and nuclei in type 1 fibers. (A) Down-sampling strategy: either the number of samples (in 5% increments, left) or nuclei from each sample (right) were down-sampled followed by e/caQTL scan. Curves showing significant eGenes on down-sampling (B) samples and (C) nuclei in samples. Significant caPeaks on down-sampling (D) samples and (E) nuclei in samples.

**Figure S10:**
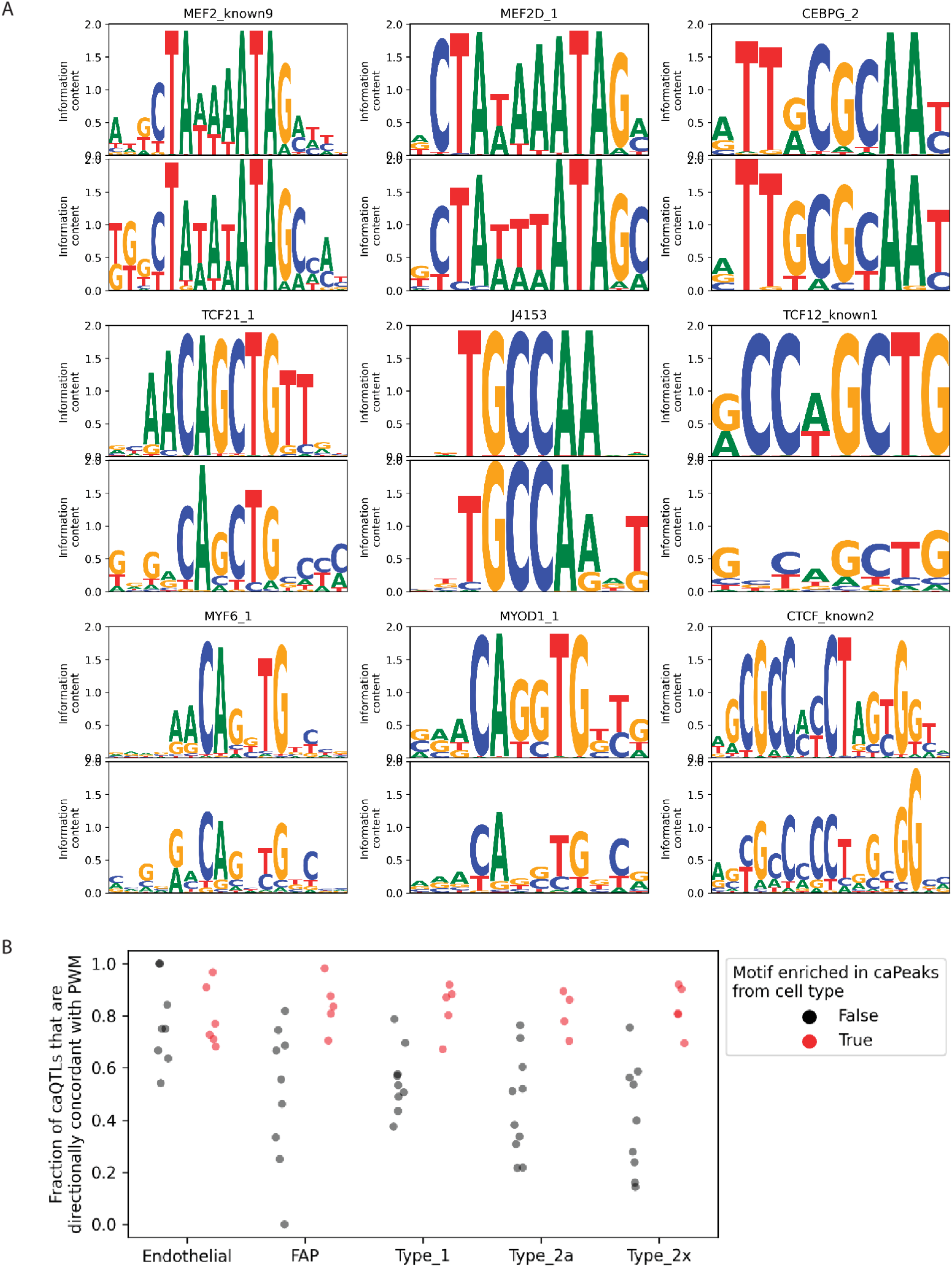
Motif reconstruction using caQTL data. Reconstruction for selected key motifs, including those enriched to occur in type 1 fiber caPeaks. Top row shows the canonical motif PWM, and the bottom row shows the reconstructed PWM. (B) Agreement between PWM motif scores (base preference in the motif) and QTL allele preferences.

**Figure S11:**
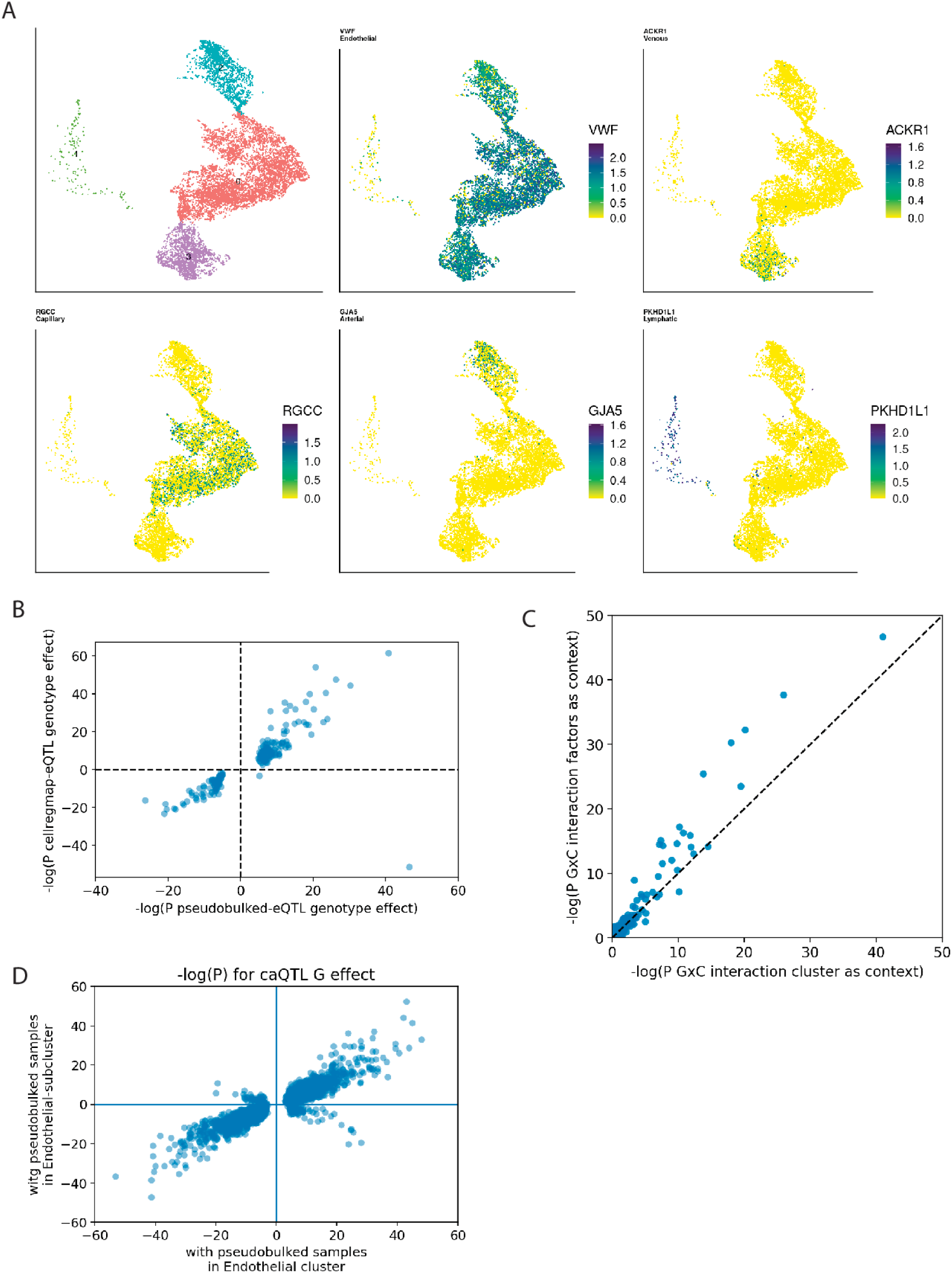
Endothelial state-specific e/caQTL using cellRegMap. (A) Joint (snRNA+snATAC) subclustering of endothelial nuclei identifies four subtypes/cell-states. snRNA nuclei UMAP plots show expression of key marker genes used to annotate the subclusters. (B) Top endothelial eSNP-eGene pairs identified from the initial pseudobulk analyses (5% FDR, **Fig 2A**) were tested for GxC interaction effect in CellRegmap. Scatter plot compares the signed -log10(P) of the additive genotype effect between the two eQTL models. (C) -log(P) for eQTL GxC interaction when using subcluster vs latent factors as the context. (D) Top endothelial caSNP-caPeak pairs identified from the initial pseudobulk analyses (5% FDR, **Fig 2B**) were tested for GxC interaction effect in CellRegmap. Scatter plot compares the signed -log10(P) of the additive genotype effect between the two caQTL models.

**Figure S12:**
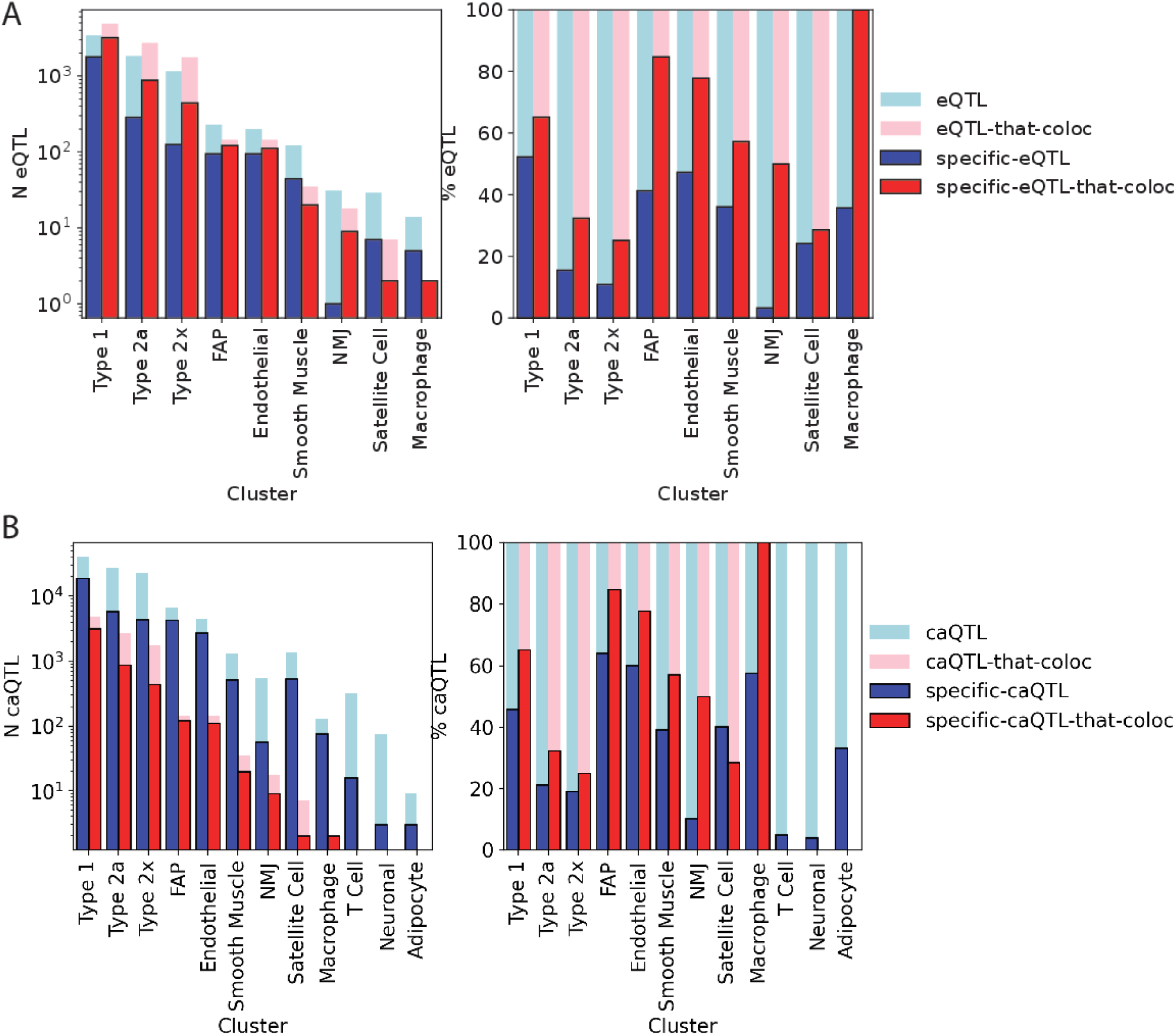
e-ca QTL colocalization. Total number of eQTL (left) and percentage of eQTL (right) that colocalize with caPeaks in each cluster, along with the number and percentage of eQTLs that are detected in only one cell type. (B) Total number of caQTL (left) and percentage of caQTL (right) that colocalize with eGenes in each cluster, along with the number and percentage of caQTLs that are detected in only one cell type.

**Figure S13:**
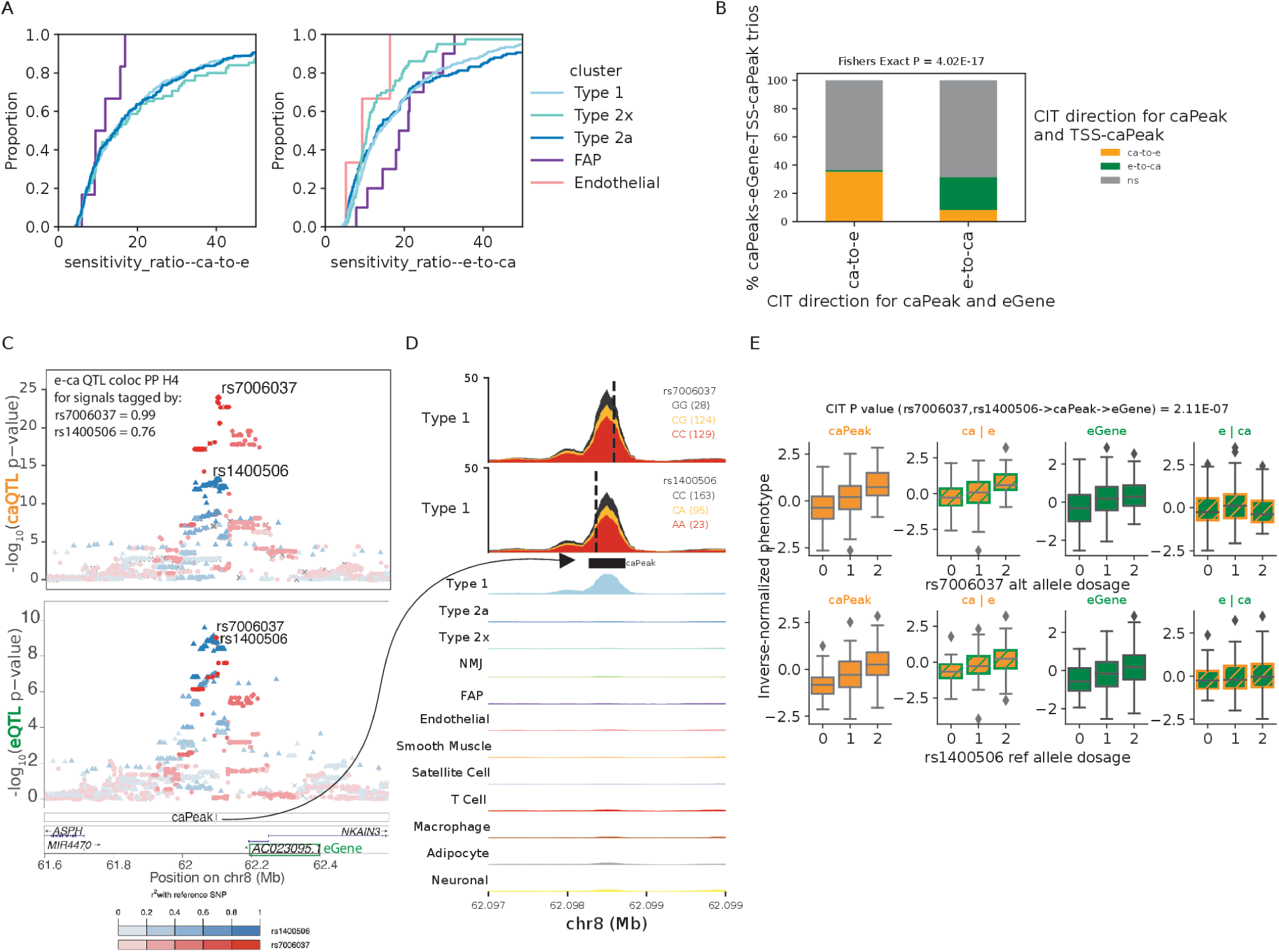
Causal inference test enables inferring causal direction between chromatin accessibility and gene expression. (A) Empirical cumulative distributions for MR Steiger directionality test sensitivity ratios for ca-to-e (top row) and e-to-ca (bottom row). Sensitivity ratio represents the estimated proportion of times the inferred direction flips over the bounds of measurement error in the exposure and outcome. Plots to the right show a zoomed-in view of the x axis. (B) CIT results between caPeak and TSS caPeak for ca-to-e or e-to-ca caPeak-eGene pairs. For caPeak and eGene pairs with significantly inferred causal direction ca-to-e or e-to-ca (x axis), where a caPeak was also identified in the TSS+1kb upstream region of the eGene, proportion of CIT outcomes between the distal caPeak and the TSS caPeak are denoted by the colors. Fisher’s exact test was performed after tallying all significant outcomes (5% FDR). (C) Example locus on chr8 where two independent eQTL signals identified for the lincRNA gene *AC023095.1* colocalize with two independent caQTL signals identified for a nearby caPeak in the type 1 cluster (Coloc PP H4 0.99, 0.76). The lead SNPs for the two signals rs7006037 and rs1400506 are labeled and the colors depict LD r^2^ relative to these variants. (D) snATAC-seq profiles in the type 1 cluster over the caPeak shown in **d** aggregated by the signal lead variant genotype classes. (E) Determined causal direction between the eGene-caPeak pair from **d** using the independent lead variants as instrument variables. Boxplots show inverse normalized chromatin accessibility, chromatin accessibility after regressing out gene expression, gene expression and gene expression after regressing out chromatin accessibility relative to the alternate allele dosages for the two lead variants rs7006037 (top) and rs1400506 (bottom).

**Figure S14:**
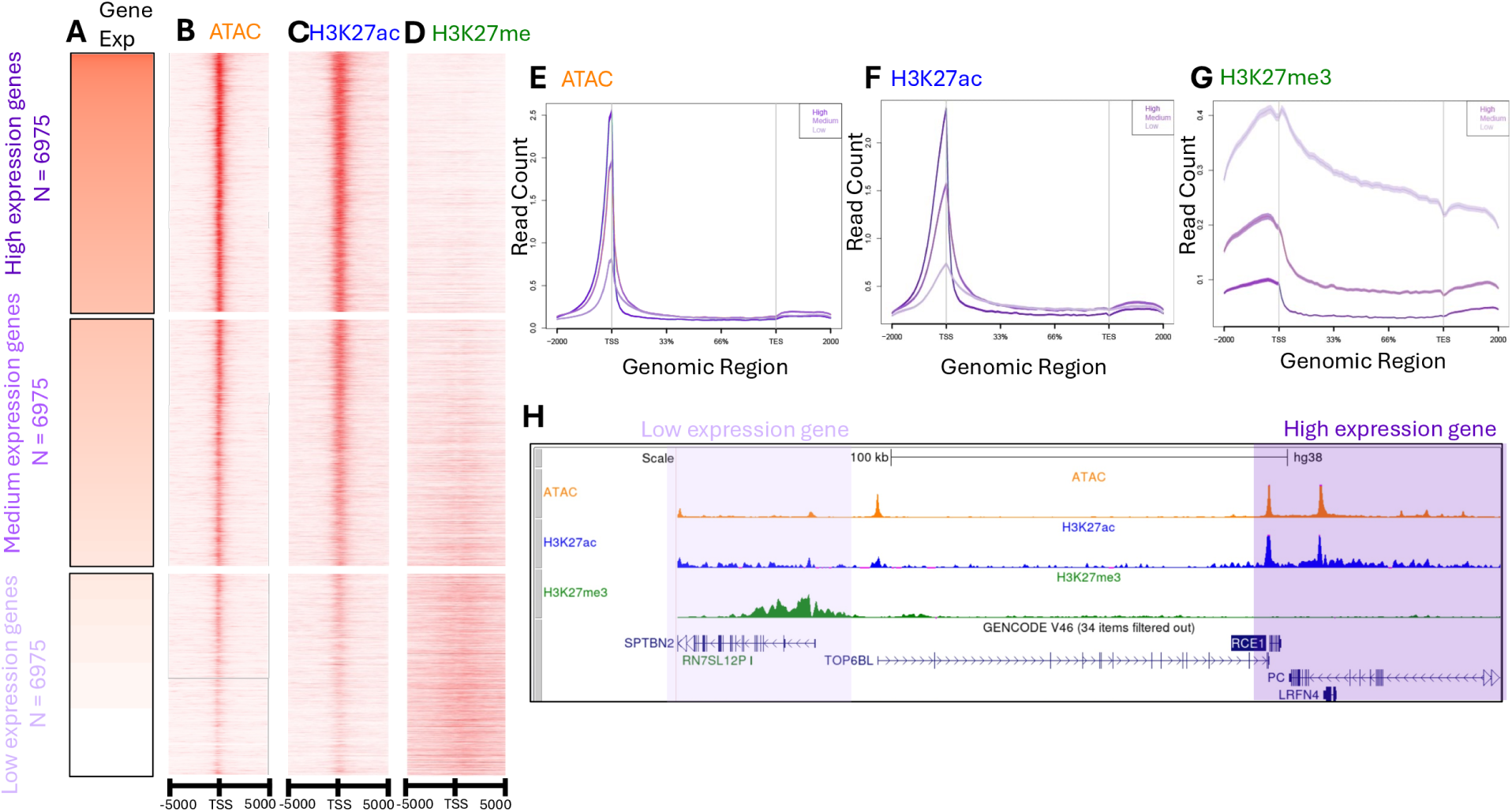
Bulk-skeletal muscle CUT&Tag profiles identify regulatory elements. (A) Genes in skeletal muscle sorted and binned by their expression levels. Darker red indicates higher expression. Panels B-D show reads at the TSS sites of the corresponding genes for (B) ATAC-seq (C) H3K27ac CUT&Tag, and (D) H3K27me3 CUT&Tag. (E) ATAC-seq (F) H3K27ac CUT&Tag, and (G) H3K27me3 CUT&Tag read-pileups over the gene bodies for the sets of genes with low, medium and high expression levels as described in (A). All genes are scaled to align the transcript start and end sites (TSS, TES). (H) UCSC browser session highlighting a repressed gene *SPTBN2* and highly expressed genes *RCE1* and *LRFN4* showing ATAC-seq, H3K27ac, and H3K27me3 tracks in skeletal muscle.

**Figure S15:**
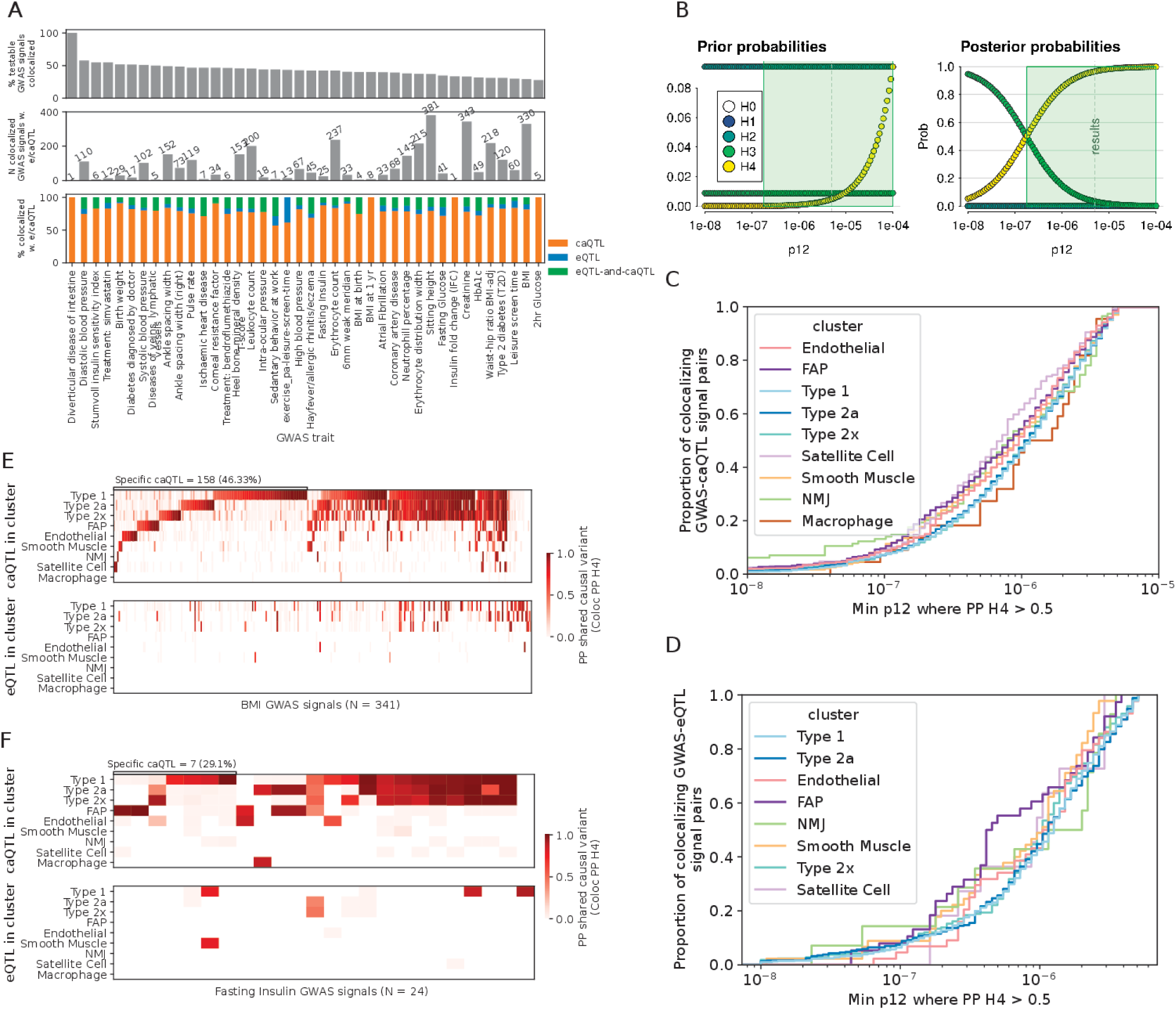
GWAS-caQTL colocalization and sensitivity. (A) Full GWAS-QTL colocalization summary showing the % and number of GWAS signals for each considered trait that colocalize with e/caQTL. (B) Sensitivity analysis for colocalization between T2D GWAS and a Type 1 caQTL at the C2CD4A/B locus comparing prior (left) and posterior (right) probabilities of each hypothesis over a range of prior probability values that any random SNP in the tested region is associated with both traits (p12). The Default p12 in coloc is 5e-6. Green boxes marks the set of p12 values at which PP H4 *>* 0.5. The lower the minimum p12 at which PP H4 *>* 0.5, the more robust the colocalization. (C) Empirical cumulative distribution of coloc sensitivity represented as the minimum p12 where PP H4 *>* 0.5 for GWAS-caQTL colocalization across the 40 GWAS traits considered, colored by cluster (D) Empirical cumulative distribution of coloc sensitivity represented as the minimum p12 where PP H4 *>* 0.5 for GWAS-eQTL colocalizations across the 40 GWAS traits considered, colored by cluster. For both B and C, all GWAS-QTL pairs observed colocalized (PP H4*>*0.5) at the default p12 of 5e-6 were considered. Heatmaps showing the coloc PP H4 for (E) BMI and (F) Fasting Insulin GWAS loci that colocalize with e/ca QTL across the five clusters.

**Figure S16:**
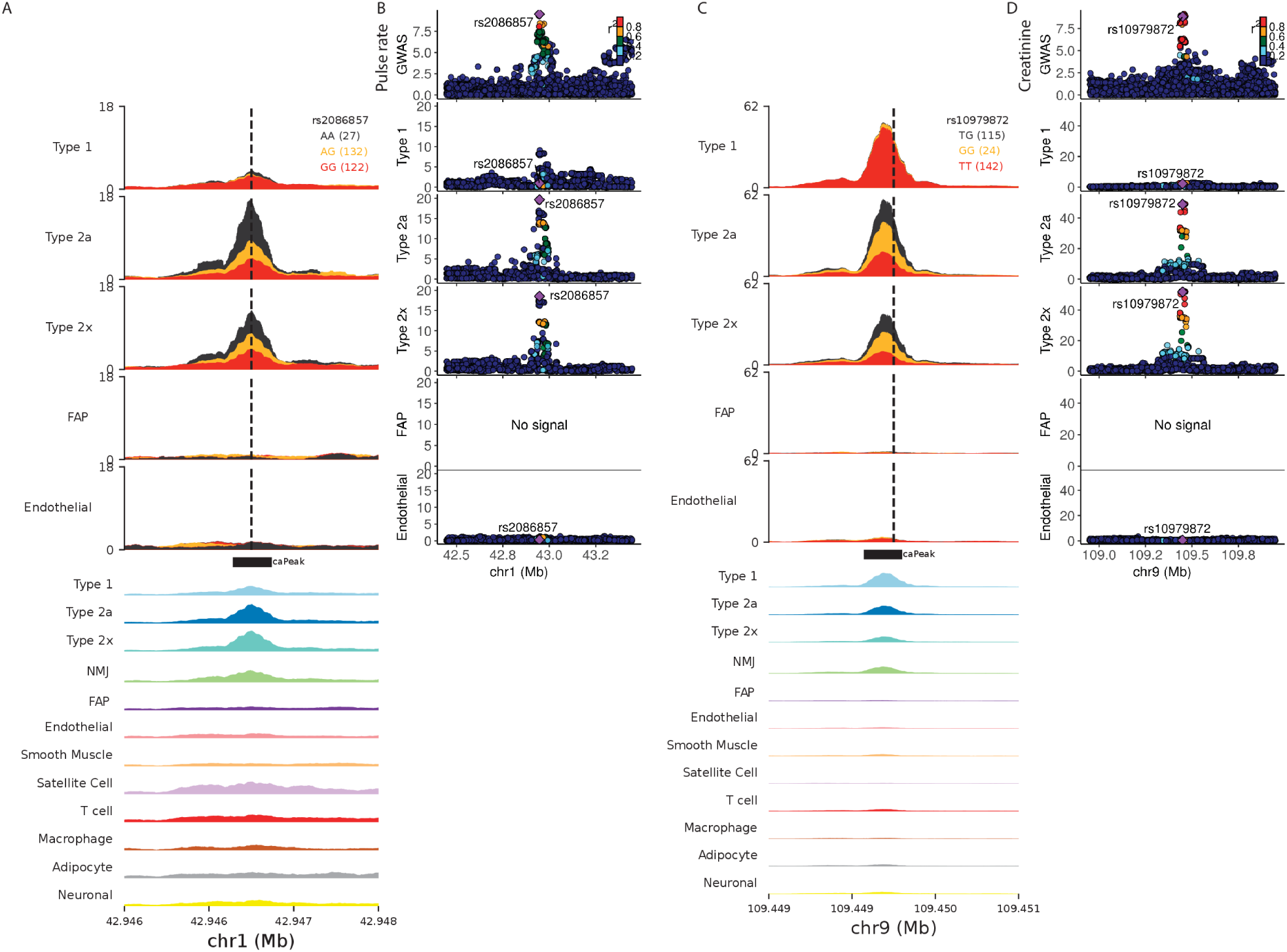
Identifying caQTL specific to individual muscle fiber types. Selected examples of caQTLs specifically identified in type 2 fibers (snATAC signal aggregated by caSNP genotype class in A, C) that colocalize with GWAS signals (locus zoom plots showing the GWAS signal and caQTL signals in five clusters in B, D)

**Figure S17:**
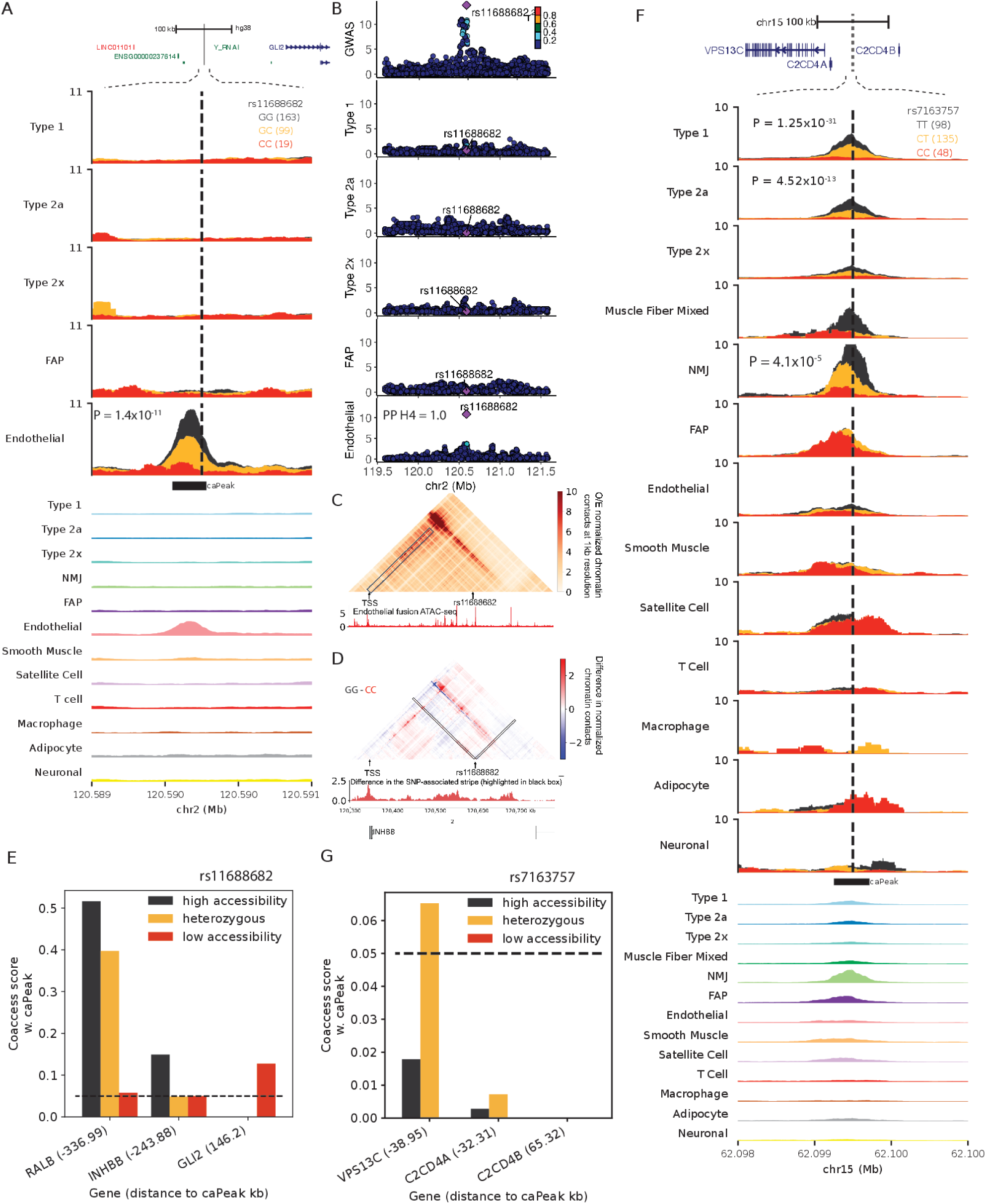
Integrating e/ca QTL signals with GWAS inform disease/trait relevant regulatory mechanisms. (A) *GLI2* genomic locus where a T2D GWAS signal is colocalized is an endothelial caQTL. snATAC profiles in five clusters by caSNP rs11688682 genotype in its *±*1kb neighborhood followed by aggregate profiles in 13 clusters. (B) Locuszoom plots for the *GLI2* GWAS signal (top) followed by the caQTL signal in the five clusters. The peak was not testable for caQTL in the type 1, 2a, 2x and FAP clusters due to low counts. (C) EPCOT-imputed micro-C chromatin contacts using endothelial ATAC data at 1kb resolution at the 500kb neighborhood centered at rs11688682. (D) Difference in the predicted normalized chromatin contacts using endothelial snATAC-seq from samples with the high (GG) and low (CC) accessibility genotype rs11688682. Interactions with rs11688682 highlighted in black are shown as a signal track below. (E) Endothelial chromatin co-accessibility scores between the GLI2 caPeak and TSS peaks of neighboring genes, classified by the caSNP genotype. Distance between the peaks is noted in parentheses. (F) C2CD4A/B genomic locus, followed by snATAC-seq profiles by caSNP rs7163757 genotypes in clusters, followed by aggregate snATAC profiles. (G) NMJ chromatin co-accessibility scores between the caPeak and TSS-peaks classified by caSNP genotype. Distance between the peaks is noted in parentheses.

### Supplementary Note

#### 4.32.1 Singlet identification and sample demultiplexing

In our study design, we multiplexed 40/41 samples in each batch and demultiplexed using known sample genotypes. Demultiplexing 41 samples per batch is well within the capability of assigning droplets to individuals, as reported by the original Demuxlet paper^107^. To demonstrate the high-quality donor assignments in our experiments, we ran Demuxlet on Batch 1 that included 40 samples with a mixture of “correct” (from batch 1) and “wrong” (from other batches) samples in the input VCF. As expected, the number of singlets identified by Demuxlet decreased linearly with the number of “wrong” samples in the VCF (**Figure SI1A**), with 0 singlets identified when all “wrong” 40 samples were provided in the VCF. None of the droplets are assigned as singlets to the “wrong” samples (**Figure SI1B**). Therefore, all samples are correctly assigned, even when incorrect samples are provided in the VCF, indicating that our QC process has resulted in high-quality nuclei with correct sample assignments.

**Figure SI1:**
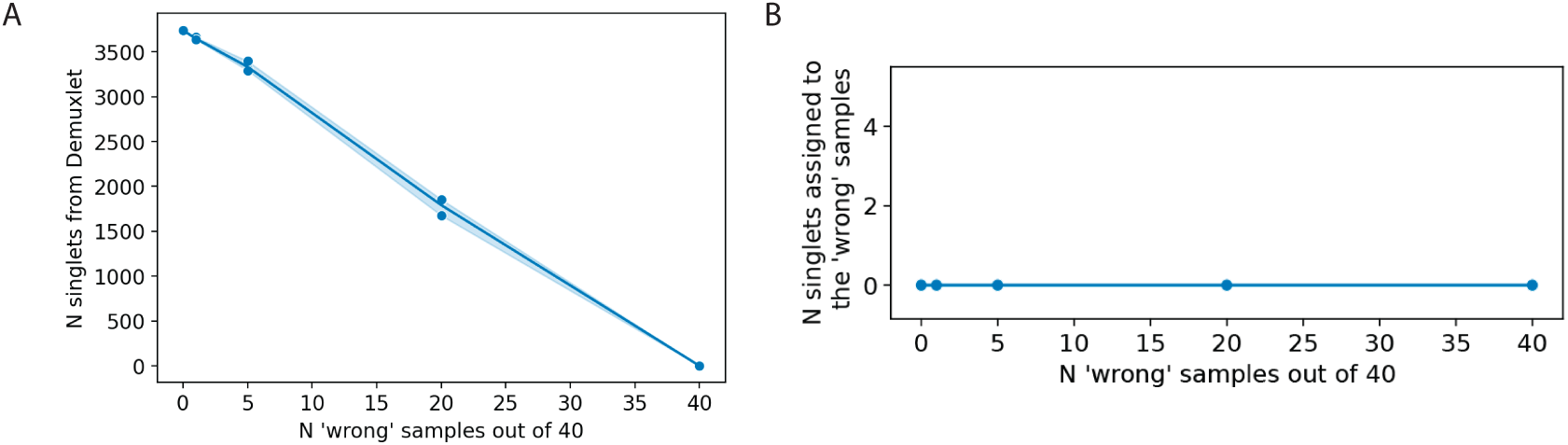
Demuxlet QC. (A) The number of correct singlets identified in the 40 sample batch decreases linearly as a function of the number of wrong samples supplied in the VCF. (B) Zero wrong samples are assigned, even when 40 wrong VCF entries are provided for genetic demultiplexing. Note the blue points at y=0 for the different ranges we tested.

#### 4.32.2 Ambient RNA and eQTL scans

While ambient transcripts are inevitable in nuclei preps, especially from processing of complex frozen tissue such as our muscle samples, genetic demultiplexing offers an improved strategy to identify clean droplets. The protocols and approaches to adjust for ambient transcripts are an active area of research. We thoroughly optimized our ambient RNA detection and correction (**Figure S3**). After this correction, a low level of muscle fiber marker gene expression still remained in clusters (**Figure 1E**). We reasoned that our eQTL scans are protected from ambient biases because Given our assay design of 40-41 multiplexed samples in each batch, the droplets with high ambient RNA are much more likely to be identified as doublets rather than being mislabeled. Because samples are pooled together, any remaining background ambient signal will not be associated with genetic variation and instead will represent a random mix of the samples in the pool.

To confirm our multiplexed study design protects us from spurious eQTL associations associated with ambient RNA levels, we ran eQTL scans with gene quantifications done before and after ambient RNA correction. **Figure SI2** shows that the eQTL p-values and slope direction for top snp-gene pairs are nearly identical both before and after ambient RNA correction. Thus, our eQTL scan results are not meaningfully influenced by ambient RNA.

#### 4.32.3 Clustering and QC

We integrated 287 FUSION snRNA+snATAC samples plus one multiome sample which included 456k nuclei spread across 10 batches plus one multiome batch. We performed the integration and clustering using Liger’s online iNMF algorithm which is capable to handle such large datasets. We factorized the RNA modality first since we found the RNA-only clusters to be more distinct and easily interpretable using known marker genes, then projected ATAC nuclei onto the factorization. We used the known multiome RNA-ATAC mapping to compare how concordant the cluster assignments were between the RNA and ATAC modality. We found that 82.8% of the non-muscle fiber multiome nuclei had the same RNA and ATAC cluster assignments. While most clusters were concordantly annotated between RNA and ATAC, the most frequent discrepancies were between the mixed fiber nuclei in RNA annotated type 1 or 2a in ATAC; or neuronal nuclei in RNA annotated type 1 or endothelial in ATAC (**Figure S4B**). In comparison, current next largest study^48^ integrated 73 brain snATAC+snRNA (on different nuclei) samples along with 19 multiome samples and achieved 79.5% - 85% concordance. The widely used Seurat program [134], obtained a 90% concordance on 12,000 blood cell nuclei ([vignette here](https://satijalab.org/seurat/articles/seurat5_atacseq_integration_vignette)) from a much smaller dataset that was easier to analyze because it was not from a solid complex tissue, and all nuclei were from one batch of 10X multiome assay rather than 456k nuclei spread across 10 batches plus one small multiome batch. These observations help put the performance of our clustering approach into perspective.

**Figure SI2:**
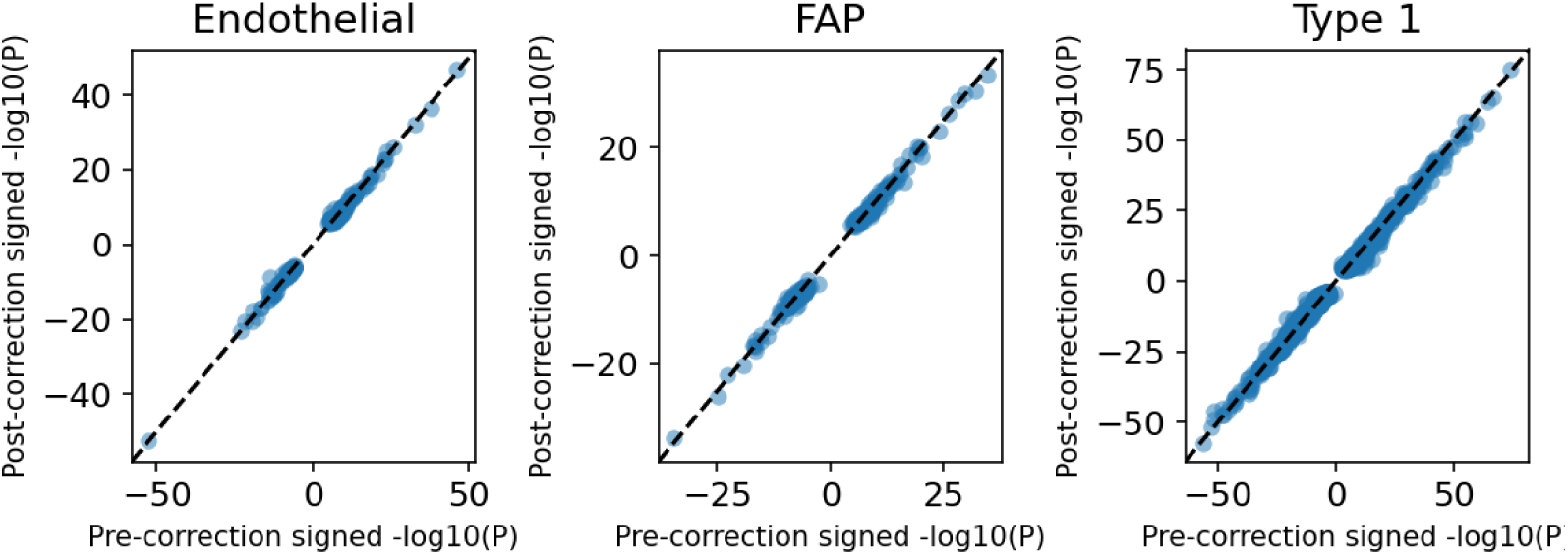
eQTL scan before and after ambient RNA adjustment. For top SNP-gene pairs (5% FDR) in endothelial, FAP, and type 1 clusters from the eQTL scans done post ambient-RNA correction, corresponding [–log10(P value) x sign of the slope] are plotted from an eQTL scan done pre-ambient-RNA correction. The striking similarity pre and post ambient-RNA correction shows that the background signal is not associated with genetic variation.

The fraction of nuclei assigned to each cluster within the RNA and ATAC modalities varied more for some clusters than others. For example, T cell cluster constituted 0.14% of RNA nuclei, but 5.15% of ATAC nuclei **Figure SI3**. These differences could be due to both technical and biological factors. For example, the ambient “soup” profiles for RNA vs ATAC are expectantly different. We considered droplets containing very low number of UMIs/HQAA as a representation of ambient profile and observed that the most highly expressed genes in the snRNA soup were muscle fiber genes, which is the most abundant cell type. Whereas, most snATAC soup reads mapped to the mitochondrial genome, which are all removed during analysis. Second, chromatin and transcription programs in a cell could manifest intrinsic cell-state differences. It has been demonstrated that chromatin accessibility information from snATAC-seq provides a coarser-grained representation of cell-states compared to transcription information from snRNA-seq profiling, which suggests that cells could retain a primed or permissive chromatin landscape that can allow dynamic state transitions in response to relevant conditions^48,88^.

While we performed extensive QC, ambient RNA adjustment and joint clustering and obtained meaningful clusters, the “muscle fiber mixed” cluster showed higher ratio of exonic reads vs reads over the entire gene body in some batches (**Figure SI4A**) and showed elevated fraction of mitochondrial reads (**Figure SI4B**). This suggests that the muscle fiber mixed cluster contained nuclei with relative higher ambient RNA, and likely represented technical variation in nuclei extraction efficiency across some batches. We account for technical variation due to batch in subsequent analyses.

**Figure SI3:**
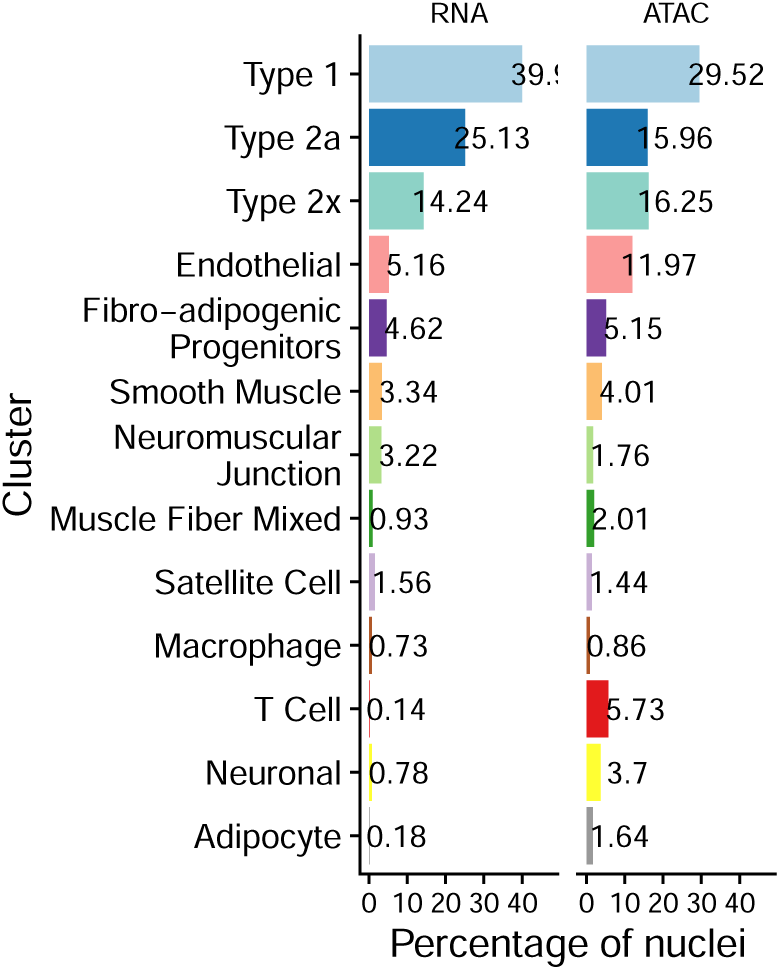
Cluster abundance by modality. Percentage of nuclei in each modality assigned to each cluster.

#### 4.32.4 Motif reconstruction

We observed several examples of TF motifs that reconstructed well using caQTL allele preferences and effect sizes (**Figure 2G**, **Figure S10**). Due to the sparsity in motif-caSNP overlaps, it was impractical to comprehensively quantify the total number of correctly reconstructed motifs out of all tested and compare across cell-types. In type 1 fibers, ∼66% of the 540 PWMs had at least one overlap at each position of the PWM, however in endothelial cells and FAPs, where a lower number of caQTLs were identified, the corresponding percentages are 15% and 19%, respectively. Nevertheless, concordance between TF and cell type is evident. For example, the MEF2 known9 motif, which is enriched in muscle fibers (**Figure S8E**), reconstructed well in Type 1 fibers (**Figure SI5A**); however in endothelial cells, even most high information content MEF2 known9 positions don’t overlap a caSNP (or proxy) (**Figure SI5B**). In contrast, the SOX motifs (e.g., SOX7 1) were enriched in endothelial caPeaks but not muscle fiber caPeaks (**Figure S8E**). SOX7 1 motif reconstruction is sparse in endothelial cells; however, we still see the high information content positions in the core motif well reconstructed in endothelial cells, whereas the reconstruction in type 1 fiber does not capture this core motif as well (**Figures SI5C**–**SI5D**).

It is important to note that many PWMs are not actually expected to be well-reconstructed. The PWM reconstruction is expected to work well only when the original PWM corresponds to a TF expressed in that cell type and, more importantly, when the TF has a large impact on chromatin accessibility in that cell type (since the variants used for reconstruction are caQTLs). If a variant impacts binding of a TF but that TF does not have much impact on chromatin architecture, the variant is unlikely to be a caQTL in the first place. A PWM of a TF that does not impact chromatin architecture is likely to overlap a subset of caQTLs just by chance, so a reconstruction can sometimes be produced, but the reconstruction in that case is not expected to be reliable.

**Figure SI4:**
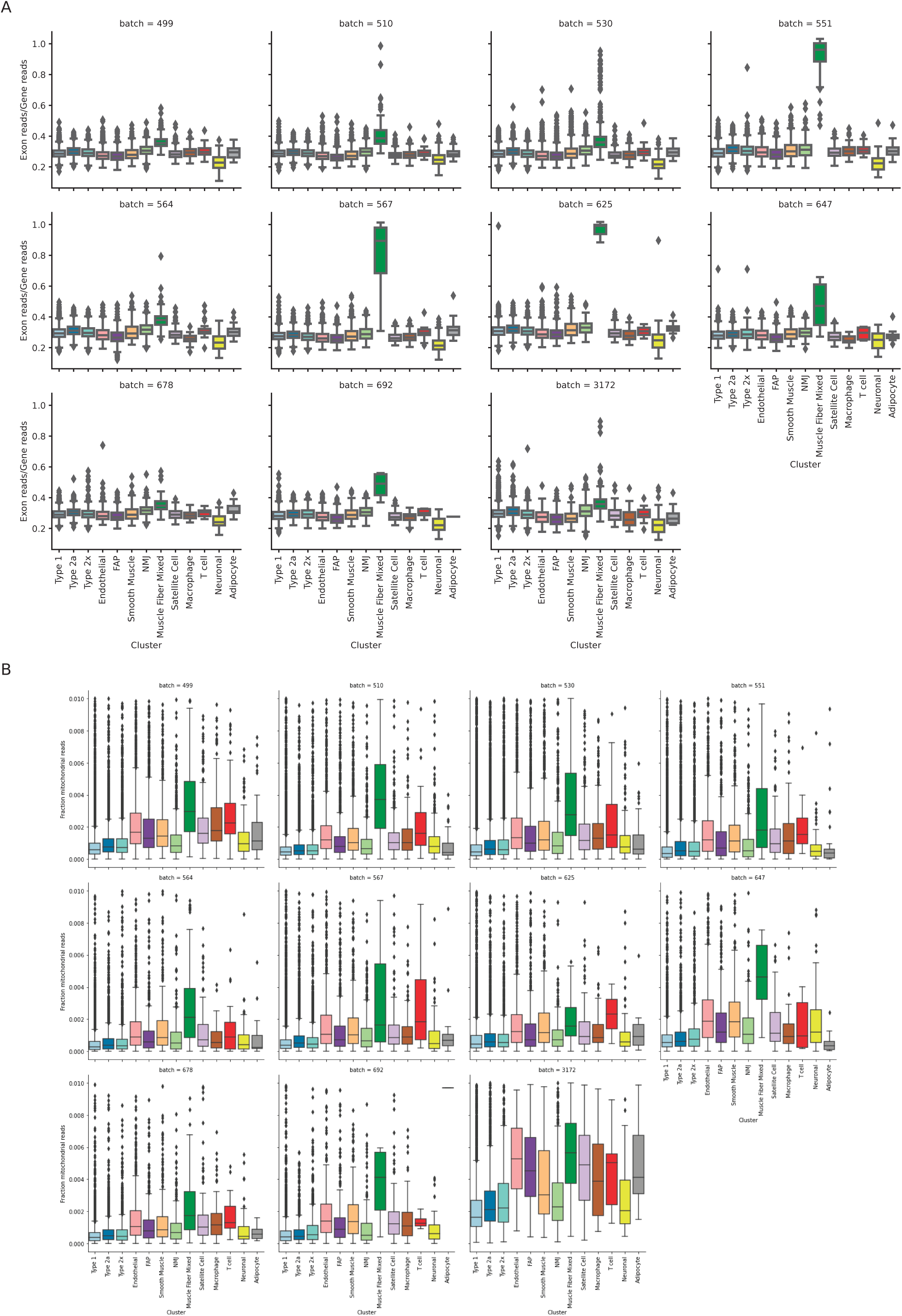
Joint clustering of the snRNA-seq and snATAC-seq modalities identified 13 cell-type clusters. (A) Fraction of exonic reads over gene reads in nuclei across clusters and batches (B) Fraction of mitochondrial reads in nuclei across clusters and batches

**Figure SI5:**
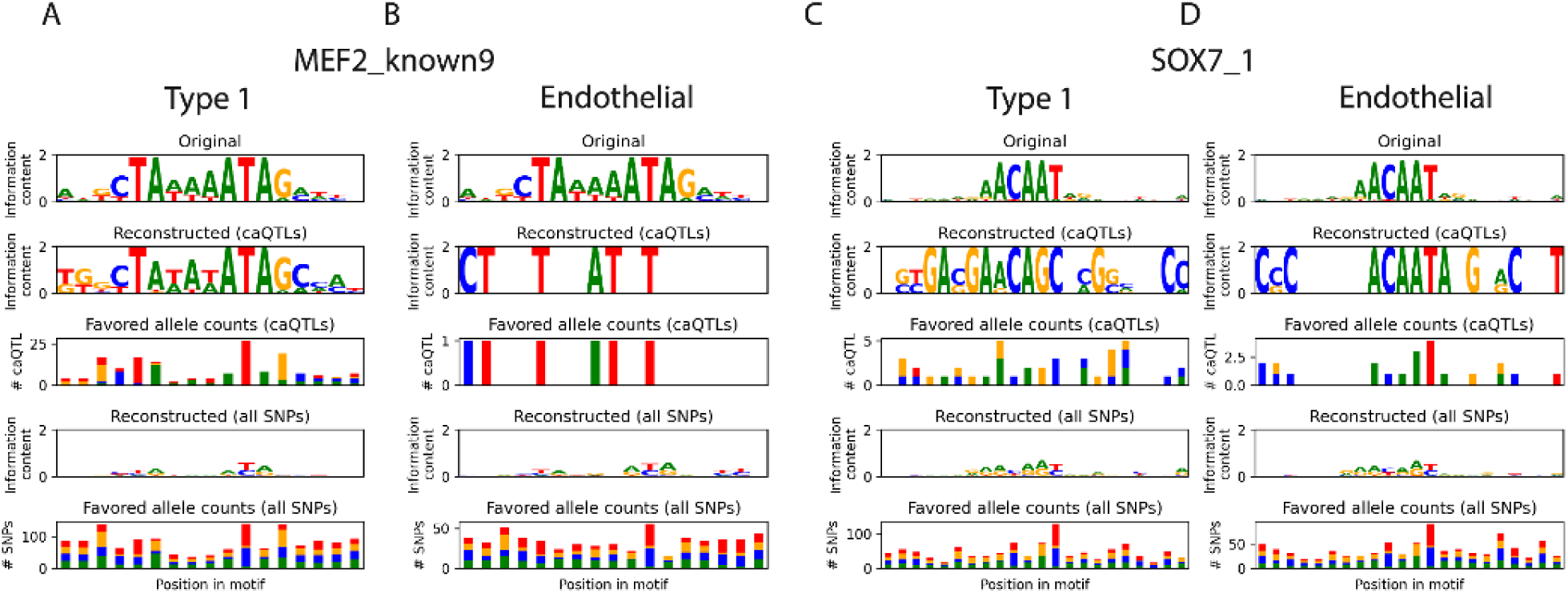
Reconstruction of selected motifs enriched in caPeaks in type 1 fibers vs endothelial. (A) MEF2 known9 motif reconstruction in type 1 fibers, and (B) endothelial. SOX7 1 motif reconstruction in (C) type 1 fibers, and (D) endothelial cell-type cluster.

### Supplementary Protocol S1

This protocol is designed for 4 samples. Please scale up or down based on the number of samples. All the steps have to be performed on ice or at 4°C. All the 2 mL and 1.5 mL tubes used in this protocol are Protein LOBIND tubes from EPPENDORF.

#### Materials

- CP02 cryoPREP automated dry pluverizer (Covaris 500001)
- Eppendorf thermomixer C (EP 5382000015)
- 2mL glass tissue grinder and pestle (Kimble chase, 885301-0002, 8853000002)
- 70 *µ*m strainer (Fisher 501457900)
- Celltrics, 20 *µ*m, 30 *µ*m cell strainer (Fisher NC9682496, Fisher NC9699018)
- Fisherbrand Sterile Plastic Culture (FACS) Tubes 149563C
- RNase Inhibitor (Thermofisher N8080119)
- Eppendorf Protein Lobind 1.5 mL tubes (Eppendorf 022431081)
- Eppendorf Protein Lobind 2.0 mL tubes (Eppendorf 022431102)

#### LB1 buffer

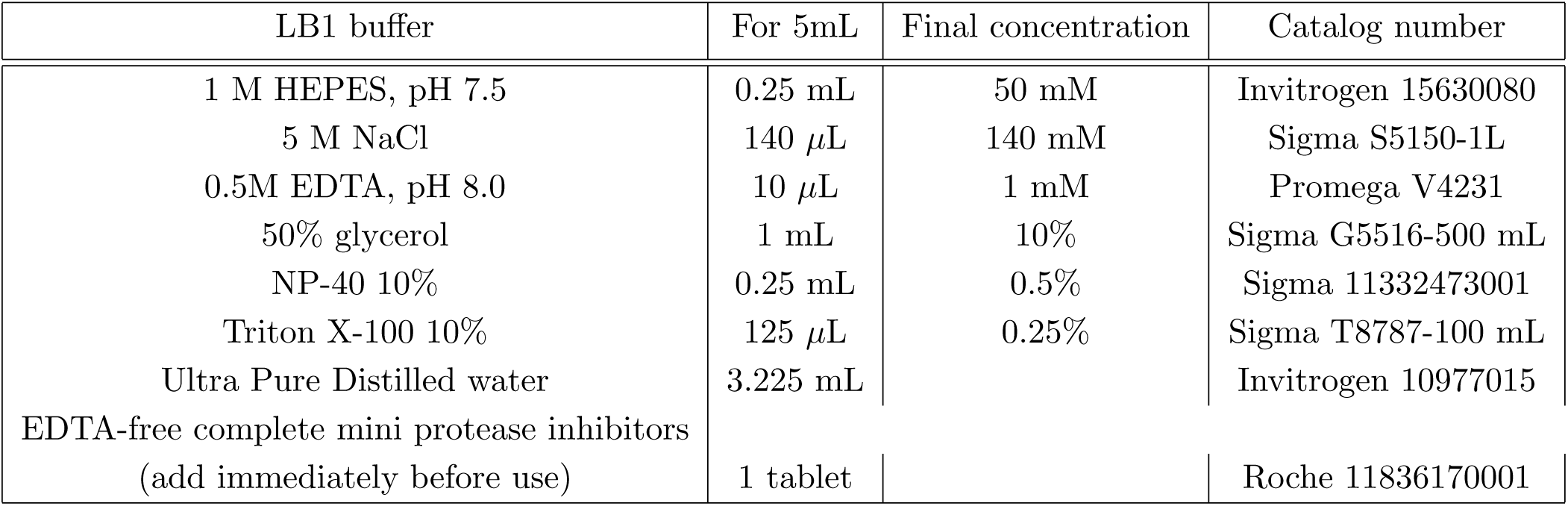

| LB1 buffer | For 5mL | Final concentration | Catalog number |
| --- | --- | --- | --- |
| 1 M HEPES, pH 7.5 | 0.25 mL | 50 mM | Invitrogen 15630080 |
| 5 M NaCl | 140 μL | 140 mM | Sigma S5150-1L |
| 0.5M EDTA, pH 8.0 | 10 μL | 1 mM | Promega V4231 |
| 50% glycerol | 1 mL | 10% | Sigma G5516-500 mL |
| NP-40 10% | 0.25 mL | 0.5% | Sigma 11332473001 |
| Triton X-100 10% | 125 μL | 0.25% | Sigma T8787-100 mL |
| Ultra Pure Distilled water | 3.225 mL |  | Invitrogen 10977015 |
| EDTA-free complete mini protease inhibitors<br>(add immediately before use) | 1 tablet |  | Roche 11836170001 |

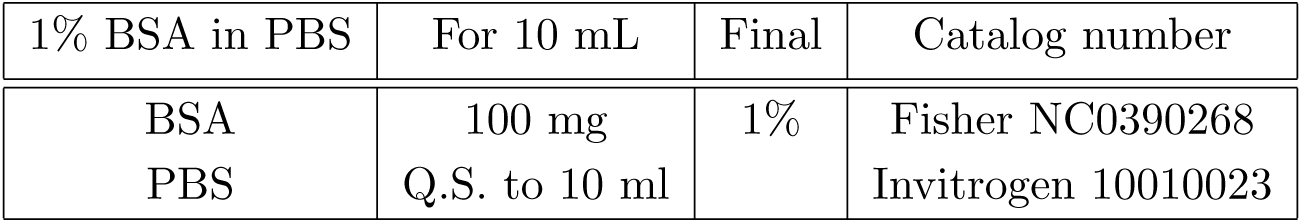

| 1% BSA in PBS | For 10 mL | Final | Catalog number |
| --- | --- | --- | --- |
| BSA | 100 mg | 1% | Fisher NC0390268 |
| PBS | Q.S. to 10 ml |  | Invitrogen 10010023 |

#### Nuclei extraction

1. Frozen tissue (40-90 mg) was pulverized into a fine powder while cold (dry ice and LN2) using an automated dry pulverizer CP02 cryoPREP.
2. Pulverized Frozen tissue (40-90 mg) was suspended in 1 mL of ice-cold 1x PBS in a 1.5 mL tube (Eppendorf 022431081) and centrifuged at 2000g for 3 min at 4°C. The supernatant was removed and the pellet was resuspended in 1 mL LB1.
3. The tissue was lysed by rocking the tubes in Eppendorf thermomixer C (EP 5382000015) at 4°C at 300 rpm for 5 min.
4. Each sample was transferred into a prechilled 2 mL glass Dounce homogenizer and homogenized with 10 strokes of loose pestle A, and 20 strokes of tight pestle B and then transferred to 1.5 mL tube and centrifuged at 2000g for 5 min at 4°C.
5. The supernatant was aspirated and the pellet was resuspended in 1 mL of ice-cold 1% BSA and centrifuged at 100g for 1 min at 4°C.
6. The supernatant was collected, discarding the loose debris pellet.
7. All the filters were prewet; 70 *µ*m, 30 *µ*m and 20 *µ*m filters, with 200 *µ*L of 1% BSA each and previously collected supernatant was sequentially filtered through 70 *µ*m, 30 *µ*m and 20 *µ*m filters respectively.
8. The supernatant was filtered through a 70 *µ*m strainer and the filtrate was collected into a 50 mL conical tube.
9. The collected filtrate was passed through a 30 *µ*m celltrix strainer and collected into a 2mL tube.
10. The collected filtrate was passed through a 20 *µ*m celltrix strainer and collected into a 2mL tube.
11. The filtrate was transferred to a 1.5mL tube and centrifuged at 350 x g for 10 min at 4°C (be careful of the tube direction). The supernatant was aspirated (the supernatant was saved as a precaution) with flexi tip gel loading tip (were very careful not to disturb the pellet) and the nuclei were resuspended in 500 *µ*L of 1% BSA in PBS.
12. The nuclei suspension was centrifuged again at 350 x g for 10 min at 4°C (be careful of the tube direction). The supernatant was aspirated very carefully (the supernatant was saved as a precaution) with flexi tip gel loading tip and the nuclei were resuspended in 100 *µ*L of 1%BSA in PBS.
13. The nuclei were counted with cell counter (Trypan blue stains nuclei; typically 4-9 *µ*m) and diluted appropriately for RNA and ATAC submissions.
14. (RNA Submission) To achieve the desired nuclei concentration, an appropriate amount of nuclei was diluted with 1% BSA in PBS. To this suspension, RNase inhibitor was added to get a final concentration of 0.2 U/*µ*L. The nuclei was counted and submitted for snRNA seq.
15. (ATAC Submission)-The rest of the nuclei was spun down at 350 x g for 10 min at 4°C (were careful of the tube direction). The supernatant was aspirated very carefully with flexi tip gel loading tip and the nuclei were resuspended in an appropriate volume of 1X diluted nuclei buffer (20x buffer supplied by 10X genomics). The nuclei was counted and submitted for snATAC seq.

### MPRA experiment and data analysis

#### Cloning

We designed oligos with a variant centered within 197 bp of flanking sequence (98 bp on each side). We also included negative control sequences selected from a previous publication (1).

Oligos were synthesized by IDT as 230 bp sequences containing the 197 bp sequences of interest flanked by two adapter sequences for cloning. We added 20 bp barcodes with additional adapters via a 5-cycle PCR reaction containing 13 ng oligo pool, 12.5 uL of NEB Q5 HotStart Hifi Mastermix, 1.25 uL of 10X SYBR Green I and 0.25 uM each of primers AT0003 and AT0039. This reaction was performed in quadruplicate with the following thermocycler conditions: 98*^◦^*C x 30 sec, [98*^◦^*C × 10 sec, 65*^◦^*C x 15 sec, 72*^◦^*C x 30 sec, plate read, 72*^◦^*C x 8 sec] x 5, 72*^◦^*C x 2 min. Barcoded oligos were diluted 1:25 in H2O, then amplified in a second PCR reaction using the same conditions with primer AT0050 in place of AT0039 and cycling for 14 cycles. We pooled replicate reactions, cleaned the final amplified, barcoded oligos with 1.8X SPRI beads, and eluted in 15 uL of LoTE.

We modified pMPRA1 (a gift from Tarjei Mikkelsen, Addgene #49349; 2) by adding in PaqCI cloning sites and an EGFP open reading frame. We cloned the barcoded oligos into this backbone (2:1 molar ratio of oligos:backbone) using PaqCI-mediated Golden Gate assembly per NEB recommendations. We incubated the assembly reaction at 37*^◦^*C for 1 hour then 65*^◦^*C for 5 min. We performed a secondary digest with 20 U SfiI to remove empty backbones, then cleaned with 0.8X SPRI beads and eluted in 10 uL of LoTE. We transformed 1 ul of the assembly into NEB 10-beta electrocompetent bacteria and expanded overnight in 150 mL of ampicillin-containing LB. In parallel, we plated serial dilutions and estimated a library complexity of ∼ 5 × 106 CFU.

We prepared sequencing libraries from the promoter-less MPRA plasmids to create an oligo-barcode pairing dictionary. In brief, we amplified the oligo-barcode region in a reaction containing 100 ng plasmid library, 20 uL 5X Kapa Fidelity Buffer, 3 uL Kapa dNTPs, 5 uL 10X SYBR Green I, 2 units Kapa HiFi HotStart DNA polymerase, and 0.5 uM each of primers jklab0343 and jklab0344. We indexed with standard Illumina primers and sequenced the library on a NovaSeq 6000.

To create the final plasmid-based MPRA library, we cloned a 350-bp MYBPC2 promoter fragment (annotated by ENCODE, hg38, chr19:50432668-50433017) into the barcoded oligo-containing assembly (3:1 molar ratio of promoter insert:backbone) using BsaI-mediated Golden Gate assembly. We incubated the assembly reaction with the following program: [37*^◦^*C x 5 min, 16*^◦^*C x 5 min] x 30, 60*^◦^*C x 5 min. We performed a secondary digest with 1 U AsiSI to remove promoter-less assemblies, then cleaned with 0.8X SPRI beads and eluted in 10 uL of LoTE. We transformed as above, but expanded only 10% of the transformant pool in 150 mL of ampicillin-containing LB to bottleneck to ∼106 unique barcodes.

We performed a final restriction cloning step to move the assembled MPRA block (barcoded oligo, promoter, GFP) to the lentiviral transfer plasmid. We separately incubated the plasmid-based MPRA library and the lentiviral transfer backbone with EcoRI and SbfI for 1 hour, then gel purified our fragments of interest. We incubated the insert and backbone (3:1 molar ratio) with T4 DNA ligase for 10 minutes at room temperature and SPRI cleaned assemblies with 0.8X beads. Transduction-ready lentivirus was created by the University of Michigan Viral Vector Core.

#### MPRA experiment

We maintained LHCN-M2 human skeletal muscle myoblasts on 0.1% porcine gelatin coated dishes in manufacturer’s suggested medium (4:1 high glucose DMEM:Medium 199, 15% FBS, 20 mM HEPES, 3 ug/mL zinc sulfate, 1.4 ug/mL vitamin B12, 55 ng/mL dexamethasone, 2.5 ng/mL HGF, and 10 ng/mL bFGF).

Per replicate, we infected 4 × 106 cells with lentivirus at an MOI of ∼10 with 4 ug/mL polybrene. We passaged cells twice, then began differentiation 7 days after initial infection. We differentiated cells by performing daily media changes with differentiation media (DMEM 1 g/L glucose + 2% heat-inactivated horse serum). After 7 days of differentiation (i.e., 14 total days since infection), we lysed cells with 1.5 mL beta mercaptoethanol-containing Qiagen Buffer RLT Plus. We triturated the cell lysates with a syringe and 18-gauge needle 10 times to homogenize, then stored homogenized lysates at −80*^◦^*C until nucleic acid extraction.

#### MPRA sequencing

We used the Qiagen AllPrep RNA/DNA mini kit with four columns per replicate to isolate RNA and gDNA. We synthesized cDNA with 150 ug of DNAse-treated RNA using SuperScript IV reverse transcriptase and 100 nM custom GFP-targeted RT primer (jklab0363) containing a 15 bp UMI. We PCR amplified the cDNA with 500 nM primers jklab0268 and jklab0356 with NEB 2X Q5 HF HotStart PCR mastermix with the following program: 98*^◦^*C x 1 min, [98*^◦^*C × 10 sec, 60*^◦^*C x 30 sec, 72*^◦^*C x 1 min, plate read, 72*^◦^*C x 8 sec] x 20, 72*^◦^*C x 5:00. We added adapters and PCR amplified the gDNA samples using an analogous protocol. We performed sample indexing using standard Illumina P5 and P7 barcoding primers, then performed molar pooling and sequence samples on a NovaSeq 6000 (2 x 150 bp reads).

#### MPRA data analysis

To create the oligo-barcode pairing dictionary, we used a custom pipeline based on bwa v0.7.17 (3) to merge paired-end 150 bp reads. We extracted the barcodes. We then used minimap2 v2.24 (4) to align merged oligo reads against our reference FASTA file containing the expected oligo sequences. After filtering, we created a final table with oligo-barcode pairs and removed any duplicate barcodes.

For cDNA and gDNA barcode counting, we used cutadapt v4.3 (5) to trim sequencing adapters and constant sequences, UMI-tools v1.1.2 to cluster UMIs (6), and starcode v1.4 (7) to cluster and count deduplicated oligo barcodes. We merged these barcode counts with the pairing dictionary, requiring an exact match between the cDNA/gDNA barcode counts and the paired barcode. Finally, we calculated the sum of all barcode counts associated with a given oligo within a sample.

Prior to statistical modeling, we required a raw count mean ¿= 25 across all cDNA samples to remove any lowly expressed oligos. We estimated oligo activity and allelic bias using DESeq2 (8) with normalized read counts. We fit a nested fixed effects model as described previously (9).

To extract effects due to enhancer activity (RNA vs. DNA), we used linear contrasts between the cDNA and gDNA levels for a given replicate. To estimate allelic bias (reference vs. alternate allele), we used a linear contrast between the cDNA and gDNA counts for the reference and alternate alleles. We report the Benjamini-Hochberg FDR here to adjust for multiple testing.

## Notes

### Competing Interest Statement

Stephen CJ Parker has a research grant from Pfizer

### Summary of Updates

Added extensive analyses such as context-specific eQTL scans, multivariate adaptive shrinkage models on e,caQTL and new datasets.

## References

1. Frontera, W. R. & Ochala, J. Skeletal Muscle: A Brief Review of Structure and Function. en. Calcified Tissue International 96, 183–195 (Mar. 2015).

2. Kim, G. & Kim, J. H. Impact of Skeletal Muscle Mass on Metabolic Health. Endocrinology and Metabolism 35, 1–6 (Mar. 2020).

3. Morgan, J. & Partridge, T. Skeletal muscle in health and disease. Disease Models & Mechanisms 13, dmm042192 (Feb. 2020).

4. Sriwijitkamol, A. et al. Effect of Acute Exercise on AMPK Signaling in Skeletal Muscle of Subjects With Type 2 Diabetes: A Time-Course and Dose-Response Study. Diabetes 56, 836–848 (Mar. 2007).

5. Musi, N. et al. Metformin Increases AMP-Activated Protein Kinase Activity in Skeletal Muscle of Subjects With Type 2 Diabetes. Diabetes 51, 2074–2081 (July 2002).

6. Mahajan, A. et al. Multi-ancestry genetic study of type 2 diabetes highlights the power of diverse populations for discovery and translation. en. Nature Genetics. Publisher: Nature Publishing Group, 1–13 (May 2022).

7. Mahajan, A. et al. Fine-mapping type 2 diabetes loci to single-variant resolution using high-density imputation and islet-specific epigenome maps. En. Nature Genetics 50, 1505 (Nov. 2018).

8. Chen, J. et al. The trans-ancestral genomic architecture of glycemic traits. en. Nature Genetics 53. Number: 6 Publisher: Nature Publishing Group, 840–860 (June 2021).

9. Spracklen, C. N. et al. Identification of type 2 diabetes loci in 433,540 East Asian individuals. eng. Nature 582, 240–245 (June 2020).

10. Yengo, L. et al. Meta-analysis of genome-wide association studies for height and body mass index in 700000 individuals of European ancestry. Human Molecular Genetics 27, 3641–3649 (Oct. 2018).

11. Pulit, S. L. et al. Meta-analysis of genome-wide association studies for body fat distribution in 694 649 individuals of European ancestry. Human Molecular Genetics 28, 166–174 (Jan. 2019).

12. Maurano, M. T. et al. Systematic Localization of Common Disease-Associated Variation in Regulatory DNA. Science (New York, N.Y.) 337, 1190–1195 (Sept. 2012).

13. Parker, S. C. J. et al. Chromatin stretch enhancer states drive cell-specific gene regulation and harbor human disease risk variants. en. Proceedings of the National Academy of Sciences 110, 17921–17926 (Oct. 2013).

14. Quang, D. X., Erdos, M. R., Parker, S. C. J. & Collins, F. S. Motif signatures in stretch enhancers are enriched for disease-associated genetic variants. Epigenetics & Chromatin 8, 23 (July 2015).

15. Thurner, M. et al. Integration of human pancreatic islet genomic data refines regulatory mechanisms at Type 2 Diabetes susceptibility loci. en. eLife 7, e31977 (Feb. 2018).

16. Hormozdiari, F. et al. Colocalization of GWAS and eQTL Signals Detects Target Genes. The American Journal of Human Genetics 99, 1245–1260 (Dec. 2016).

17. Civelek, M. & Lusis, A. J. Systems genetics approaches to understand complex traits. eng. Nature Reviews. Genetics 15, 34–48 (Jan. 2014).

18. Viñuela, A., et al. Genetic variant effects on gene expression in human pancreatic islets and their implications for T2D. en. Nature Communications 11. Number: 1 Publisher: Nature Publishing Group, 4912 (Sept. 2020).

19. Wallace, C. A more accurate method for colocalisation analysis allowing for multiple causal variants. en. PLOS Genetics 17. Publisher: Public Library of Science, e1009440 (Sept. 2021).

20. Khetan, S. et al. Type 2 Diabetes–Associated Genetic Variants Regulate Chromatin Accessibility in Human Islets. en. Diabetes 67, 2466–2477 (Nov. 2018).

21. Currin, K. W. et al. Genetic effects on liver chromatin accessibility identify disease regulatory variants. en. The American Journal of Human Genetics (May 2021).

22. Robertson, C. C. et al. Fine-mapping, trans-ancestral and genomic analyses identify causal variants, cells, genes and drug targets for type 1 diabetes. en. Nature Genetics 53, 962–971 (July 2021).

23. Barbeira, A. N. et al. Exploiting the GTEx resources to decipher the mechanisms at GWAS loci. Genome Biology 22, 49 (Jan. 2021).

24. Liang, D. et al. Cell-type-specific effects of genetic variation on chromatin accessibility during human neuronal differentiation. en. Nature Neuroscience. Publisher: Nature Publishing Group, 1–13 (May 2021).

25. Aygün, N., et al. Brain-trait-associated variants impact cell-type-specific gene regulation during neurogenesis. eng. American Journal of Human Genetics 108, 1647–1668 (Sept. 2021).

26. Aygün, N., et al. Inferring cell-type-specific causal gene regulatory networks during human neurogenesis. eng. Genome Biology 24, 130 (May 2023).

27. Soskic, B. et al. Immune disease risk variants regulate gene expression dynamics during CD4+ T cell activation en. Tech. rep. Section: New Results Type: article (bioRxiv, Dec. 2021), 2021.12.06.470953.

28. Turner, A. W. et al. Single-nucleus chromatin accessibility profiling highlights regulatory mechanisms of coronary artery disease risk. en. Nature Genetics. Publisher: Nature Publishing Group, 1–13 (May 2022).

29. Scott, L. J. et al. The genetic regulatory signature of type 2 diabetes in human skeletal muscle. en. Nature Communications 7, ncomms11764 (June 2016).

30. Taylor, D. L. et al. Integrative analysis of gene expression, DNA methylation, physiological traits, and genetic variation in human skeletal muscle. en. Proceedings of the National Academy of Sciences 116. Publisher: National Academy of Sciences Section: Biological Sciences, 10883–10888 (May 2019).

31. Consortium, T. G. The GTEx Consortium atlas of genetic regulatory effects across human tissues. en. Science 369. Publisher: American Association for the Advancement of Science Section: Research Article, 1318–1330 (Sept. 2020).

32. Biferali, B., Proietti, D., Mozzetta, C. & Madaro, L. Fibro–Adipogenic Progenitors Cross-Talk in Skeletal Muscle: The Social Network. Frontiers in Physiology 10 (2019).

33. Contreras, O., Rossi, F. M. V. & Theret, M. Origins, potency, and heterogeneity of skeletal muscle fibro-adipogenic progenitors—time for new definitions. Skeletal Muscle 11, 16 (July 2021).

34. Molina, T., Fabre, P. & Dumont, N. A. Fibro-adipogenic progenitors in skeletal muscle homeostasis, regeneration and diseases. eng. Open Biology 11, 210110 (Dec. 2021).

35. Dumont, N. A., Bentzinger, C. F., Sincennes, M.-C. & Rudnicki, M. A. Satellite Cells and Skeletal Muscle Regeneration. eng. Comprehensive Physiology 5, 1027–1059 (July 2015).

36. Travers, M. E. et al. Insights into the molecular mechanism for type 2 diabetes susceptibility at the KCNQ1 locus from temporal changes in imprinting status in human islets. eng. Diabetes 62, 987–992 (Mar. 2013).

37. Hilton, T. N., Tuttle, L. J., Bohnert, K. L., Mueller, M. J. & Sinacore, D. R. Excessive Adipose Tissue Infiltration in Skeletal Muscle in Individuals With Obesity, Diabetes Mellitus, and Peripheral Neuropathy: Association With Performance and Function. Physical Therapy 88, 1336–1344 (Nov. 2008).

38. Teng, S. & Huang, P. The effect of type 2 diabetes mellitus and obesity on muscle progenitor cell function. Stem Cell Research & Therapy 10, 103 (Mar. 2019).

39. Kolka, C. M. & Bergman, R. N. The Endothelium in Diabetes: its role in insulin access and diabetic complications. Reviews in endocrine & metabolic disorders 14, 13–19 (Mar. 2013).

40. Pillon, N. J., Bilan, P. J., Fink, L. N. & Klip, A. Cross-talk between skeletal muscle and immune cells: muscle-derived mediators and metabolic implications. eng. American Journal of Physiology. Endocrinology and Metabolism 304, E453–465 (Mar. 2013).

41. Orchard, P. et al. Human and rat skeletal muscle single-nuclei multi-omic integrative analyses nominate causal cell types, regulatory elements, and SNPs for complex traits. en. Genome Research 31, 2258–2275 (Dec. 2021).

42. Zhang, K. et al. A single-cell atlas of chromatin accessibility in the human genome. English. Cell 184. Publisher: Elsevier, 5985–6001.e19 (Nov. 2021).

43. De Micheli, A. J., Spector, J. A., Elemento, O. & Cosgrove, B. D. A reference single-cell transcriptomic atlas of human skeletal muscle tissue reveals bifurcated muscle stem cell populations. Skeletal Muscle 10, 19 (July 2020).

44. Rubenstein, A. B. et al. Single-cell transcriptional profiles in human skeletal muscle. en. Scientific Reports 10. Number: 1 Publisher: Nature Publishing Group, 229 (Jan. 2020).

45. Welch, J. D. et al. Single-Cell Multi-omic Integration Compares and Contrasts Features of Brain Cell Identity. Cell 177, 1873–1887.e17 (June 2019).

46. Gao, C. et al. Iterative single-cell multi-omic integration using online learning. Nature biotechnology 39, 1000–1007 (Aug. 2021).

47. Duren, Z. et al. Regulatory analysis of single cell multiome gene expression and chromatin accessibility data with scREG. Genome Biology 23, 114 (May 2022).

48. Xiong, X. et al. Epigenomic dissection of Alzheimer’s disease pinpoints causal variants and reveals epigenome erosion. English. Cell 186. Publisher: Elsevier, 4422–4437.e21 (Sept. 2023).

49. Pliner, H. A. et al. Cicero Predicts cis-Regulatory DNA Interactions from Single-Cell Chromatin Accessibility Data. English. Molecular Cell 71, 858–871.e8 (Sept. 2018).

50. Liu, N. et al. Requirement of MEF2A, C, and D for skeletal muscle regeneration. Proceedings of the National Academy of Sciences 111. Publisher: Proceedings of the National Academy of Sciences, 4109–4114 (Mar. 2014).

51. Anderson, C. M. et al. Myocyte enhancer factor 2C function in skeletal muscle is required for normal growth and glucose metabolism in mice. Skeletal Muscle 5, 7 (Feb. 2015).

52. Goveia, J. et al. Endothelial Cell Differentiation by SOX17. Circulation Research 115. Publisher: American Heart Association, 205–207 (July 2014).

53. Liu, M. et al. Sox17 is required for endothelial regeneration following inflammation-induced vascular injury. en. Nature Communications 10. Number: 1 Publisher: Nature Publishing Group, 2126 (May 2019).

54. Yao, Y., Yao, J. & Boström, K. I. SOX Transcription Factors in Endothelial Differentiation and Endothelial-Mesenchymal Transitions. Frontiers in Cardiovascular Medicine 6, 30 (Mar. 2019).

55. Von Maltzahn, J., Jones, A. E., Parks, R. J. & Rudnicki, M. A. Pax7 is critical for the normal function of satellite cells in adult skeletal muscle. Proceedings of the National Academy of Sciences 110. Publisher: Proceedings of the National Academy of Sciences, 16474–16479 (Oct. 2013).

56. Wang, Y. et al. Myeloid cell-specific mutation of Spi1 selectively reduces M2-biased macrophage numbers in skeletal muscle, reduces age-related muscle fibrosis and prevents sarcopenia. Aging Cell 21, e13690 (Oct. 2022).

57. Jimenez, M. A., Akerblad, P., Sigvardsson, M. & Rosen, E. D. Critical role for Ebf1 and Ebf2 in the adipogenic transcriptional cascade. eng. Molecular and Cellular Biology 27, 743–757 (Jan. 2007).

58. Kanki, Y. et al. Epigenetically coordinated GATA2 binding is necessary for endothelium-specific endomucin expression. eng. The EMBO journal 30, 2582–2595 (June 2011).

59. Kim, D. W. et al. Gene regulatory networks controlling differentiation, survival, and diversification of hypothalamic Lhx6-expressing GABAergic neurons. en. Communications Biology 4. Publisher: Nature Publishing Group, 1–16 (Jan. 2021).

60. Li, S. et al. Endothelial cell-derived GABA signaling modulates neuronal migration and postnatal behavior. Cell Research 28, 221–248 (Feb. 2018).

61. Varshney, A. et al. A Transcription Start Site Map in Human Pancreatic Islets Reveals Functional Regulatory Signatures. en. Diabetes 70, 1581–1591 (July 2021).

62. Varshney, A. et al. Cell Specificity of Human Regulatory Annotations and Their Genetic Effects on Gene Expression. en. Genetics 211, 549–562 (Feb. 2019).

63. Urbut, S. M., Wang, G., Carbonetto, P. & Stephens, M. Flexible statistical methods for estimating and testing effects in genomic studies with multiple conditions. En. Nature Genetics 51, 187 (Jan. 2019).

64. Cuomo, A. S. E. et al. CellRegMap: a statistical framework for mapping context-specific regulatory variants using scRNA-seq. Molecular Systems Biology 18. Publisher: John Wiley & Sons, Ltd, e10663 (Aug. 2022).

65. Wang, J. et al. CAUSALdb: a database for disease/trait causal variants identified using summary statistics of genome-wide association studies. en. Nucleic Acids Research 48. Publisher: Oxford Academic, D807–D816 (Jan. 2020).

66. Riazi, A. M., Lee, H., Hsu, C. & Van Arsdell, G. CSX/Nkx2.5 modulates differentiation of skeletal myoblasts and promotes differentiation into neuronal cells in vitro. eng. The Journal of Biological Chemistry 280, 10716–10720 (Mar. 2005).

67. Millstein, J., Zhang, B., Zhu, J. & Schadt, E. E. Disentangling molecular relationships with a causal inference test. BMC Genetics 10, 23 (May 2009).

68. Millstein, J., Chen, G. K. & Breton, C. V. cit: hypothesis testing software for mediation analysis in genomic applications. Bioinformatics 32, 2364–2365 (Aug. 2016).

69. Hemani, G., Tilling, K. & Smith, G. D. Orienting the causal relationship between imprecisely measured traits using GWAS summary data. PLOS Genetics 13, e1007081 (Nov. 2017).

70. Bulik-Sullivan, B. K. et al. LD Score regression distinguishes confounding from polygenicity in genome-wide association studies. en. Nature Genetics 47, 291–295 (Mar. 2015).

71. Finucane, H. K. et al. Partitioning heritability by functional annotation using genome-wide association summary statistics. en. Nature Genetics 47, 1228–1235 (Nov. 2015).

72. Cano-Gamez, E. & Trynka, G. From GWAS to Function: Using Functional Genomics to Identify the Mechanisms Underlying Complex Diseases. eng. Frontiers in Genetics 11, 424 (2020).

73. Zhang, Z., Feng, F., Qiu, Y. & Liu, J. A generalizable framework to comprehensively predict epigenome, chromatin organization, and transcriptome. eng. Nucleic Acids Research 51, 5931–5947 (July 2023).

74. Singh, R. et al. Follistatin Targets Distinct Pathways To Promote Brown Adipocyte Characteristics in Brown and White Adipose Tissues. Endocrinology 158, 1217–1230 (Jan. 2017).

75. Singh, R. et al. Metabolic profiling of follistatin overexpression: a novel therapeutic strategy for metabolic diseases. Diabetes, Metabolic Syndrome and Obesity: Targets and Therapy 11, 65–84 (Mar. 2018).

76. Zheng, H. et al. Follistatin N terminus differentially regulates muscle size and fat in vivo. en. Experimental & Molecular Medicine 49. Number: 9 Publisher: Nature Publishing Group, e377–e377 (Sept. 2017).

77. Braga, M. et al. Follistatin promotes adipocyte differentiation, browning, and energy metabolism. eng. Journal of Lipid Research 55, 375–384 (Mar. 2014).

78. Williamson, A. et al. Genome-wide association study and functional characterization identifies candidate genes for insulin-stimulated glucose uptake. en. Nature Genetics 55. Number: 6 Publisher: Nature Publishing Group, 973–983 (June 2023).

79. Mueckler, M. Insulin resistance and the disruption of Glut4 trafficking in skeletal muscle. Journal of Clinical Investigation 107, 1211–1213 (May 2001).

80. Kycia, I. et al. A Common Type 2 Diabetes Risk Variant Potentiates Activity of an Evolutionarily Conserved Islet Stretch Enhancer and Increases C2CD4A and C2CD4B Expression. The American Journal of Human Genetics 102, 620–635 (Apr. 2018).

81. Mehta, Z. B. et al. Changes in the expression of the type 2 diabetes-associated gene VPS13C in the -cell are associated with glucose intolerance in humans and mice. American Journal of Physiology - Endocrinology and Metabolism 311, E488–E507 (Aug. 2016).

82. Umans, B. D., Battle, A. & Gilad, Y. Where are the disease-associated eQTLs? Trends in genetics : TIG 37, 109–124 (Feb. 2021).

83. Mostafavi, H., Spence, J. P., Naqvi, S. & Pritchard, J. K. Limited overlap of eQTLs and GWAS hits due to systematic differences in discovery en. Pages: 2022.05.07.491045 Section: New Results. May 2022.

84. Varshney, A. et al. Genetic regulatory signatures underlying islet gene expression and type 2 diabetes. en. Proceedings of the National Academy of Sciences 114, 2301–2306 (Feb. 2017).

85. Alasoo, K. et al. Shared genetic effects on chromatin and gene expression indicate a role for enhancer priming in immune response. En. Nature Genetics, 1 (Jan. 2018).

86. Aracena, K. A. et al. Epigenetic variation impacts individual differences in the transcriptional response to influenza infection. en. Nature Genetics 56. Publisher: Nature Publishing Group, 408–419 (Mar. 2024).

87. Matoba, N., et al. Wnt activity reveals context-specific genetic effects on gene regulation in neural progenitors en. Pages: 2023.02.07.527357 Section: New Results. Apr. 2023.

88. Sun, N. et al. Human microglial state dynamics in Alzheimer’s disease progression. English. Cell 186. Publisher: Elsevier, 4386–4403.e29 (Sept. 2023).

89. Wijst, M. G. P. v. d., et al. Single-cell RNA sequencing identifies celltype-specific cis-eQTLs and co-expression QTLs. en. Nature Genetics 50, 493–497 (Apr. 2018).

90. Yazar, S. et al. Single-cell eQTL mapping identifies cell type–specific genetic control of autoimmune disease. Science 376. Publisher: American Association for the Advancement of Science, eabf3041.

91. Neavin, D. et al. Single cell eQTL analysis identifies cell type-specific genetic control of gene expression in fibroblasts and reprogrammed induced pluripotent stem cells. Genome Biology 22, 76 (Mar. 2021).

92. Cuomo, A. S. E. et al. Single-cell RNA-sequencing of differentiating iPS cells reveals dynamic genetic effects on gene expression. en. Nature Communications 11. Number: 1 Publisher: Nature Publishing Group, 810 (Feb. 2020).

93. Bryois, J. et al. Genetic identification of cell types underlying brain complex traits yields insights into the etiology of Parkinson’s disease. en. Nature Genetics. Publisher: Nature Publishing Group, 1–12 (Apr. 2020).

94. Jerber, J. et al. Population-scale single-cell RNA-seq profiling across dopaminergic neuron differentiation. en. bioRxiv. Publisher: Cold Spring Harbor Laboratory Section: New Results, 2020.05.21.103820 (May 2020).

95. Benaglio, P. et al. Mapping genetic effects on cell type-specific chromatin accessibility and annotating complex immune trait variants using single nucleus ATAC-seq in peripheral blood. PLOS Genetics 19, e1010759 (June 2023).

96. Ahlmann-Eltze, C. & Huber, W. Analysis of multi-condition single-cell data with latent embedding multivariate regression en. Pages: 2023.03.06.531268 Section: New Results. Mar. 2023.

97. Attaf, N. et al. FB5P-seq: FACS-Based 5-Prime End Single-Cell RNA-seq for Integrative Analysis of Transcriptome and Antigen Receptor Repertoire in B and T Cells. English. Frontiers in Immunology 11. Publisher: Frontiers (Mar. 2020).

98. Katzenelenbogen, Y. et al. Coupled scRNA-Seq and Intracellular Protein Activity Reveal an Immunosuppressive Role of TREM2 in Cancer. Cell 182, 872–885.e19 (Aug. 2020).

99. Duvall, E. et al. Single-cell transcriptome and accessible chromatin dynamics during endocrine pancreas development. Proceedings of the National Academy of Sciences of the United States of America 119, e2201267119 (June 2022).

100. Lal, A. et al. Deep learning-based enhancement of epigenomics data with AtacWorks. en. Nature Communications 12. Number: 1 Publisher: Nature Publishing Group, 1507 (Mar. 2021).

101. Rai, V. et al. Single cell ATAC-seq in human pancreatic islets and deep learning upscaling of rare cells reveals cell-specific type 2 diabetes regulatory signatures. en. Molecular Metabolism (Dec. 2019).

102. Tewhey, R. et al. Direct Identification of Hundreds of Expression-Modulating Variants using a Multiplexed Reporter Assay. eng. Cell 165, 1519–1529 (June 2016).

103. Abell, N. S. et al. Multiple Causal Variants Underlie Genetic Associations in Humans. Science (New York, N.Y.) 375, 1247–1254 (Mar. 2022).

104. Siraj, L., et al. Functional dissection of complex and molecular trait variants at single nucleotide resolution en. Pages: 2024.05.05.592437 Section: New Results. May 2024.

105. Das, S. et al. Next-generation genotype imputation service and methods. eng. Nature Genetics 48, 1284–1287 (2016).

106. Loh, P.-R. et al. Reference-based phasing using the Haplotype Reference Consortium panel. en. Nature Genetics 48. Number: 11 Publisher: Nature Publishing Group, 1443–1448 (Nov. 2016).

107. Kang, H. M. et al. Multiplexed droplet single-cell RNA-sequencing using natural genetic variation. en. Nature Biotechnology 36, 89–94 (Jan. 2018).

108. Lun, A. T. L. et al. EmptyDrops: distinguishing cells from empty droplets in droplet-based single-cell RNA sequencing data. en. Genome Biology 20. Number: 1 Publisher: BioMed Central, 1–9 (Dec. 2019).

109. Yang, S. et al. Decontamination of ambient RNA in single-cell RNA-seq with DecontX. en. Genome Biology 21. Number: 1 Publisher: BioMed Central, 1–15 (Dec. 2020).

110. Li, H. & Durbin, R. Fast and accurate short read alignment with Burrows-Wheeler transform. eng. Bioinformatics (Oxford, England) 25, 1754–1760 (July 2009).

111. Zhang, Y. et al. Model-based Analysis of ChIP-Seq (MACS). Genome Biology 9, R137 (Sept. 2008).

112. Quinlan, A. R. & Hall, I. M. BEDTools: a flexible suite of utilities for comparing genomic features. en. Bioinformatics 26, 841–842 (Mar. 2010).

113. Amemiya, H. M., Kundaje, A. & Boyle, A. P. The ENCODE Blacklist: Identification of Problematic Regions of the Genome. en. Scientific Reports 9. Number: 1 Publisher: Nature Publishing Group, 9354 (June 2019).

114. Beshnova, D. A., Cherstvy, A. G., Vainshtein, Y. & Teif, V. B. Regulation of the Nucleosome Repeat Length In Vivo by the DNA Sequence, Protein Concentrations and Long-Range Interactions. PLoS Computational Biology 10, e1003698 (July 2014).

115. Kent, W. J., Zweig, A. S., Barber, G., Hinrichs, A. S. & Karolchik, D. BigWig and BigBed: enabling browsing of large distributed datasets. Bioinformatics 26, 2204–2207 (Sept. 2010).

116. Love, M. I., Huber, W. & Anders, S. Moderated estimation of fold change and dispersion for RNA-seq data with DESeq2. Genome Biology 15, 550 (Dec. 2014).

117. Frankish, A. et al. GENCODE 2021. Nucleic Acids Research 49, D916–D923 (Jan. 2021).

118. Raudvere, U. et al. g:Profiler: a web server for functional enrichment analysis and conversions of gene lists (2019 update). Nucleic Acids Research 47, W191–W198 (July 2019).

119. Yanai, I. et al. Genome-wide midrange transcription profiles reveal expression level relationships in human tissue specification. eng. Bioinformatics (Oxford, England) 21, 650–659 (Mar. 2005).

120. D’Oliveira Albanus, R., et al. Chromatin information content landscapes inform transcription factor and DNA interactions. en. Nature Communications 12. Number: 1 Publisher: Nature Publishing Group, 1307 (Feb. 2021).

121. Grant, C. E., Bailey, T. L. & Noble, W. S. FIMO: scanning for occurrences of a given motif. en. Bioinformatics 27, 1017–1018 (Apr. 2011).

122. Delaneau, O. et al. A complete tool set for molecular QTL discovery and analysis. En. Nature Communications 8, 15452 (May 2017).

123. Mohammadi, P., Castel, S. E., Brown, A. A. & Lappalainen, T. Quantifying the regulatory effect size of cis-acting genetic variation using allelic fold change. en. Genome Research (Oct. 2017).

124. Taylor-Weiner, A. et al. Scaling computational genomics to millions of individuals with GPUs. en. Genome Biology 20. Number: 1 Publisher: BioMed Central, 1–5 (Dec. 2019).

125. Wang, G., Sarkar, A., Carbonetto, P. & Stephens, M. A simple new approach to variable selection in regression, with application to genetic fine mapping. en. Journal of the Royal Statistical Society: Series B (Statistical Methodology) 82, 1273–1300 (Dec. 2020).

126. The Roadmap Epigenomics Consortium et al. Integrative analysis of 111 reference human epigenomes. en. Nature 518, 317–330 (Feb. 2015).

127. Hinrichs, A. S. et al. The UCSC Genome Browser Database: update 2006. eng. Nucleic Acids Research 34, D590–598 (Jan. 2006).

128. Finucane, H. K. et al. Heritability enrichment of specifically expressed genes identifies disease-relevant tissues and cell types. en. Nature Genetics 50, 621–629 (Apr. 2018).

129. Sudlow, C. et al. UK Biobank: An Open Access Resource for Identifying the Causes of a Wide Range of Complex Diseases of Middle and Old Age. en. PLOS Medicine 12. Publisher: Public Library of Science, e1001779 (Mar. 2015).

130. Tashman, K. C., Cui, R., O’Connor, L. J., Neale, B. M. & Finucane, H. K. Significance testing for small annotations in stratified LD-Score regression en. ISSN: 2124-9938 Pages: 2021.03.13.21249938. Mar. 2021.

131. Pickrell, J. Joint Analysis of Functional Genomic Data and Genome-wide Association Studies of 18 Human Traits. American Journal of Human Genetics 94, 559–573 (Apr. 2014).

132. Vanhille, L. et al. High-throughput and quantitative assessment of enhancer activity in mammals by CapStarr-seq. Nature Communications 6, 6905 (Apr. 2015).

133. Melnikov, A., Zhang, X., Rogov, P., Wang, L. & Mikkelsen, T. S. Massively Parallel Reporter Assays in Cultured Mammalian Cells. Journal of Visualized Experiments : JoVE, 51719 (Aug. 2014).

134. Hao, Y., et al. Dictionary learning for integrative, multimodal, and scalable single-cell analysis en. Pages: 2022.02.24.481684 Section: New Results. Feb. 2022.

